# Binding domain mutations provide insight into CTCF’s relationship with chromatin and its contribution to gene regulation

**DOI:** 10.1101/2024.01.11.575070

**Authors:** Catherine Do, Guimei Jiang, Giulia Cova, Christos C. Katsifis, Domenic N. Narducci, Theodore Sakellaropoulos, Raphael Vidal, Priscillia Lhoumaud, Aristotelis Tsirigos, Faye Fara D. Regis, Nata Kakabadze, Elphege P Nora, Marcus Noyes, Anders S. Hansen, Jane A Skok

**Author notes:** These authors contributed equally (Catherine Do, Guimei Jiang, Giulia Cova, Christos C. Katsifis, Domenic N. Narducci).

## Abstract

Here we used a series of CTCF mutations to explore CTCF’s relationship with chromatin and its contribution to gene regulation. CTCF’s impact depends on the genomic context of bound sites and the unique binding properties of WT and mutant CTCF proteins. Specifically, CTCF’s signal strength is linked to changes in accessibility, and the ability to block cohesin is linked to its binding stability. Multivariate modelling reveals that both CTCF and accessibility contribute independently to cohesin binding and insulation, however CTCF signal strength has a stronger effect. CTCF and chromatin have a bidirectional relationship such that at CTCF sites, accessibility is reduced in a cohesin-dependent, mutant specific fashion. In addition, each mutant alters TF binding and accessibility in an indirect manner, changes which impart the most influence on rewiring transcriptional networks and the cell’s ability to differentiate. Collectively, the mutant perturbations provide a rich resource for determining CTCF’s site-specific effects.

## Introduction

The CCCTC-binding factor (CTCF) is an eleven-zinc finger DNA-binding protein that together with its binding partner, cohesin plays a key role in organizing chromatin into TAD structures by promoting the formation of loops and boundaries that are important for gene regulation. This involves a loop-extrusion mechanism in which cohesin complexes create chromatin loops by actively extruding DNA until movement of the complex is blocked by two CTCF binding sites in convergent orientation^1,2^. CTCF, in conjunction with cohesin, is enriched at TAD boundaries that function as insulators, contributing to gene regulation by restricting the interaction of regulatory elements to promoters of target genes located within the same TAD^3,4^.

CTCF has been degraded by auxin in numerous cell types, and although these studies provide important insight into its role in gene regulation, degron models are insufficient for (i) analyzing the contribution of genomic context to site-specific CTCF binding and function, or (ii) distinguishing direct from indirect effects that can be a confounding issue for interpreting site specific impact. Instead, information related to CTCF’s direct effects on individual loci has come from the genetic ablation of CTCF binding sites at specific regions in the genome in a given cell type. However, interrogating the general mechanisms underlying the crosstalk between CTCF function and the cell- and locus-specific context of its binding by genetic manipulation of individual CTCF binding sites is laborious and has the limitation of providing insight into the control of only a limited number of loci in the neighboring region.

The *CTCF* locus, located on chromosome 16q band 22, corresponds to one of the smallest regions of overlap for common deletions in breast and prostate cancers^5,6^. Moreover, point mutations and deletions have been identified in many other tumors. Together these findings indicate that CTCF acts as a tumor suppressor^7^. Additionally, genetic alterations in *CTCF* and changes in dosage are associated to varying degrees with intellectual disability and microcephaly, and thus CTCF is thought to play an important role in brain development and neurological disorders^8–10^.

CTCF binds to chromatin through a subset of its eleven zinc fingers (ZFs)^11,12^. Specifically, ZFs3- 7 make sequence specific contacts with a 12-15 base-pair (bp) core consensus binding site. While the crystal structure of CTCF in complex with a known DNA binding site^13^ has revealed new insights into the contribution of each ZF to sequence-specific binding in an *in vitro* setting, there has been no complementary, comprehensive analysis of cancer and neurological development disorders (NDDs) associated CTCF mutants and it is thus not known how mutations within each ZF impact binding stability and DNA sequence specificity, and the consequences this has on binding profile, cohesin overlap, accessibility, chromosome architecture and gene regulation. Given this lack of information, we have no context in which to determine the functional relevance of mutations.

Using an inducible mouse ESC complementation system, we combined genetic, imaging, molecular, AlphaFold3 and bioinformatic analyses to examine the impact of nine, high frequency cancer associated CTCF mutations, a subset of which are also associated with NDDs. Selected mutations occur in eight amino acid residues of ZFs that contact the core consensus binding motif. Compared to a complete knockout of CTCF or deletion of a CTCF binding site, the mutants offer a more subtle and in-depth approach for teasing out (i) the relationship between WT and mutant CTCF and chromatin, and (ii) direct or indirect site-specific effects on accessibility, TF binding, cohesin overlap, chromatin interactivity and gene expression, since each mutation produces its own unique effect.

Here we demonstrate that the functional impact of WT and mutant CTCF depends both on the genomic context of bound sites and the unique binding properties of each WT and mutant CTCF protein. Analyses of the relationship between CTCF, cohesin and accessibility reveals that although CTCF can bind both accessible and inaccessible sites, inaccessible bound sites predominantly have weaker signals and are less competent at blocking cohesin. Multivariate modelling demonstrates that both CTCF and accessibility contribute independently to cohesin binding, however CTCF signal strength has a dominant effect, linked to a stronger insulation score. Indeed, CTCFs ability to block cohesin is linked to the residence time or ‘OFF’ rate of each mutant on chromatin as determined by FRAP experiments, while occupancy or ‘ON’ rate is correlated with their binding at common accessible sites, indicating that accessibility dominates the search space. Together, the mutants highlight the fact that occupancy and residence time can be uncoupled such that although some mutants have fewer binding sites than WT CTCF, they have a stronger residence time at these locations which increases their ability to block cohesin and contribute to chromatin folding. The reverse situation also occurs. Our analyses also highlight that in contrast to CTCF-bound sites, cohesin binding at CTCF-free sites is weak and does not track with accessibility even at enhancers and promoters, perhaps because TFs that block cohesin have considerably shorter residence times compared to CTCF.

The unique properties of WT and mutant CTCF perturb TF binding and cancer, brain, immune and metabolic gene expression pathways. Mutants have a more pronounced effect on transcriptional networks than auxin inducible degradation, because of indirect effects that can be explained by altered TF binding profiles. Indeed, indirect effects that do not overlap changes in CTCF binding impart the most influence on rewiring chromatin looping and transcriptional networks. Changes in each mutant’s transcriptional program in turn have an impact on the cell’s ability to exit from pluripotency and differentiate. Collectively, the mutants offer a rich resource to investigate site specific CTCF-mediated effects on chromatin folding and gene regulation, while distinguishing cause from consequence and direct from indirect outcomes.

## RESULTS

### CTCF complementation system

To explore CTCF’s relationship with chromatin and its contribution to gene regulation, we made use of a small-molecule auxin inducible degron (AID) mouse ESC (mESC) system^14,15^. In these cells, both endogenous CTCF alleles are tagged with AID as well as eGFP (CTCF-AID-eGFP) and the auxin-activated ubiquitin ligase TIR1 (from *Oryza sativa*) is constitutively expressed from the *Rosa26* locus (**Figure 1A)**. Addition of indole acetic acid (IAA), an analogue of auxin leads to rapid poly-ubiquitination and proteasomal degradation of the AID tagged protein^16^.

**Figure 1:**
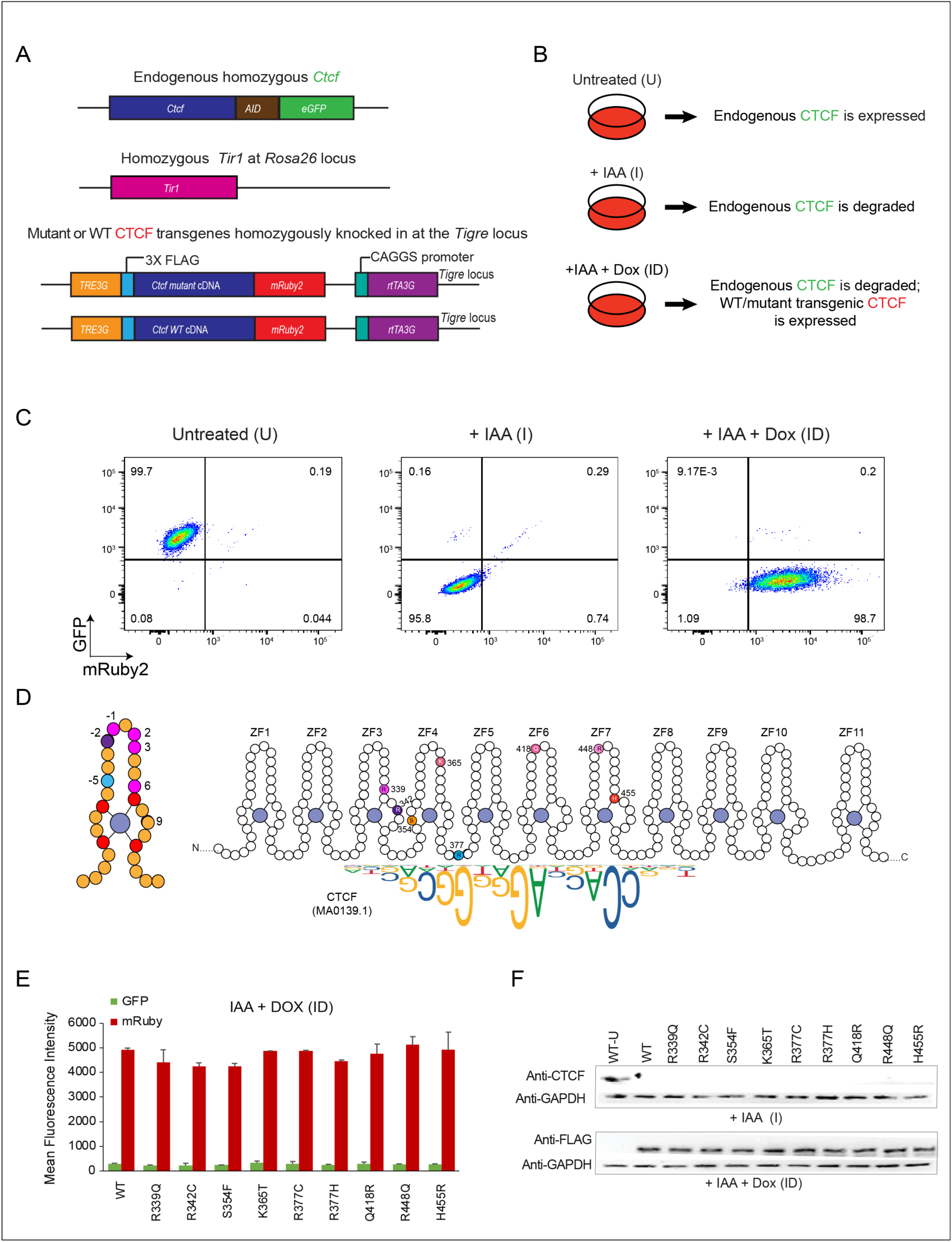
CTCF complementation system. (**A)** Scheme of the CTCF doxycycline-inducible degron system. (**B**) Experimental strategy for the expression of WT and mutant transgenic CTCF. (**C**) Flow cytometry showing the level of GFP (endogenous CTCF) and mRuby (transgenic WT CTCF). (**D**) Left, scheme showing the locations of the different types of CTCF mutations within a ZF. Amino acids making contacts with the DNA are shown in shades of pink, residues that coordinate the zinc ion in red, boundary residues in purple, and residues that contact the sugar phosphate backbone of DNA in blue. Right, representation of CTCF showing the locations of each mutation under investigation. (**E**) Bar graph showing the expression levels (mean fluorescence intensity) of endogenous CTCF (GFP) and transgenic WT or mutant CTCF (mRuby2) after ID treatment. The error bars represent the standard deviation between the two replicates. (**F**) Western blot of endogenous CTCF (CTCF antibody) and transgenic CTCF (FLAG antibody) in untreated (WT-U), IAA, ID conditions. See also **Figures S1-5**.

To study the impact of CTCF mutations, we established a rescue system, modifying the mESC degron cell line to express either a stable doxycycline-inducible control wild-type *Ctcf* or a mutant *Ctcf* (*mCtcf*) transgene in the absence of endogenous CTCF^17^. Transgenes, knocked into the *Tigre* locus, have a 3 x FLAG tag, which allows us to distinguish mutant or wild-type transgenic CTCF from endogenous CTCF using a FLAG antibody in Western blot and ChIP-seq analysis. An mRuby fluorescent tag enables us to accurately determine expression levels of mutant or wild-type transgenic CTCF by flow cytometry (**Figure 1A**). The three conditions used for our analysis are shown in **Figure 1B**, and an example of the FACS profile for cells expressing a WT CTCF transgene in each condition is shown (**Figure 1C**).

We selected a subset of missense mutations (identified from cBioPortal ^18^ and COSMIC^19^) that occur at high frequency in cancer patients (**Figure S1**), focusing on those found in ZFs that contact the 12-15bp consensus CTCF binding motif (**Figure 1D** and highlighted with an asterisk in **Figure S2**). We analyzed mutations in **(i)** amino acids that make base-specifying contacts with the alpha helix of DNA (at locations -2, 2, 3 and 6 on the ZF), **(ii)** residues that coordinate the zinc ion which are essential for providing stability to ensure the proper folding of the ZF domain, **(iii)** two other classes of amino acid residues in the ZFs: boundary residues (that are important for interactions between ZFs), and residues that contact the sugar phosphate backbone of DNA, and **(iv)** other highly mutated residues in the region. Each class of mutation is color-coded as shown in **Figure 1D** (left) and **Figures S1-2** and the color code is maintained throughout the figures.

In total, we analyzed mutations in eight amino acid residues and for R377, the most highly mutated residue in cancer, we included two distinct mutations of R377, both of which (R377H and R377C) are equally represented in patients (**Figure S2**). Selected mutations (R339Q, R342C, R448Q, R377H and R377C) are also implicated in NDDs (**Figure S2**)^10,20^. Individual clones with comparable levels of transgene expression (mutant and WT) were selected based on flow cytometry and Western blot analysis (**Figure 1E, F and Figures S3-4**). To assess rescue of endogenous CTCF by WT transgenic CTCF, we performed ChIP-seq using an anti-CTCF antibody and RNA-seq in WT untreated and WT transgene expressing (ID) cells. This analysis showed that about half of the overall CTCF binding sites (but 97% of the CTCF motif-containing sites) and gene expression changes are rescued by transgenic CTCF (**Figure S5**), consistent with the lower expression of the transgene reported previously^17^. This data provides a rationale for comparing the effect of the mutant and WT transgenic CTCF.

### Retained binding sites are enriched in accessible regions with strong CTCF signals

We next analyzed the profiles of mutant versus WT CTCF using cells expressing transgenic WT and mutant CTCF in the absence of endogenous CTCF (ID). Briefly, we degraded endogenous CTCF-mAID-GFP by IAA addition, induced WT or mutant transgene expression with Dox for 48 hours, and then performed FLAG ChIP-seq. For each mutation we identified WT only sites (bound by WT CTCF that mutants no longer bind), common sites (bound by both WT and mutant proteins) and mutant only sites (*de novo* sites that only mutant proteins bind).

The location of each mutation and the complementary DNA triplet that the relevant ZF binds is marked by a box in the consensus motif shown in **Figure 2A** and **Figure S6A**. Below that we highlight the most frequent motif identified by MEME^21^ for the set of *de novo* sites. These motifs differ from that of the consensus sequence by either (i) a diminished requirement for DNA bases that contact the mutated zinc finger (**Figure 2A**), or (ii) an alteration to the sequence of the consensus motif, which could be explained by the impact of mutations on DNA-base interacting zinc finger residues. For example, K365 binds to the second guanine in the triplet GGC of the consensus motif and the *de novo* motif found in the K365T mutant is consistent with loss of hydrogen bonds with the central guanine due to the threonine substitution, as previously demonstrated^13^. Motifs found at WT only and common sites were similar to the consensus motif (**Figure S6B**), with the exception of R377C for which few WT only sites were observed. On average, the percentages of CTCF containing motifs at common sites were higher than at WT only sites, suggesting that the latter might be lower affinity sites with degenerate motifs, consistent with the weaker CTCF signal observed. Each mutant displays their own unique binding profiles including the two distinct mutations of residue R377. R377C binds to less common and more *de novo* sites compared to R377H and each mutant binds a motif that is similar to the consensus motif but lacking bases associated with ZF3 (**Figure 2A** and **Figure S6**).

**Figure 2:**
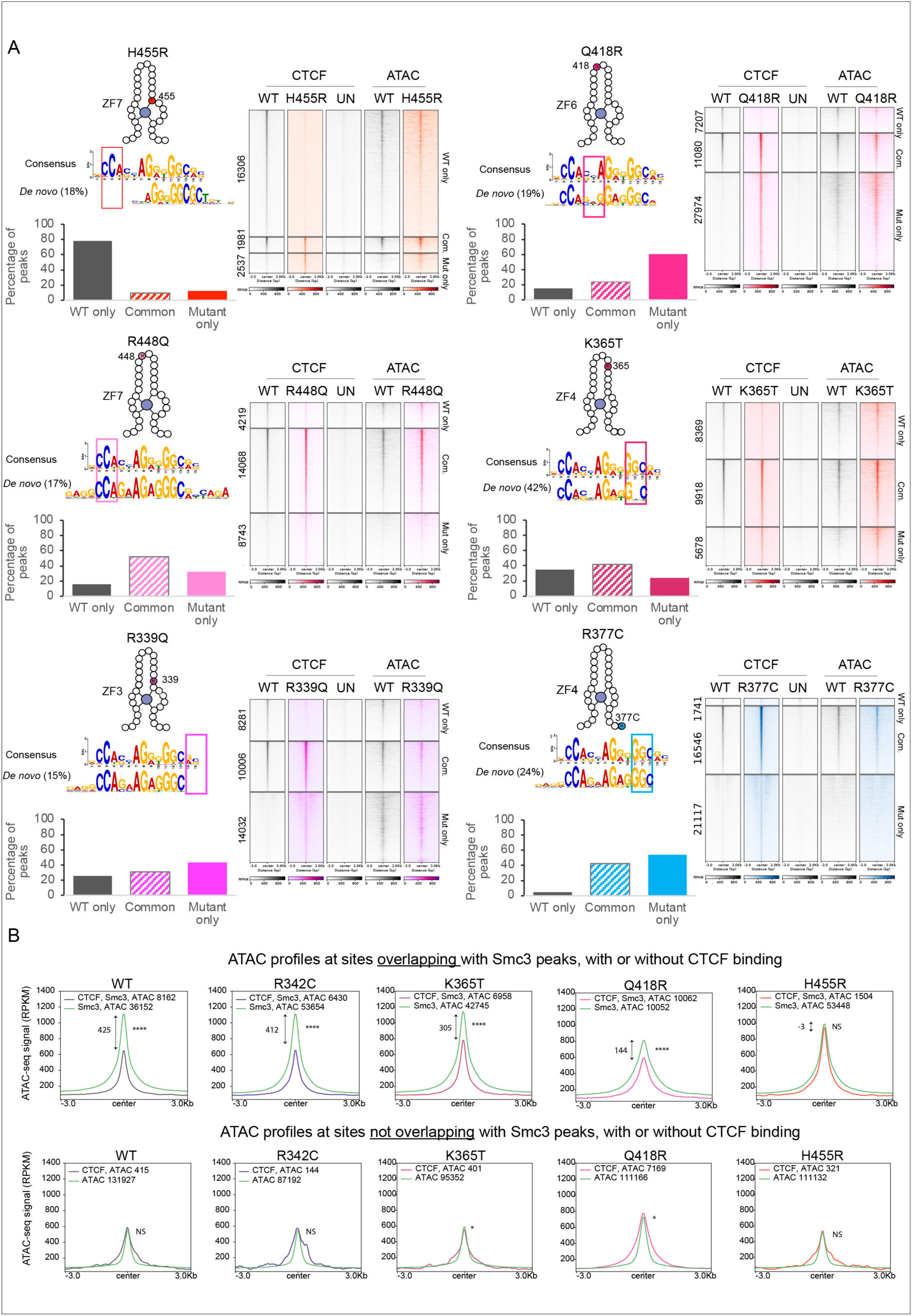
CTCF mutations have unique chromatin binding profiles. (**A)** Scheme showing the locations of the CTCF mutations within the ZF. The consensus CTCF motif highlights the triplet to which each mutant ZF binds (top). The most common motif for the *de novo* binding sites is shown below.. Each bar graph shows the percentage of WT only, common and mutant only CTCF binding sites. The heatmaps show the profile of CTCF and ATAC-seq signal at those sites. The UN condition corresponds to the FLAG control. (**B**) Profiles of ATAC-seq in WT and mutants. Wilcoxon p-values were coded as follow: NS, not significant, * 5×10^-2^-5×10^-3^, ** 5×10^-3^-5×10^-4^, *** 5×10^-4^-5×10^-5^, **** <5×10^-5^. Data were generated on 2 replicates. See also **Figures S6-9.**

The bar graphs show the percentage of WT only, common and mutant only binding sites, while the heatmaps reveal the number of binding sites for each subset and highlight differences in CTCF ChIP-seq signal between WT and mutants at the three locations. Differential binding analysis using DiffBind (**Figure S7)** showed that only a fraction of the mutant and WT only peaks reached statistical significance, which can partly be explained by the fact that these peaks are weaker and have a smaller effect size compared to the stronger, common peaks. However, the ratios of gained versus lost CTCF binding sites are consistent with the proportion of ‘WT only’ and ’mutant only’ peaks (**Figure 2A** and **S6A**). CTCF signal strength is strongest at common sites, however, CTCF this is altered in a mutant specific manner, suggesting that a ’binding dosage effect’ contributes to their functional impact. As expected, the H455R mutation in the zinc coordinating residue of ZF7 loses the most binding sites. OmniATAC^22^ profiles for the different binding groups indicate that CTCF ChIP-seq signal strength is a function of chromatin accessibility, with common sites being the most accessible (**Figure 2A and Figure S6A**).

In sum, ChIP-seq analysis reveals that CTCF mutants lose, retain and gain a subset of binding sites throughout the genome with mutant specific profiles. Loss and gain of CTCF binding occurs predominantly at weak inaccessible CTCF sites, while binding is retained at the stronger common binding sites which have high accessibility, although CTCF ChIP-seq signal and accessibility can vary depending on the mutant.

### CTCF reduces ATAC-seq signal in a cohesin-dependent manner

To further investigate the link between accessibility and CTCF binding, we compared the ATAC-seq signal at CTCF-bound versus unbound sites in the presence or absence of SMC3, a subunit of the cohesin complex (**Figure 2B** and **Figure S8**). Accessibility was reduced in an SMC3-dependent manner at both WT and mutant CTCF-bound sites. Each CTCF mutant exhibited a differential ability to decrease the height and breadth of ATAC-seq peaks, suggesting that CTCF is driving the change in accessibility and that the relationship between CTCF and accessibility is bidirectional. We do not observe the same phenomenon at sites without SMC3, which lends support to the idea that loop formation decreases accessibility, indicating a unique chromatin structure at loop bases. The same phenomenon is observed at enhancer and promoter sites, although the effect is weaker, particularly at promoters (**Figure S9**). The weakened effect may be due to interference of TF binding which can affect CTCF-dependent nucleosome phasing and insulation at CTCF bound sites or competitive binding with CTCF^23^. These results highlight the usefulness of the mutants in distinguishing cause from effect.

### Mutations reduce the chromatin residence time and bound fraction of CTCF

Fluorescence recovery after photobleaching (FRAP) was performed on the 9 mutants and WT CTCF^24,25^, using the same auxin and Dox conditions as described for the ChIP-seq and ATAC-seq. For the FRAP a circular 1 μm diameter circle was bleached, and recovery monitored over 10 minutes, recording 30 movies per condition in three replicates. While the recovery of WT CTCF largely matched prior CTCF FRAP results^24^, all mutants exhibited faster recovery consistent with reduced and/or less stable DNA-binding (**Figure 3A and S10**). We fit the FRAP curves to a reaction-dominant kinetic model ^24,25^ and estimated the residence time (duration of CTCF binding to DNA) and bound fraction (proportion of total CTCF bound to DNA) of each mutant with subsequent comparison to the WT (**Figure 3B, C**). We found a general reduction in the residence time and/or bound fraction across each mutant, with WT CTCF exhibiting the largest specific bound fraction and residence time (**Figure 3B, C**). Each mutant displayed unique parameters, regardless of the mutant category. The R377H and R377C mutants – mutations in the phosphate contacts – have similar bound fractions, but distinct residence times. The mutations in the amino acids making direct contact with DNA (R339Q, K365T, Q418R, and R448Q) have varied residence times and bound fractions, despite belonging to the same group. R448Q showed the second lowest bound fraction, despite exhibiting a residence time comparable to WT. Q418R had the lowest residence time of all mutants examined, with an intermediate bound fraction. Conversely, K365T exhibited a bound fraction and residence time most closely comparable to WT. H455R, which assists in coordinating the zinc, had the lowest bound fraction and the second lowest residence time of all examined mutants, accurately reflecting the most deleterious mutation to canonical CTCF function. R342C, which occurs in a boundary residue between the two zinc-ligand histidine residues of ZF3, showed a residence time similar to that of wild type, but had a lower bound fraction, possibly suggesting a role for boundary residues in binding efficiency. Lastly, S354F, located between the two zinc-ligand cysteine residues of ZF4, which substitutes a bulky phenylalanine in place of a serine in ZF4, showed both a moderate decrease in bound fraction as well as residence time.

**Figure 3:**
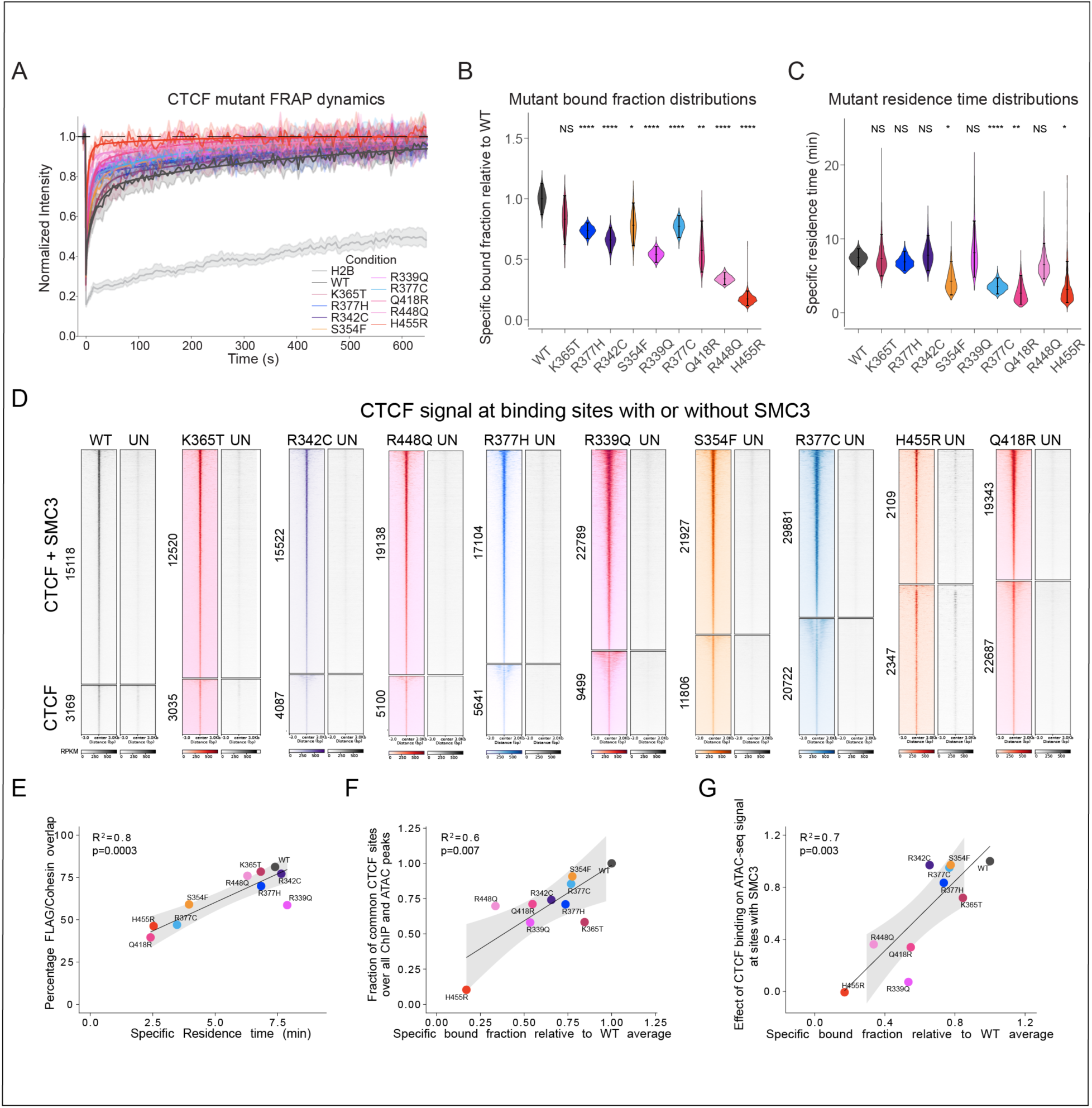
Each mutation uniquely impacts CTCF’s chromatin bound fraction, residence time and interaction with DNA. (**A)** Plots of FRAP dynamics for WT and mutant CTCF. The bold lines show the fitted model of the average recovery, and the outlines give the 95% confidence intervals (95%CIs). (**B**) Violin plots of specific bound fractions. (**C**) Violin plots of specific residence times (min). p-values were determined by bootstrapping (n=2500). (**D**) Heatmaps show the proportion of CTCF-cohesin versus CTCF only binding sites. UN corresponds to the FLAG control in untreated cells. (**E**) Correlation between residence time and the percentage of CTCF-cohesin overlap. (**F**) Correlation between the FRAP specific bound fraction relative to WT and the fraction of common CTCF sites relative to all potential binding sites. (**G**) Correlation between the FRAP specific bound fraction relative to WT and the effect of CTCF binding on ATAC-seq signal at CTCF-SMC3 sites (**Figure 2B**). For **E**, **F** and **G**, data were generated in 2 replicates, the p-values were calculated using linear regression, the shaded area corresponds to the 95% CI. See also **Figures S10-14**.

In summary, our FRAP results demonstrate that all mutants show changes in DNA binding either through an altered residence time or bound fraction. Surprisingly, some mutations strongly affect either the residence time or the bound fraction, with minimal or less of an effect on the other. This suggests that the CTCF target search (ON-rate) can be decoupled from binding stability (OFF rate) ^26^, such that some mutants affect the efficiency of the search for binding sites, without affecting the residence time of CTCF.

### The impact of mutations on the DNA-CTCF complex

To assess the impact of each mutation on the DNA-CTCF complex, we generated predictions using AlphaFold3. First, we compared the WT and K365T mutant DNA-CTCF complexes between the predicted and experimental structure and found a high concordance (**Figure S11**). We then predicted the DNA-CTCF (ZF1-11) structure for each mutant and compared the alignment against WT (**Figure S12**). Like K365T, the mutations have little effect on the overall conformation of CTCF. However, Q418R the mutant with the lowest residence time has the highest deviation from WT (RMSD=1.6 Å), although the RMSD is still <2Å, which is considered a good alignment with the WT structure. It is of note that AlphaFold3 does not take account of the effect of ions, therefore, the prediction for H455R might not be accurate since the mutation disrupts the coordination of H455 with the zinc ion. Some of the mutations found at residues making direct contact with the DNA, have lost or weakened hydrogen bonds (K365T, R339Q and R448Q). Q418R gains hydrogen bonds (**Figure S12**), which could explain the paradox of this mutation: that it has the lowest residence time but strong binding at common sites.

### CTCF mutations have a graded impact on binding and function

To understand the relationship between FRAP and ChIP-seq data, we asked whether the residence time is linked to CTCF’s binding stability and ability to block cohesin. In **Figure S13A** we show the stratified overlap of cohesin/FLAG at WT only, common and mutant only CTCF sites. Overlap with cohesin is lower at WT only and mutant only sites, consistent with their weaker CTCF and ATAC-seq signals. These changes do not reflect changes in cohesin protein levels as Western blots demonstrate that CTCF mutants do not impact the levels of RAD21 or SMC1, two subcomponents of the cohesin complex (**Figure S13B**). The proportion of global CTCF-cohesin versus CTCF only binding sites shown in the heatmaps (**Figure 3D**), demonstrate that CTCF-cohesin overlap is graded across mutants, to some extent mirroring a gene dosage effect that could occur as a consequence of altered residence time (**Figure 3C**). Indeed, analysis of SMC3 ChIP-seq revealed that the residence time was strongly associated (R^2^=0.8, p=0.002) with the percentage of CTCF-cohesin overlap for each mutant (**Figure 3E**) which reflects the impact of binding stability on CTCF’s function in loop extrusion. Conversely, the proportion of SMC3 overlapping CTCF might reflect changes in the number of CTCF peaks across the mutants since the expression level of components of the cohesin complex, SMC1 and RAD 21 do not change in CTCF mutant expressing cells. Indeed, **Figure S14A** shows a strong correlation between the percentage of SMC3 peaks overlapping CTCF and the number of CTCF peaks (**Figure S14B**).

We next compared the specific bound fraction estimated by FRAP, to the bound fraction using FLAG ChIP-seq and ATAC-seq peaks to define all potential binding sites. While we did not find a good correlation when assessing the overall CTCF ChIP-seq bound fraction (**Figure S14C**), there was a significant correlation (R^2^=0.6, p=0.008) between the chromatin bound fraction of the mutants and the fraction of mutant binding at common CTCF sites (**Figure 3F**). These data suggest that the FRAP bound fraction mostly captures the effect of the mutations on the strong, accessible binding sites. Furthermore, we found a strong correlation with the fraction bound and the overall ability of CTCF-cohesin overlapping sites to reduce accessibility, (R^2^=0.7, p=0.003), in line with CTCF functioning in this context in a dose dependent manner (**Figure 3G**).

### The relationship between cohesin and accessibility at CTCF-bound and CTCF-free sites

Consistent with previous findings^27^, our data links accessibility with CTCF binding signal strength. Furthermore, the FRAP data support an association between CTCF residence time and cohesin overlap, which is also supported by simulation analysis^28^. To separate out the contribution of CTCF and ATAC strength on SMC3 signals, we performed multivariate modelling to determine (i) the effect of CTCF strength on SMC3 overlap, adjusted for the effect of ATAC signal, and (ii) the effect of ATAC strength on SMC3 overlap, adjusted for the effect of CTCF signal. **Figure 4A** (top) shows that SMC3 enrichment independent of ATAC-seq signal, is strongly associated with CTCF signal strength in a dose-dependent manner. Only strong accessibility significantly influences SMC3 binding independent of CTCF, however, accessibility has much less contribution compared to CTCF binding stability **(Figure 4A**, bottom). Thus, CTCF signal strength plays a dominant role in determining the probability of CTCF-cohesin overlap, consistent with the FRAP and ChIP-seq data (**Figure 3E**). The heatmap in **Figure 4B** and **Table S1**, further shows that, in contrast to CTCF-bound sites, the percentage of SMC3 overlap at CTCF-free sites is increased with increasing accessibility, however signal strength at highly accessible sites is much lower than sites overlapping even weak CTCF-bound sites. The same effect is observed for the mutants, but in some cases SMC3 overlap at CTCF-bound sites is weaker than WT CTCF, in line with their reduced residence times.

**Figure 4.**
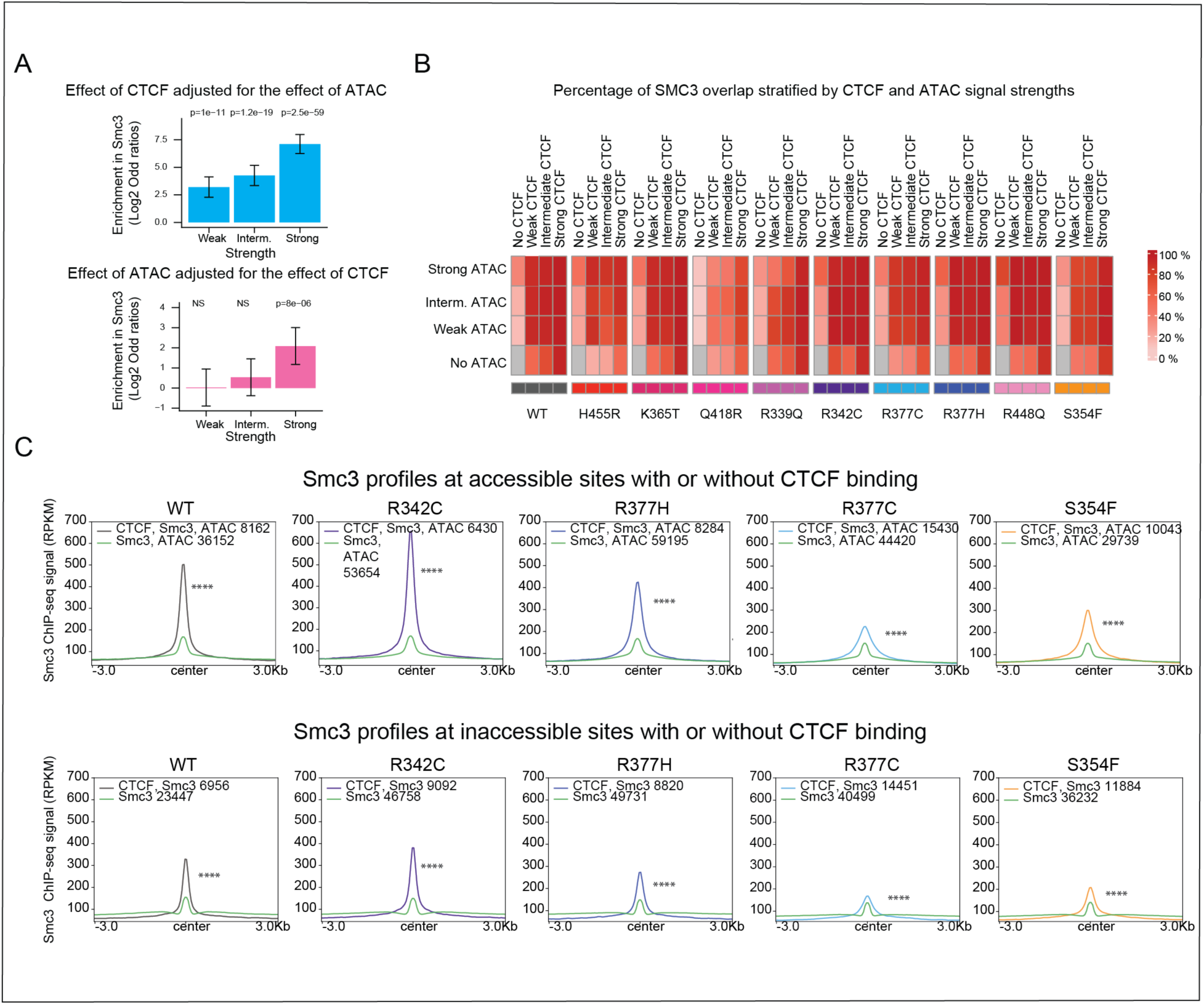
Effect of CTCF binding and accessibility on SMC3 overlap. **(A)** Bar graph showing the independent effect of CTCF (top) and ATAC-seq (bottom) signal on SMC3 enrichment in WT. The error bars correspond to the 95%CIs. The p-values were calculated using a multivariate logistic model. (**B**) Percentage of SMC3 overlap in WT and CTCF mutants, stratified by ATAC-seq and CTCF signals. **(C)** SMC3 profiles in WT and mutant CTCF. p-values were calculated using Wilcoxon tests. See also **Figures S15-16** and **Table S1**.

Overall SMC3 profiles at CTCF-bound sites confirmed that CTCF is most effective at blocking cohesin at accessible sites enriched with strong CTCF signals, and further highlight that each mutant CTCF performs this function with variable ability depending on their residence time (**Figure 3E, 4C and Figure S15**). A comparison of SMC3 profiles at CTCF bound versus unbound sites, demonstrates that other factors such as TFs, contribute very little compared to CTCF likely because residence times of TFs on chromatin are much shorter than CTCF’s^29^. Indeed, at CTCF-free sites, the SMC3 profiles are similar at CTCF-free accessible and inaccessible sites, suggesting that accessibility has little contribution to cohesin binding in the absence of CTCF. A similar trend was observed at enhancers and promoters where accessibility is higher and TFs are known to be enriched (**Figure S16**). These data suggest that for a given cell type, the subset of strong and accessible CTCF binding sites are predominantly involved in loop extrusion. However, weaker, inaccessible CTCF binding sites might also have a role in more transient, dynamic looping events with a reduced loop lifetime^30,31^. Moreover, a subset of inaccessible sites (16% in the WT) have strong CTCF signals.

### Each CTCF mutation alters gene expression and TF binding in a unique manner

To determine how CTCF mutations impact gene expression we performed RNA-seq and did an unsupervised clustering analysis, comparing gene expression in cells in which endogenous CTCF was degraded in the absence (IAA) or presence of the WT or mutant CTCF transgenes (ID). Changes in transcriptional output were determined by comparison with cells expressing WT transgenic CTCF (ID) (**Figure 5A**). Overall, 6474 genes had altered gene expression across all mutants including the IAA condition (**Table S2**). **Figure S17A** shows that the majority of differentially expressed genes (DEGs) are found only in one mutant.

**Figure 5:**
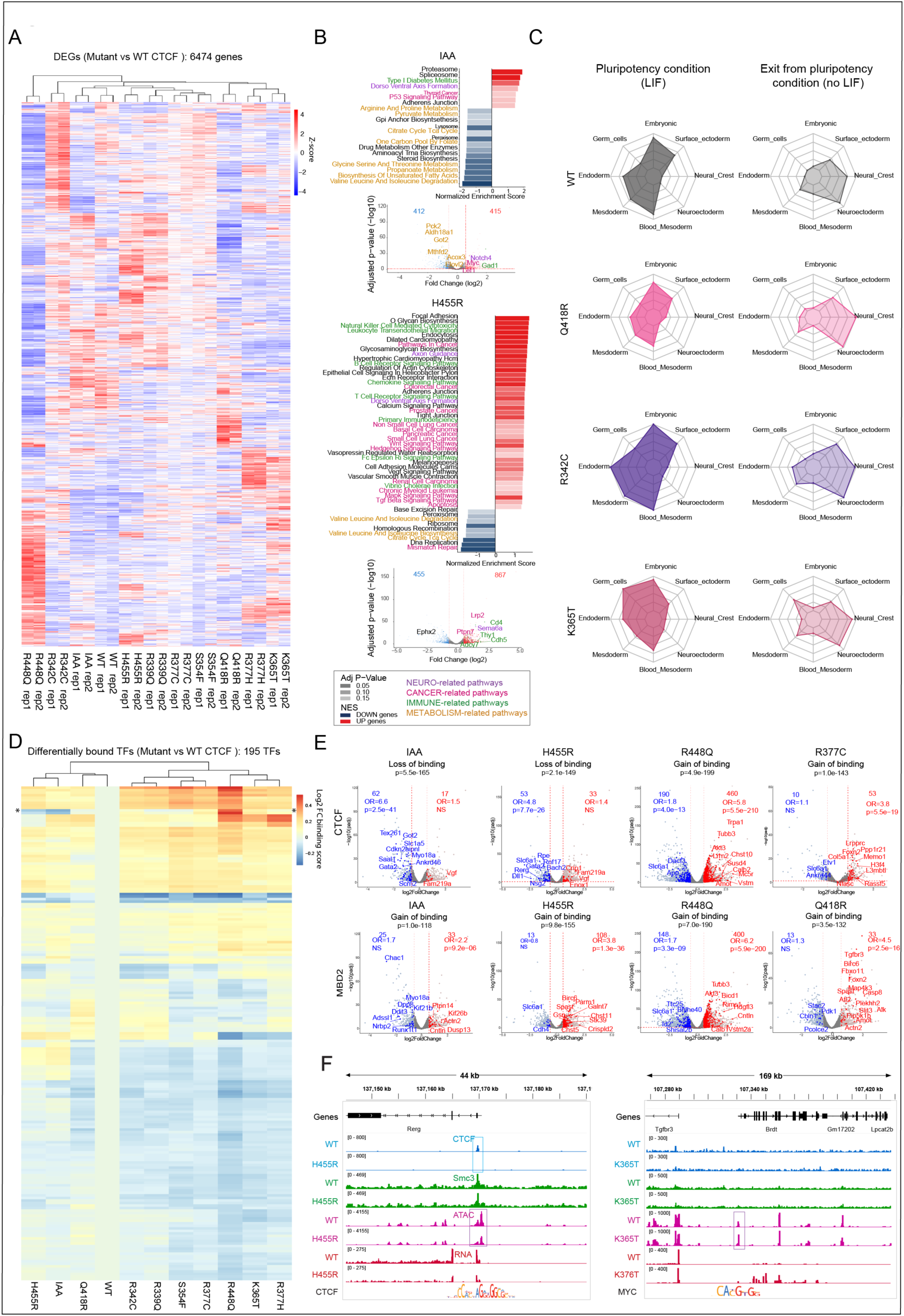
CTCF mutations alter gene expression, cellular reprogramming and TF binding. (**A)** Heatmap showing supervised clustering of the cell lines based on the expression levels of DEGs identified across the comparisons to WT. (**B**) Gene set enrichment analysis of DEGs in IAA condition and in H455R. The volcano plots below highlight the DEGs belonging to these enriched pathways. (**C**) Radar plots showing the averaged expression of developmental germ layer genes in WT and mutant mESCs cultured in LIF and no LIF conditions. (**D**) Heatmap showing predicted differentially bound TFs in WT, mutants and IAA. CTCF is highlighted with a *. (**E**) Volcano plots highlighting the differentially expressed target genes of CTCF and MBD2 in IAA and CTCF mutants. The metrics for enrichment of the target genes among the DEGs are reported on top on the volcanos (ORs and logistic p-values). (**F**) Examples of altered CTCF binding and footprinting at *Rerg* promoter (left) and altered MYC footprinting at *Brdt* promoter (right). All data in this figure were generated on 2 replicates. See also **Figures S17-20** and **Table S2-4**.

Gene set enrichment analysis using the KEGG database^32^ showed a strong enrichment in functional pathways related to cancer, brain, immune and metabolic processes among DEGs across mutants (color coded in **Figure 5B**, with genes outside these pathways represented in black). This finding is compatible with the human diseases associated with CTCF mutations. The top DEGs of these pathways are shown in the bar graphs below. Examples in **Figure 5B** show changes in pathway and gene expression in the absence of CTCF (IAA condition) and in cells expressing the mutant CTCF transgenes, H455R, S354F, R448Q and R339Q (ID condition). All other mutants are shown in **Figure S17B**.

We also observed an enrichment of dysregulated genes observed in patients with CTCF associated neurodevelopmental disorders (NDDs^9^) (**Figure S17C and Table S2**). CTCF plays an important role in imprinting, and alterations in imprinted genes could contribute to the CTCF-dependent neurodevelopmental phenotype as suggested by congenital imprinting disorders^33^. Indeed, we found a strong enrichment of imprinted genes across all mutants (**Figure S17D and Table S2**).

Together, our analyses demonstrate that although each mutant exhibits a unique impact on binding and gene expression, all mutants affect common pathways, consistent with phenotypes linked to human CTCF-associated disorders. Differences in the impact of each mutant provide insight into CTCF-associated NDD inter-patient variability. Interestingly, cells in which endogenous CTCF is degraded (IAA condition) show fewer gene expression changes compared to mutant expressing cells, except for the R377C mutation (**Figure S17B**). Some mutants have alterations in many pathways (R399Q), while others highlight changes in only a few (Q418R and R377C) as shown in **Figure 5B and Figure S17B**, suggesting that in the latter, the alteration in gene expression may not accumulate in specific pathways.

### Mutant specific alterations in transcriptional networks impact exit from pluripotency

To investigate, whether mutant specific transcriptional networks could potentially affect exit from pluripotency and the ability to differentiate we compared expression profiles of WT and a selection of mutant CTCF expressing cells before and after 72 hours of LIF withdrawal^34^. The mutants analyzed were selected because they cover a range of CTCF residence times as measured by FRAP. R342C exhibits a similar residence time to WT, K365T an intermediate residence time and Q418R has one of the lowest residence times. RNA-seq revealed that each mutant demonstrated a distinct differentiation trajectory compared to WT cells, which are most skewed towards a neuroectodermal fate (**Figure 5C** and **Table S3**). A subset of the DEGs is shown in **Figure S18**.

In the mutants, we identified several differentially expressed imprinted genes, such as H19 and PEG10, as well as ZFP57, which plays an important role in regulating imprinting. To further assess the biological relevance of the differentiation behavior, we annotated DEGs using the DECIPHER developmental disorder database (https://www.deciphergenomics.org/ddd) (**Table S3**). We found 44 genes associated with developmental disorders, including NDDs such as *Camk2a*, *Cnksr2*, *Sobp* that are involved in intellectual disability, and *Dchs1* which is responsible for periventricular neuronal heterotopia. In addition, we detected genes responsible for congenital connective tissue syndromes (*Fras1* in Fraser syndrome and *Fbn1* in Marfan syndrome) as well as genes involved in skeletal dysplasia (*Sox9*) and congenital cardiomyopathy (*Speg*). While CTCF-related congenital disorders are most commonly characterized by neurodevelopmental symptoms, patients can also exhibit other congenital anomalies such as heart defects, craniofacial malformations or cleft palate^10^, suggesting the biological relevance of the changes in differentiation behavior observed in our system.

### Transcriptional changes correspond to predicted alterations in TF binding at promoters

To investigate whether mutant specific transcriptional changes could be explained by altered TF binding, we performed a footprinting analysis of ATAC-seq data using the TOBIAS pipeline ^35^ in cells which have endogenous CTCF degraded in the absence (IAA) or presence of WT or mutant transgenes (ID). For this analysis replicates were merged and comparisons made with WT ID cells. Overall, 195 TFs had predicted differential binding across all mutants including the IAA condition. As a proof of principle, we detected predicted differential binding for CTCF, which has altered binding in the mutants. Numerous other predicted differentially bound TFs (n = 194) were identified as shown in **Figure 5D, S19A** and **Table S4.** About one third were found in only one mutant (**Figure S19B and Table S4**). Annotation of differentially bound TFs revealed a significant enrichment for TFs associated with NDDs (Odd Ratio =2.2, p=0.02) (**Table S4**).

We overlapped predicted differentially bound TFs with a region of 2kb around the promoters of DEGs in cells expressing different CTCF mutants, to analyze the impact of predicted differential binding of TFs on gene expression and observed a strong enrichment (**Figure 5E, Figure S20** and **Table S4**). Differentially expressed target genes were also significantly enriched in genes associated with developmental disorders (OR=3.6, p=7x10^-36^) and in differentially expressed genes found in CTCF associated NDDs (OR=4, p=10^-63^). It is of note, that the majority of the predicted changes in TF binding at the promoters of DEGs do not overlap CTCF ChIP-seq peaks or predicted differential CTCF binding. This data suggests that many of the mutant-mediated transcriptional changes could be explained by indirect effects resulting from the disruption of specific TF pathways.

Figure 5E **and Figure S20** highlight the impact of two of the top predicted differentially bound TFs (CTCF and MBD2) at promoters of DEGs in different mutants. These are shown in volcano plots for cells in which endogenous CTCF is degraded (IAA) and in cells expressing different CTCF mutations (ID). MBD2 is a methyl-sensitive factor that belongs to the NurD chromatin remodeling complex, that has been implicated in cancer and NDDS. NurD is also required for exit from pluripotency and lineage commitment^36–39^. For each CTCF mutant, predicted differentially bound TFs target the promoters of a distinct set of up (red) and down (blue) regulated genes. Both CTCF and MBD2 exhibit an overall activator profile with a positive association between binding and expression (Figure 5E **and Figure S20**). However, for some mutants, a weaker but significant negative association was also observed suggesting that both factors can act as activators or repressors depending on the cell-specific genomic context, consistent with previous reports^38,40^.

Examples of altered CTCF binding at the promoter of differentially expressed *Rerg* and predicted differentially bound MYC at the promoter of differentially expressed *Brdt*, are shown in Figure 5F. Loss of CTCF binding at the promoter of the *Rerg* gene leads to a decrease in its expression, while expression of *Brdt* increases in line with increased predicted binding of MYC at its promoter. The latter is an example of an indirect effect because CTCF is not bound at the promoter of this gene. In sum, combined analysis of RNA-seq and ATAC-seq highlight the variable effects of each CTCF mutation, connecting gene expression changes with 195 predicted differentially bound TFs, to provide a deeper understanding of the factors that underlie the unique transcriptomic profile of each CTCF mutation and their impact on developmental processes particularly relevant to brain disorders.

### Chromatin interactivity and insulation are linked to mutant binding properties

Hi-C demonstrated that each mutant has a distinct interaction profile and exhibits (i) variable loss of intra TAD interactions and boundary strength as well as (ii) aggregated loop strength. The IAA condition, in which CTCF is degraded shows a loss of intra TAD interactions and a gain of inter TAD interactions as well as the most reduction in aggregated loop strength (Figure 6A), while mutants display their own unique effects. No major compartment changes were observed as shown by the interaction matrices at the chromosome level (**Figure S21A**).

**Figure 6:**
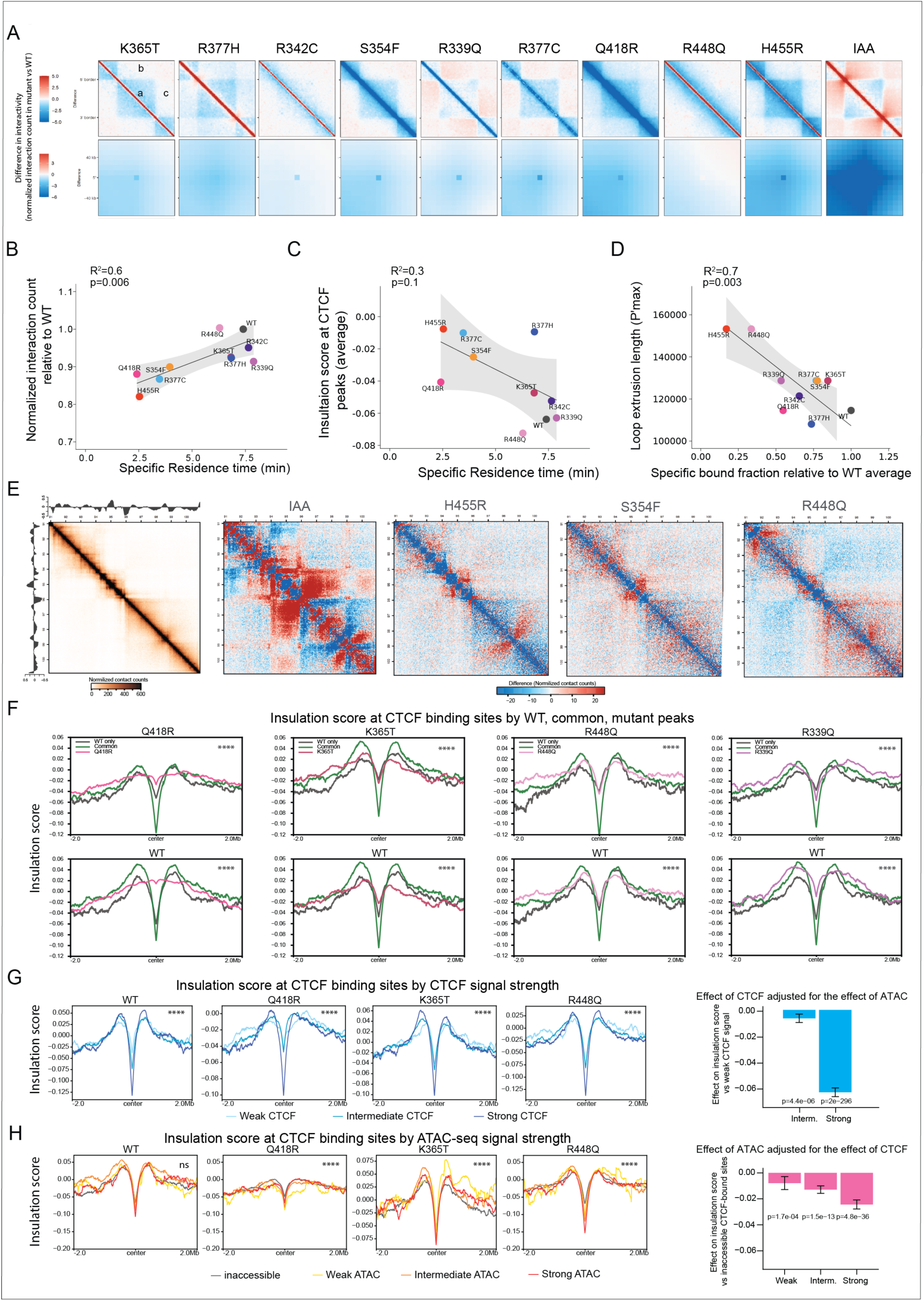
CTCF mutations alter chromatin interactivity. **(A)** The top panels show the aggregated differential TAD analysis between mutants and WT**. (a)** Reflects the intra-TAD interaction, **(b)** and **(c)** highlight the inter-TAD interactions. The lower panels show the aggregated differential peak analysis. The data was generated by Hi-C on 2 replicates. (**B**) Correlation between the interaction counts and the FRAP residence times. (**C**) Correlation between the insulation score at CTCF peaks and the FRAP residence times. (**D**) Correlation between the loop extrusion length and the FRAP bound fractions. (**E**) Example of differential interactions between mutants and WT (right). The left matrix shows the interaction in WT within a 10 Mb region (40 kb resolution) with the insulation score on the side. (**F**) Profiles show the averaged insulation score at WT only, common and mutant only binding sites. (**G**) Profiles show the insulation score at CTCF binding sites stratified by CTCF signals. The bar graph shows the independent effect of CTCF signal on insulation score. (**H**) Profiles show the insulation score at CTCF binding sites stratified by ATAC signals. The bar graph shows the independent effect of ATAC signal on insulation score. For **F, G** and **H**, p-values reported in the profiles were calculated using Kruskal-Wallis tests. The fitted estimates for the bar graphs in (**G**) and (**H**) were obtained using a mixed multivariate model. The error bars correspond to the 95%CI. See also **Figures S21-24**.

Consistent with our finding that cohesin overlap is correlated with CTCF mutant residence time as determined by FRAP analysis (Figure 3E), we found that residence time is also associated with chromatin interactivity and insulation score (Figure 6B**, C**).The correlation with the latter was modest and did not reach significance, probably because of the lower Hi-C resolution (10 kb) compared to ChIP-seq. The loop extrusion length (P’max^41^) for each mutant is associated with the fraction that is bound to chromatin (Figure 6D), with the longest loops associated with mutants that have the lowest bound fraction, indicating a lower density of CTCF-cohesin anchors in these mutants (Figure 3B). The variability in the interaction profile of each mutant can be clearly seen across a 10Mb region of chromosome (Figure 6E **and Figure S21B**).

To further examine the impact of each mutant on chromatin organization, we first compared the aggregated insulation score at lost sites upon CTCF degradation (comparison of IAA versus WT-ID condition) (**Figure S21C**) using a 2Mb window centered on CTCF peak summits. The insulation scores were generated from Hi-C data at a 10kb resolution. This analysis confirmed that the mutants partially retain insulation at lost sites depending on their binding stability. We then analyzed the insulation score at WT only, common and mutant only binding sites (Figure 6F **and Figure S22A**). This analysis revealed that the strong, common accessible binding sites with the highest cohesin overlap, have the highest insulation compared to WT only and mutant only sites. At the WT only sites (comparison of WT-ID versus mutant-ID), some mutants exhibit weak insulation which is not detected when CTCF is degraded (Figure 6F**, Figure S22A** and **Figure S22B**), suggesting there might be weak mutant CTCF binding (non-significantly detected by ChIP-seq) at some of these sites. Whether insulation scores at WT only and mutant only sites are lower or higher than each other depends on the residence time and cohesin overlap of each mutant, in line with what we observed in Figure 3C**, E and 6C**. These studies further demonstrate that imaging and molecular analyses can be functionally integrated to provide new insight into mutant specific effects.

Since WT and mutant only sites are associated with weaker CTCF peaks and lower accessibility, we examined the insulation score at CTCF binding sites, stratified by CTCF and ATAC-seq strength. The profiles in Figure 6G**, 6H, S22C and S22D** indicate that stronger CTCF signals are associated with stronger insulation, while accessibility has a more modest impact. To quantify their independent effects, we performed a mixed multivariate model, the results of which are consistent with the analysis of SMC3 overlap shown in Figure 4A.

### Changes in gene expression are linked to changes in chromatin interactivity

To determine if changes in gene expression are linked to altered chromatin folding, we analyzed interactions involving the promoters of genes that were differentially expressed and found that the promoters of overexpressed genes were enriched in gained chromatin loops (Figure 7A). Enrichment was not observed among the under-expressed genes, suggesting that under-expression observed in mESC expressing mutant CTCF proteins might be an indirect effect, while overexpression might reflect a direct CTCF-mediated effect on loop extrusion. This is supported by the sub-analysis of overexpressed and underexpressed genes shown in **Figure S23,** focusing on loops with both anchors overlapping a CTCF binding site, which show a similar result. These loops are more likely to capture the direct effect of CTCF on loop extrusion. This is shown in examples in Figure 7B and **Figure S24A** which depict gain of interactions from CTCF binding sites toward the promoters of overexpressed genes, including *Msh6*, *Epcam*, *Foxn2* and *Cox7a2l* (Figure 7B and **Figure S24B)**. Interestingly, all these genes are implicated in tumorigenesis^42–48^

**Figure 7:**
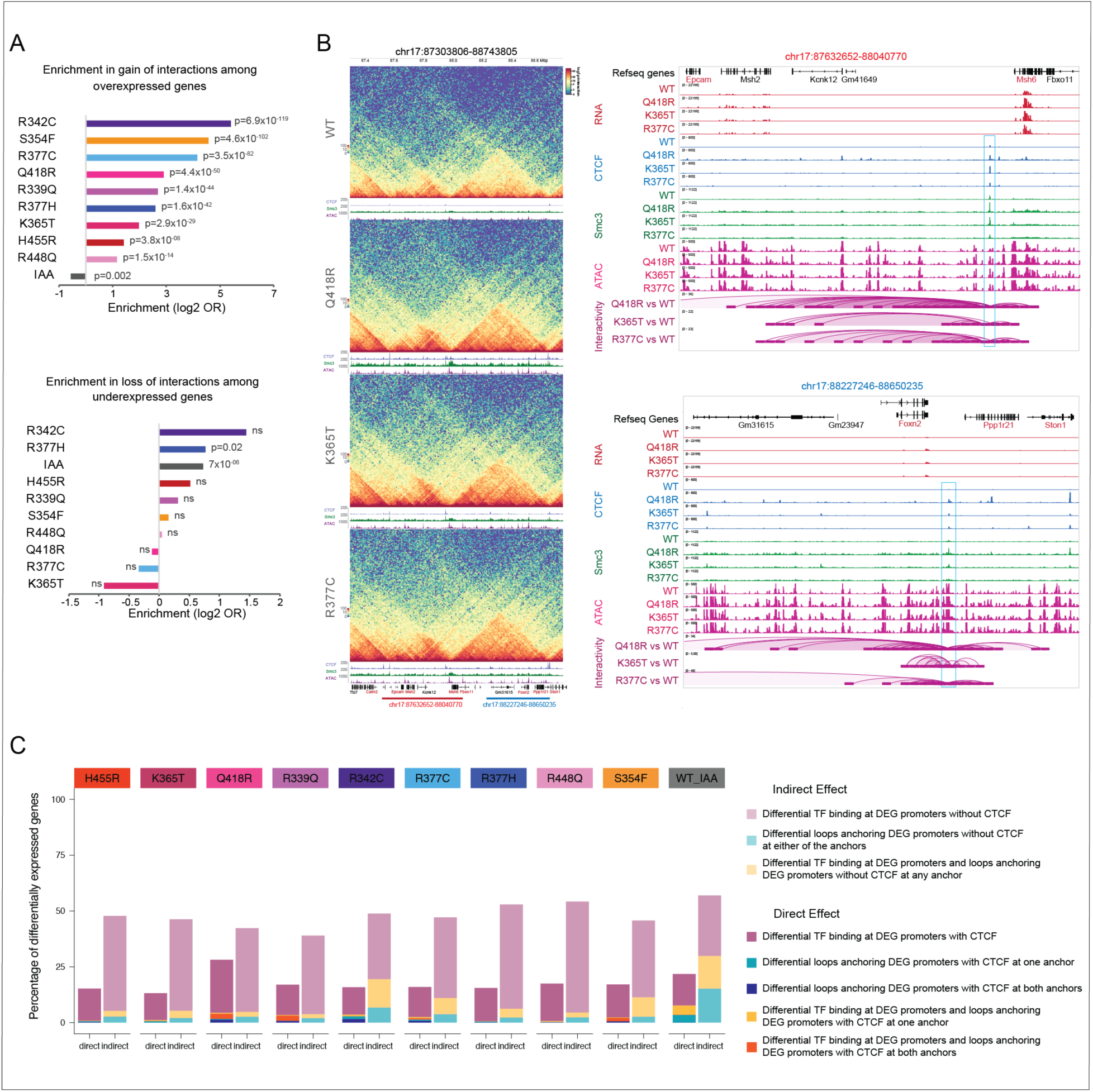
Changes in gene expression are linked to changes in chromatin interactivity but to a lesser extent) than changes in TF binding at gene promoters: (**A**) Bar graphs showing the enrichment of over-expressed (top) or under-expressed (bottom) genes in gained (top) or lost (bottom) loops in IAA and mutants compared to WT. (**B**) Example of 2 loci (blue and red rectangles) with a direct effect of gain in CTCF binding and chromatin interactivity. The interaction matrices (left) show both gain of intra- and inter-TAD interactions in some mutants compared to WT. The left panels show the zoom-in tracks of these loci. Only significant differential chromatin loops are shown. Overexpressed genes are highlighted in red. **C**) Bar graph showing the percentage of DEGs resulting from direct or indirect effect of CTCF binding distinguishing loop dependent and independent effect.

### Most of the change in expression are the result of indirect effects

Our data show that most CTCF mutations have more impact on gene expression than the IAA condition. Indeed, CTCF degradation has been shown to induce limited gene expression changes ^14,15^. To ask whether this could be explained by indirect effects which depend on cell-specific transcriptomes, we estimated the direct versus indirect effects of CTCF mutations on gene expression. Since CTCF can act as a traditional transcription and chromatin regulator ^40^, we defined indirect CTCF effects as (i) differentially expressed genes (DEGs) with a predicted differentially bound TF at the promoters that do not overlap a CTCF ChIP-seq peak or a predicted differentially bound CTCF site, and (ii) DEGs with differential looping of the promoter without CTCF binding at either loop anchor. Conversely, for direct effects, we required (i) the presence of a CTCF peak and/or a predicted differentially bound CTCF site at the DEG promoter or (ii) loops anchoring the DEG promoter with at least one loop anchor overlapping a CTCF binding site.

Figure 7C shows that indirect effects account for approximately twice as many DEGs compared to the direct effects, except for Q418R. This mutant has a short residence time but a high number of peaks, supporting the hypothesis that unstable peaks might be involved in dynamic and functional looping. Furthermore, a higher percentage of DEGs can be explained by changes in TF or CTCF binding at gene promoters, rather than by changes in chromatin interactivity in all the mutants. For the IAA condition, the contribution of differential loops to expression changes is greater. A similar result was observed when segregating up- and down-regulated genes but with a higher proportion of direct versus indirect effects in the IAA condition for down-regulated genes consistent with loss of loops upon CTCF degradation (**Figure S24B**). These findings suggest that although CTCF mutations induce changes in chromatin looping, the primary mechanism driving alterations in gene expression is loop-independent and mediated by changes in transcription factor binding.

## DISCUSSION

Using a combination of genetic, imaging, molecular, AlphaFold3 and bioinformatic approaches we have examined the impact of nine, high frequency cancer associated CTCF mutations in ZFs that make contact with the core consensus binding motif. Our studies reveal that the specificity of interaction between the ZFs in CTCFs binding domain and its target DNA sequence is a key determinant of attachment. However, if the same sequence is found in both accessible and inaccessible chromatin, CTCF will preferentially bind accessible sites as highlighted by the fact that mutants bind common accessible sites in preference to less accessible WT only sites. This is underscored by the strong correlation between the fraction of chromatin bound mutants detected by FRAP, and mutant binding at common, accessible CTCF sites. These findings are consistent with studies from Dirk Schubeler’s group showing that CTCF can bind both accessible and inaccessible sites, but that the latter are less likely to be involved in insulation ^27,49^. We also observed a good correlation between each mutant’s residence time and CTCF-cohesin overlap, indicating that binding stability is important for blocking cohesin. A similar, but more modest correlation was found between residence time and chromatin interactivity as well as insulation score, linking binding stability with cohesin overlap as well as loop and boundary formation.

By dissecting out the contribution of CTCF signal strength and accessibility to SMC3 overlap, we were able to reveal the dominant effect of CTCF. SMC3 overlap at accessible and inaccessible CTCF-bound sites is stronger than overlap at accessible, CTCF-free sites, including enhancers and promoters, likely because TFs have a short residence time compared to CTCF ^29^. Our data also demonstrate that strong accessible CTCF bound sites are associated with higher insulation, consistent with higher SMC3 stalling and more stable loops.

In principle, these features could have been uncovered by analyzing WT CTCF in the context of cohesin overlap and accessibility. However, collectively the graded effects of the mutants act as perturbations that lend support to our models, underscoring how different aspects of CTCFs binding properties contribute. Moreover, they reveal that each mutant’s search for binding sites can be decoupled from its chromatin bound residence time, as previously demonstrated for a mutant form of CTCF with ZF8 deleted ^50^. For example, the Q418R mutant binds an intermediate number of sites but at these sites it is has a very low residence time and is less competent at blocking cohesin. In contrast, R448Q binds fewer sites with a similar residence time compared to WT CTCF, and it is slightly better at blocking cohesin than WT protein. The low chromatin bound fractions of H455R and R448Q, which have very different chromatin residence times, are correlated to an increased loop length of both mutants (P’-max), which can be explained by the unobstructed passage of cohesin past what would normally be a bound CTCF site.

The crosstalk between accessibility and CTCF binding goes in both directions, such that accessibility affects binding and binding affects accessibility. Binding at accessible sites, narrows and reduces the ATAC-seq peak. Evidence for this being a CTCF driven effect comes from analysis of the CTCF mutants. Those with low residence times (H455R and Q418R) are unable to function in this capacity, while the R342C mutant has a stronger effect compared to WT CTCF. We speculate that these differences reflect decreased accessibility at the bases of CTCF-SMC3-mediated loops, because in the absence of SMC3, where loops cannot form there is no reduction in accessibility.

While loss or gain of CTCF binding at promoters can account for direct mutant specific effects on accessibility and gene expression, we could only demonstrate a direct effect of gained CTCF binding associated with gained loops involving the promoters of over-expressed genes. Under-expression could mostly reflect indirect effects of CTCF binding alterations. Indeed, all mutants have their own unique indirect impact in globally changing accessibility at sites where CTCF is not bound. This effect links changes in TF footprinting with altered transcriptional output, explaining most of the mutant specific effects that we observed. We also identified a higher proportion of DEGs mediated by changes in TF or CTCF binding at the gene promoters than by changes in chromatin looping, suggesting that the primary mechanism driving alterations in gene expression might be loop independent. Indeed, the impact of each mutant on TF expression and altered TF binding at the promoters of differentially expressed genes is less likely to give rise to detectable changes in interactivity by Hi-C, consistent with our finding that SMC3 signals at CTCF-free sites are far weaker than at CTCF-bound sites, and thus likely to be involved in weaker, more dynamic loops.

Aside from the R377C mutant, we observed fewer changes in gene expression in IAA treated cells compared to CTCF mutant expressing cells, which can be explained by the predominant contribution of indirect changes mediated by TF binding on the mutant-specific transcriptome. Although the mutants all uniquely affect gene expression pathways, we detected enrichment in pathways affecting the brain, immune system, cancer and metabolism, which is consistent with the clinical setting in which the mutations are found, namely cancer and brain disorders. The added effect on immune and metabolic pathways provides insight into other changes that could occur in CTCF mutant-mediated diseases.

CTCF mutations also affect the trajectory of exit from pluripotency. Indeed, while both WT and CTCF mutant cells tend to acquire a neuroectodermal expression profile upon LIF withdrawal, we observed mutant-specific transcriptional signatures of other germ layers, suggesting a more heterogenous differentiation trajectory, which could be confirmed in future work by scRNA-seq. This finding provides a potential mechanism for CTCF mutation-associated neurodevelopmental disorders. Indeed, while CTCF has been involved in neuro-progenitor maintenance and survival^51^, our analyses suggest that CTCF mutations might also alter brain development by affecting germ layer commitment.

Taken together the binding domain mutants we have analyzed here, provide a new appreciation of CTCF’s bidirectional relationship with chromatin, its ability to bind and function in different contexts as well as the potential impact of each mutation in clinical settings. Furthermore, our analyses provide a better understanding of how any genetic or epigenetic disorder that alters the landscape of chromatin can in turn impact CTCF binding and function.

## LIMITATIONS OF THE STUDY

In our complementation system, the expression of the transgene is lower than the endogenous CTCF. Since CTCF binding and function is dose-dependent, our analyses of CTCF binding sites might be biased toward higher affinity sites. To distinguish the binding of mutant transgene from residual endogenous WT CTCF in the ID condition, we performed ChIP-seq using a FLAG antibody which resulted in multiple non-specific binding sites. To address this issue, we used a stringent peak calling resulting in a lower number of CTCF binding sites compared to previous reports^15^.

In this study, we found decreased accessibility at CTCF-SMC3 sites and hypothesize a link with the loop extrusion process. However, despite the evidence from the mutant perturbation, the reduction in overall accessibility at these sites could reflect a preference for CTCF binding involved in insulation at less open chromatin regions, which we cannot entirely rule out.

We found that most of the differences in transcriptomic profiles observed in CTCF mutant cells were the result of indirect and loop-independent effects. However, our Hi-C data has a resolution of 10 kb, which could lead to an underestimation of CTCF loop-dependent expression changes.

## RESOURCE AVAILABILITY

### Lead contact

Further information and requests for resources should be directed to and will be fulfilled by the lead contact, Jane Skok.

### Materials availability

The WT and mutant CTCF mESC lines are available upon reasonable request.

### Data and code availability

All raw and processed sequencing data files will be deposited at NCBI’s Gene Expression Omnibus (GEO) upon publication. FRAP analysis was performed using custom Python scripts available at https://github.com/ahansenlab/ZNF143_analysis_code/tree/main and https://zenodo.org/doi/10.5281/zenodo.14056922 ^52^. No other custom code/software was used in the study. The publicly available software used is indicated in the Method details.

## Acknowledgements

This work was supported by 1R35GM122515 (J.S) and NIH P01CA229086 (J.S). A.T. was supported by the American Cancer Society (RSG-15-189-01-RMC) and St. Baldrick’s foundation (581357). ASH additionally acknowledges support from US National Institutes of Health grants R00GM130896, DP2GM140938, R33CA257878 and UM1HG011536, National Science Foundation grant 2036037, the and a Pew-Stewart Cancer Research Scholar grant. GC and GJ were supported by fellowships from the NCC.

The authors thank Skok lab members for helpful scientific discussions, New York University School of Medicine High Performance Computing Facility (HPCF) for computing technical support, Adriana Heguy and the Genome Technology Center (GTC) core for sequencing efforts, NYU Flow Cytometry and Cell Sorting Center for FACS analysis and sorting. GTC is a shared resource partially supported by the Cancer Center Support Grant P30CA016087 at the Laura and Isaac Perlmutter Cancer Center.

## Author contributions

These studies were designed by J.A.S., C.D. and M.N. The analysis was performed by C.D and T.S. G.J, P.L and F.F.D.R. established the CTCF mutant cell lines with the guidance of E.P.N. Sequencing libraries were performed by G.J., G.C., R.V., N.K. A.T. reviewed the manuscript. FRAP analysis was performed by C.C.K, D.N.N and A.S.H. The paper was written by J.A.S and C.D.

## STAR Methods

### Key resources table

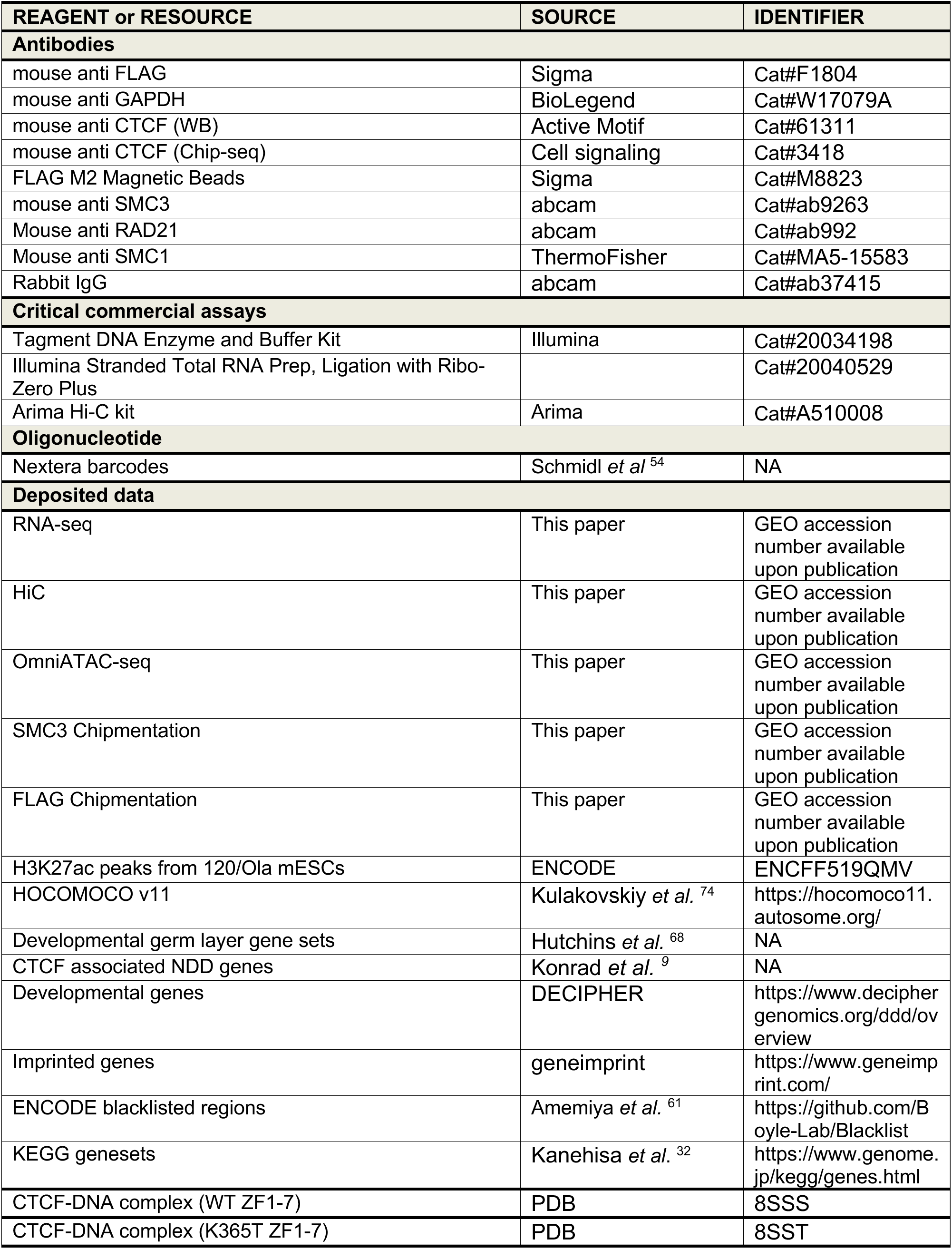
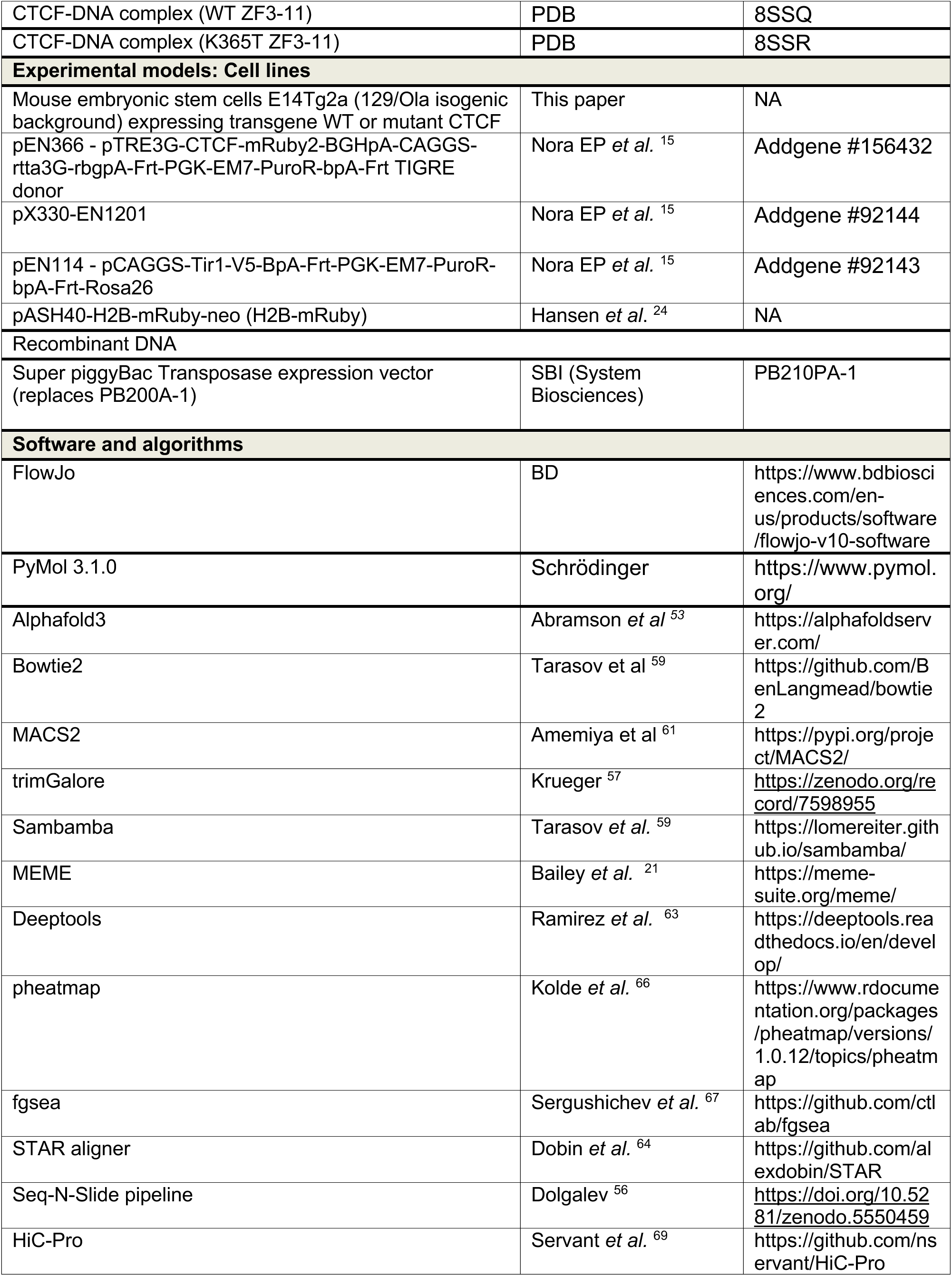
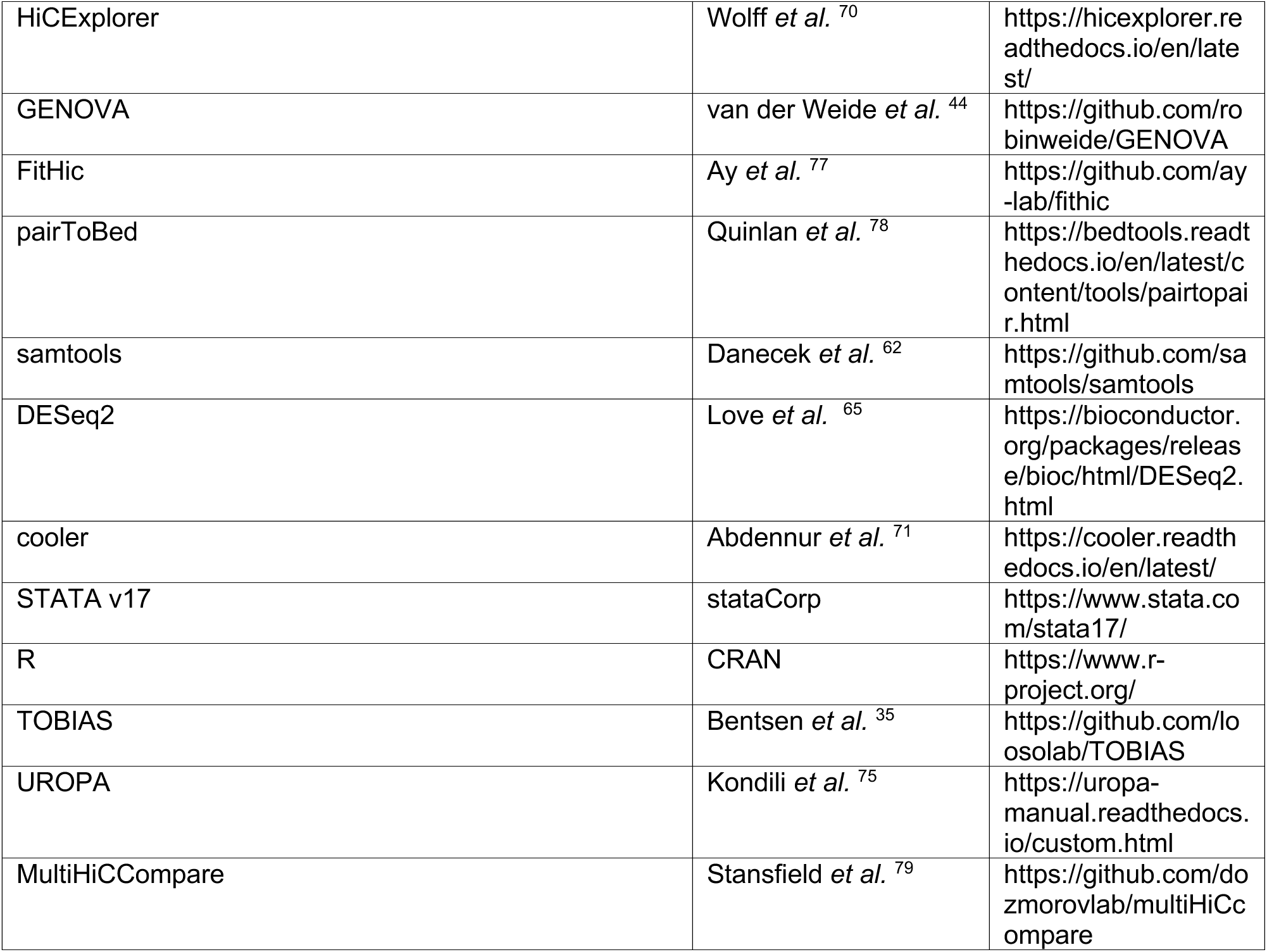

## EXPERIMENTAL MODEL AND SUBJECT DETAILS

### Cell lines

Mouse embryonic stem cells E14Tg2a (karyotype 19, XY; 129/Ola isogenic background) and all clones derived from these were cultured under feeder-free conditions in 0.1% gelatin (Sigma ES-006-B) coated dishes (Falcon, 353003) at 37°C and 5% CO2 in a humidified incubator. The cells were grown in DMEM (Thermo Fisher, 11965-118) supplemented with 15% fetal bovine serum (Thermo Fisher, SH30071.03), 100 U/ml penicillin - 100 μg/ml streptomycin (Sigma, P4458), 1 X GlutaMax supplement (Thermo Fisher, 35050-061), 1 mM sodium pyruvate (Thermo Fisher, 11360-070), 1 X MEM non-essential amino-acids (Thermo Fisher, 11140-50), 50 μM b-mercaptoethanol (Sigma, 38171), 10^4^ U/ml leukemia inhibitory factor (Millipore, ESG1107), 3 μM CHIR99021 (Sigma, SML1046) and 1 μM MEK inhibitor PD0325901 (Sigma, PZ0162). The cells were passaged every alternate day by dissociation with TrypLE (Thermo Fisher, 12563011). For the exit from pluripotency experiments, mESCs were cultured for 3 days in the same medium but without the addition of LIF (leukemia inhibitory factor, CHIR99021 (Sigma, SML1046)) and MEK inhibitor (PD0325901).

## METHOD DETAILS

### DNA constructs

Construction of vector for cloning transgenic, doxycycline-inducible expression of WT and mutant mouse *Ctcf* cDNA were obtained from GenScript in pUC19 vectors. The cDNA was amplified such that it harbors an AflII sequence at the 3’ end of the gene and fused to a FLAG tag (that harbors NotI sequence) at the 5’ end with the help of a fusion PCR. The resultant fragment was digested with NotI and AflII. The *Ctcf* gene was removed from pEN366^15^ by digesting with the same enzymes. This backbone was used for insertion of each *Ctcf* mutant.

In brief, the cDNA region corresponding to each of the C and N terminals and zinc fingers were PCR amplified in such a way that it included a short stretch of the 5′ and/or 3′ region of the neighboring fragment to be connected. The desired PCR products were then annealed, amplified by PCR and cloned into the NotI and AflII sites of the pEN366 backbone (Addgene #156432). All the constructs were verified by DNA sequence analysis. For all transgenes, the final vector harbors an N terminal 3 X FLAG tag and a C terminal *mRuby* as in-frame fusion to WT and mutant *Ctcf*. The vector also harbors a *TetO-3G* element and *rtTA3G* for doxycycline induced expression of the transgene, and homology arms surrounding the sgRNA target site of the *Tigre* locus for locus-specific insertion. The selection of stable integrants was achieved by virtue of *FRT-PGK-puro-FRT* cassette. Further details of the vector are described elsewhere ^15^. The vector pX330-EN1201 ^15^ harboring spCas9 nuclease and *Tigre*-targeting sgRNA was used for targeting of *Tigre* locus (Addgene #92144).

### Gene targeting

Mouse embryonic stem cell E14Tg2a harboring *Ctcf-AID-eGFP* on both alleles and a knock-in of pEN114 - *pCAGGS-Tir1-V5-BpA-Frt-PGK-EM7-PuroR-bpA-Frt-Rosa26* at *Rosa26* locus was used as the parental cell line for making all the transgenes ^15^. pEN366 derived vectors harboring the rescue transgenes (WT and mutant *Ctcf*) were used for targeting transgenes to the *Tigre* locus (clone ID# EN156.3.5) ^14^. For nucleofections, 15 μg each of plasmids harboring the transgenes and 2.5 μg of those with sgRNA targeting the *Tigre* locus was used. Nucleofection were performed using Amaxa P3 Primary Cell kit (Lonza, V4XP-3024) and 4D-transfector. 2 million cells were transfected with program CG-104 in each case. The cells were recovered for 48 h with no antibiotic followed by selection in puromycin (1 μg/mL) (Thermo Fisher, A1113803). Single colonies were manually picked and expanded in 96 well plates. Clones were genotyped by PCR and flow cytometry was performed to confirm that the level of expression of transgenes were comparable. All the clones that were used for the analyses were homozygous for the integration of the transgenes and their levels of expression were comparable.

### Induction of auxin inducible degradation of CTCF and doxycycline induced expression

For degradation of endogenous CTCF, the auxin-inducible degron was induced by adding 500 μM indole-3-acetic acid (IAA, chemical analog of auxin) (Sigma, I5148) to the media. Expression of transgenes was achieved by the addition of doxycycline (Dox, 1 μg/ml) (Sigma, D9891) to the media. The cells were treated with IAA and/or Dox for 2 days.

### Western Blotting

mESCs were dissociated using TrypLE, washed in PBS, pelleted and used for western blotting. Approximately 2 million cells were used to prepare cell extract. Cell pellets were resuspended in RIPA lysis buffer (Thermo Fisher, 89900) with 1X HALT protease inhibitors (Thermo Fisher, 78430), incubated on ice for 30 min, spun at 4°C at 13,000 rpm for 10 min and supernatant was collected. For the western blot of CTCF, low salt lysis buffer (0.1 M NaCl, 25 mM HEPES, 1 mM MgCl2, 0.2 mM EDTA and 0.5% NP40) was used supplemented with 125 U/ml of benzonase (Sigma E1014). Protein concentration was measured using the Pierce BCA assay kit (Thermo Fisher, 23225). 20 μg of protein were mixed with Laemmli buffer (Biorad, 1610737) and β-mercaptoethanol, heated at 95°C for 10 min and run on a Mini-protean TGX 4%-20% polyacrylamide gel (Biorad, 456-1095). The proteins were transferred onto PVDF membranes using the Mini Trans-Blot Electrophoretic Transfer Cell (Bio-Rad, 170-3930) at 80 V, 120 mA for 90 min. PVDF membranes were blocked with 5% BSA in 1 X TBST prior to the addition of antibody. The membranes were probed with appropriate antibodies overnight at 4°C, mouse anti-FLAG antibody (Sigma, F1804; 1: 1,000 dilution), GAPDH (BioLegend, W17079A; 1:5,000 dilution), CTCF (active motif, 61311; 1: 2,000 dilution), SMC1 (ThermoFischer, MA5-15583. 1:1000 dilution), RAD21 (abcam, ab992,1:2000 dilution). Membranes were washed five times in PBST (1 × PBS and 0.1% Tween 20) for 5 min each and incubated with respective secondary antibodies in 5% BSA at room temperature for 1 h. The blots were rinsed in PBST and developed using enhanced chemiluminescence (ECL) and imaged by Odyssey LiCor Imager (Kindle Biosciences).

### Flow cytometric analysis

Cells were dissociated with TrypLE, washed and resuspended in MACS buffer for flow cytometric analysis on LSRII UV (BD Biosciences). Analysis was performed using the FlowJo software.

### Cell culture – FRAP

Control cells expressing H2B-mRuby were engineered using the same background as the CTCF transgene expressing cells, E14Tg2a mESCs. Briefly, pEN114 - *pCAGGS-Tir1-V5-BpA-Frt-PGK-EM7-PuroR-bpA-Frt-Rosa26* was used as the parental cell line to be nucleofected by the pASH40-H2B-mRuby-neo with the PiggyBac transposase vector (SBI, PB210PA-1). Halo-H2B plasmid was previously published in Hansen *et al,* ^24^ and we replaced the Halotag with mRuby. For the FRAP experiment, CTCF and H2B expressing cells were cultured for two days on 35 mm no 1.5H glass-bottom imaging dishes (MatTek, Ashland, MA, P35G-1.5-14C) coated with Geltrex (Gibco, A1413201) according to manufacturer’s instructions. 48 hours prior to the experiment, the culture medium was changed to medium supplemented with 500 μM 3-Indoleacetic acid (IAA; Sigma-Aldrich, I2886-5G) and 1 μg/mL doxycycline (Sigma-Aldrich, D3072-1ML). Just prior to imaging, the medium was changed to imaging medium: phenol red free DMEM with all other aspects of the medium the same.

### Fluorescence recovery after photobleaching (FRAP)

FRAP experiments were performed on an LSM900 confocal microscope (Zeiss, Germany) equipped with a full, humidified incubation chamber that was maintained at 37°C with 5.5% CO2. The time series were acquired in confocal mode using two-stage acquisition with the following excitation parameters: 561 nm excitation laser at 0.8% power, 53 μm pinhole size corresponding to 1 AU, scan speed 7, and 3x crop factor corresponding to a pixel dwell time of 3.06 μs. During the first phase, we imaged 25 frames with 0.5 seconds between frames, and during the second phase we imaged 160 frames with 4 seconds between frames, resulting in a total movie length of 657 seconds. For bleaching, a 1μm diameter ROI was selected. Bleaching was performed after frame 8 using both the 488nm and 561nm lasers at 100% power, 53μm pinhole size, and scan speed 7 corresponding to a pixel dwell time of 3.06μs. For each condition, 30 movies were acquired and between 3 and 11 were excluded due to drift.

### ChIPmentation

mESCs were dissociated using TrypLE (Thermofisher #12605010) and washed once in 1X PBS. After counting, cells were divided in 10 million aliquots and resuspended in fresh 1X PBS (1million cells/1ml). For double cross linking 25mM EGS (ethylene glycol bis(succinimidyl succinate); Thermofisher #21565) were added and cells were put in rotation for 30 min at room temperature, followed by addition of 1% formaldehyde (Tousimis #1008A) for 10 min also in rotation at room temperature. Quenching was performed by adding glycine to a final concentration of 0.125 M followed by incubations of 5 min at room temperature in rotation. Fixed cells were washed twice with 5 ml of 1 X PBS containing 0.5% BSA and centrifuged at 3000 rpm for 5 min at 4°C. Pellets were finally resuspended in 500 μl 1X PBS containing 0.5% BSA, transferred to 1.5 ml Eppendorf and centrifuged at 3000rpm for 3 min at 4°C. Supernatant was completely removed, pellets were snap-frozen in liquid nitrogen and stored at -80°C. Fixed cells (10 million) were thawed on ice, resuspended in 350 μl ice cold lysis buffer (10 mM Tris-HCl (pH 8.0), 100 mM NaCl, 1 mM EDTA (pH 8.0), 0.5 mM EGTA (pH 8.0), 0.1% sodium deoxycholate, 0.5% N-lauroysarcosine and 1X protease inhibitors) and lysed for 10 min by rotating at 4° C. Chromatin was sheared using a bioruptor (Diagenode) for 15 minutes (30 sec on, 30 sec off, high output level). 100 μl of cold lysis buffer and 50 μl of 10% Triton X-100 (final concentration of 1) were then added and the samples were centrifuged for 5 min at full speed at 4°C. Supernatant was collected, transferred to a new tube (Protein Low Binding tube) and shearing was continued for another 10 min, then the chromatin was quantified. FLAG M2 Magnetic Beads (Sigma, M8823) were used for FLAG immunoprecipitation. In other cases, SMC3 (abcam, ab9263), CTCF (cell Signaling Technology, 3418) and Rabbit IgG (abcam, ab37415) antibodies were bound to protein A magnetic beads by incubation on a rotator for one hour at room temperature. 10 μl each of antibody was bound to 50 μl of protein-A magnetic beads (Dynabeads) and added to the sonicated chromatin for immunoprecipitation at 4°C overnight. Next day, samples were washed and tagmentation were performed as per the original ChIPmentation protocol ^54^. In short, the beads were washed successively twice in 500 μl cold low-salt wash buffer (20 mM Tris-HCl (pH 7.5), 150 mM NaCl, 2 mM EDTA (pH 8.0), 0.1% SDS, 1% tritonX-100), twice in 500 μl cold LiCl-containing wash buffer (10 mM Tris-HCl (pH 8.0), 250 mM LiCl, 1 mM EDTA (pH 8.0), 1% triton X-100, 0.7% sodium deoxycholate) and twice in 500 μl cold 10 mM cold Tris-Cl (pH 8.0) to remove detergent, salts and EDTA. Subsequently, the beads were resuspended in 25 μl of the freshly prepared tagmentation reaction buffer (10 mM Tris-HCl (pH 8.0), 5 mM MgCl2, 10% dimethylformamide) and 1 μl Tagment DNA Enzyme from the Tagment DNA Enzyme and Buffer Kit (Illumina #20034198) and incubated at 37°C for 10 min in a thermomixer. Following tagmentation, the beads were washed successively twice in 500 μl cold low-salt wash buffer (20 mM Tris-HCl (pH 7.5), 150 mM NaCl, 2 mM EDTA (pH8.0), 0.1% SDS, 1% triton X-100) and twice in 500 μl cold Tris-EDTA-Tween buffer (0.2% tween, 10 mM Tris-HCl (pH 8.0), 1 mM EDTA (pH 8.0)). Chromatin was eluted and de-crosslinked by adding 70 μl of freshly prepared elution buffer (0.5% SDS, 300 mM NaCl, 5 mM EDTA (pH 8.0), 10 mM Tris-HCl (pH 8.0) and 10 ug/ml proteinase K for 1 hour at 55°C 850rpm and overnight at 65°C 850rpm. Next day, the supernatant was collected, transferred to new DNA Low Binding tubes and supplemented with an additional 30 μl of elution buffer. DNA was purified using MinElute Reaction Cleanup Kit (Qiagen #28204) and eluted in 20 ul. Purified DNA (20 μl) was amplified as per the ChIPmentation protocol (Schmidl, Rendeiro et al. 2015) using indexed and non-indexed primers and NEBNext High-Fidelity 2X PCR Master Mix (NEB M0541) in a thermomixer with the following program: 72°C for 5 m; 98°C for 30 s; 14 cycles of 98°C for 10 s, 63°C for 30 s, 72°C for 30 s and a final elongation at 72°C for 1 m. DNA was purified using Agencourt AMPure XP beads (Beckman, A63881) to remove fragments larger than 700 bp as well as the primer dimers. Library quality and quantity were estimatred using Tapestation (Agilent High Sensitivity D1000 ScreenTape #5067-5584 and High Sensitivity D1000 reagents #5067-5585) and quantified by Qubit (Life Technologies Qubit™ 1X dsDNA High Sensitivity (HS) #Q33230). Libraries were then sequenced with the Novaseq6000 Illumina technology according to the standard protocols and with around 200 million 150bp paired-end total per sample.

### Total RNA-seq

mESCs were dissociated using TrypLE, washed in 1X PBS, pelletted and 2.5 million cells were used for extracting RNA with RNeasy plus kit (Qiagen #74134) in each case. RNA quality was checked in Tapestation (Agilent High Sensitivity RNA ScreenTape #5067-5579, High Sensitivity RNA Sample Buffer #5067-5580 and High Sensitivity RNA Ladder #5067-5581) and quantified by Nanodrop. 500 ng were used for Total RNA libraries preparation using Illumina kit (Illumina Stranded Total RNA Prep, Ligation with Ribo-Zero Plus #20040529). Library concentrations were estimated using Tapestation (Agilent High Sensitivity D1000 ScreenTape #5067-5584 and High Sensitivity D1000 reagents #5067-5585) and quantified by Qubit (Life Technologies Qubit™ 1X dsDNA High Sensitivity (HS) #Q33230). Libraries were then sequenced with the Novaseq6000 Illumina technology according to the standard protocols and with around 100 million 150bp paired-end total per sample.

### Hi-C

Hi-C was performed in duplicates using around 1 million cells each. mESCs were dissociated using TrypLE (Thermofishe #12605010), washed once in 1X PBS and resuspended in fresh 1X PBS (1 million cells/ 1 ml). For double cross linking 25 mM EGS (ethylene glycol bis(succinimidyl succinate); Thermofisher #21565) were added and cells were put in rotation for 30 min at room temperature, followed by addition of 1% formaldehyde (Tousimis #1008A) for 10 min also in rotation at room temperature. Quenching was performed by adding glycine to a final concentration of 0.125M followed by incubations of 5 min at room temperature in rotation. Fixed cells were washed twice with 5 ml of 1X PBS containing 0.5% BSA and centrifuged at 3000 rpm for 5 min at 4°C. Pellets were finally resuspended in 500 μl 1X PBS containing 0.5% BSA, transferred to 1.5ml Eppendorf and centrifuged at 3000 rpm for 3 min at 4°C. Supernatant was completely removed, pellets were snap-frozen in liquid nitrogen and stored at -80°C. Samples were subsequently processed using the Arima Hi-C kit (A510008) as per the manufacturer’s protocol and sequenced with the Novaseq6000 Illumina technology according to the standard protocols and with around 600 million 150bp paired-end reads per sample.

### ATAC-seq

We used an improved ATAC-seq protocol (Omni-ATAC) adopted from Corces et al., ^22^ with small adaptations. Briefly, cells in culture were treated with 200 U/ml DNase (Worthington # LS002007) for 30 min at 37°C to remove free-floating DNA and any DNA from dead cells. Cells were then harvested via trypsinization and resuspended in regular medium. After counting, 500,000 cells were collected, resuspended in 1X cold PBS and spin down at 500 g for 5 min at 4°C in a fixed-angle centrifuge. After centrifugation, the supernatant was removed and the pellet resuspended in 500 μl of cold ATAC-seq resuspension buffer 1 (10 mM Tris-HCl pH 7.4, 10 mM NaCl, 3 mM MgCl2, 0.1% NP-40 and 0.1% Tween-20 in water). 50 μl (50.000 cells) were transferred to a new 1.5 ml Eppendorf tube and 0.5 μl l 1% Digitonin (Promega #G9441) was added by pipetting well few times. The cell lysis reaction was incubated on ice for 3 min. After lysis, 1 ml of cold ATAC-seq resuspension buffer 2 (10 mM Tris-HCl pH 7.4, 10 mM NaCl, 3 mM MgCl2 and 0.1% Tween-20 in water) was added. Tube was inverted three times to mix and nuclei were pelleted by centrifugation for 10 min at 500 g at 4°C. Supernatant was very carefully removed (pellet could be almost invisible at this step) and nuclei resuspended in 45 μl of transposition mix (25 μl 2X TD buffer (Illumina #20034198), 16.5 μl 1X PBS, 0.5 μl 1% digitonin, 0.5 μl 10% Tween-20 and 2.5 μl water) by pipetting up and down few times. 5 μl of transposase enzyme (Illumina #20034198) was then added and the transposition reaction was incubated at 37°C for 30 min in a thermomixer with shaking at 1000 rpm. Reaction was cleaned up with Zymo DNA Clean and Concentrator-5 kit (Zymo#11-302C). Transposed DNA fragments were then amplified as described previously in Buenrostro et al., ^55^. Final libraries were purified with with Zymo DNA Clean and Concentrator-5 kit (Zymo#11-302C; Note: better to use two separate kits for pre- and post-amplification clean up), checked by Tapestation (Agilent High Sensitivity D1000 ScreenTape #5067-5584 and High Sensitivity D1000 reagents #5067-5585) and quantified by Qubit (Life Technologies Qubit™ 1X dsDNA High Sensitivity (HS) #Q33230). Libraries were then sequenced with the Novaseq6000 Illumina technology according to the standard protocols and with around 50 million 150bp paired-end reads per sample.

## QUANTIFICATION AND STATISTICAL ANALYSIS

### Fluorescence recovery after photobleaching (FRAP) analysis

FRAP analysis was performed using custom Python scripts. Briefly, movies were loaded into a custom Python GUI (https://github.com/ahansenlab/FRAP-drift-correction-GUI). The bleached nucleus was segmented using manually selected segmentation parameters and was used to estimate the average nuclear intensity and the average background intensity. To correct for cell drift, the ROI positions were manually updated at a number of frames and the ROI position was linearly interpolated between manual updates. Cells with excessive drift were manually excluded and the mean fluorescence intensity of the bleach was taken as the average intensity within the ROI.

To correct for photobleaching, *I_nonbleach_*(*t*) was smoothed using an averaging filter, and a series of correction factors (*C*(*t*)) were computed as the ratio of these values to their average prebleach value as follows:

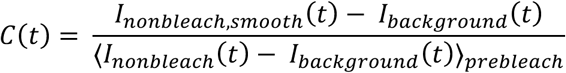

Subsequently, the FRAP curves were normalized according to the following equation:

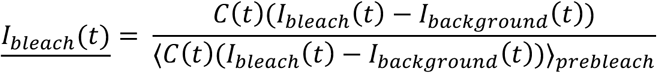

where *I_bleach_*(*t*) is the normalized, background-subtracted, and photobleaching corrected FRAP signal.

Finally, since previous measurements of CTCF’s free diffusion coefficient have indicated a reaction-dominant model is most appropriate, we fit the following, previously demonstrated model (Sprague, Pego et al. 2004, Hansen, Pustova et al. 2017).

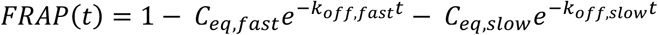

where *k_off,slow_* < *k_off,fast_*.*C_eq,slow_* was taken as the specific bound fraction and the specific residence time was taken as 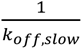. To estimate parameter distributions, we implemented a bootstrapping routine in which a set of 16 movies were randomly resampled from the full dataset 2500 times with replacement. For each sample, the above equation was fit and the parameter values were taken as one data point. The 2500 resampled points were then used as distributions for plotting the specific bound fraction and residence time. For ease of comparison, the specific bound fractions were normalized to the average WT value when plotting.

### ChIP-seq data processing and quality control

Data were processed using Seq-N-Slide pipeline ^56^. Briefly, after trimming for NEXTERA adaptor sequence using trimGalore ^57^. The reads were aligned to mm10 genome with Bowtie2 ^58^. Ambiguous reads were filtered to use uniquely mapped reads in the downstream analysis. PCR duplicates were removed using Sambamba ^59^). Narrow peaks were called using MACS2 ^60^ in pair-end mode and with IgG as control for SMC3 and CTCF ChIP-seq or WT untreated condition (in which FLAG CTCF is not expressed) for CTCF FLAG to control for unspecific binding, Peaks overlapping ENCODE blacklisted regions were filtered out ^61^. These procedures allowed us to minimize the false positive calls, as reflected by the high percentage of called peaks containing CTCF consensus motifs as detected by MEME ^21^ with 63% for WT. To ensure comparability between conditions, FLAG and SMC3 chip-seq samples were down-sampled to the same number of aligned (100 millions reads per replicates), except for 2 samples R377H-rep1 and R339Q-rep which were under-sequenced (with 68 and 90 millions reads, respectively) and good quality reads before peak calling using samtools ^62^. Bigwigs were obtained for visualization on individual as well as merged bam files using Deeptools ^63^. For CTCF FLAG visualization, the differential signals were generated after subtracting non-specific signal present in the WT untreated condition. Heatmaps and average profiles were performed on merged bigwig files using Deeptools.

### RNA-seq data processing and quality control

Data were processed using Seq-N-Slide pipeline ^56^. Briefly, after trimming for the Illumina adaptor and the T overhang as per Illumina recommendation using TrimGalore ^57^, reads were aligned against the mouse reference genome (mm10) using the STAR ^64^ aligner and differentially expressed genes were called using DESeq2 ^65^ with an adjusted p-value of 0.05 and a fold change cutoff of 1.5. Heatmap was performed using pheatmap R package ^66^ and GSEA using fgsea R package ^67^ and KEGG genesets ^32^. Developmental germ layer gene sets were downloaded from Hutchins *et a*l.,^68^. Radarplots were generated using the R package fsmb after averaging the expression level of the genes of each germ layers and of the 2 replicates. CTCF-associated neurodevelopmental disorder (NDD) genes were retrieved from Konrad *et al.* ^9^, while genes associated with developmental disorders were downloaded from DECIPHER (https://www.deciphergenomics.org/ddd) and imprinted genes from geneimprint (https://www.geneimprint.com/). Enrichment analyses for those sets were performed using logistic regressions. To compare the change in gene expression during the exit from pluripotency between WT and mutant (longitudinal analysis), linear regressions on the difference of log2 between LIF versus no LIF condition were performed.

### ATAC-seq data processing and quality control

Data were processed using Seq-N-Slide pipeline ^56^. Briefly, Reads were aligned to mm10 genome with Bowtie2 ^58^. Ambiguous reads were filtered to use uniquely mapped reads in the downstream analysis. PCR duplicates were removed using Sambamba ^59^. Mitochondrial contamination was assessed using samtools ^62^. All the samples showed less than 0.6% of mitochondrial DNA, Peaks were called using MACS2 ^60^and peaks overlapping ENCODE blacklisted regions were filtered out^61^. Bigwigs were obtained for visualization on individual as well as merged bam files using Deeptools ^63^. Heatmaps and average profiles were performed on merged bigwig files using Deeptools.

### Hi-C Processing and Quality Control

HiC-Pro ^69^ was used to align and filter the Hi-C data. To generate Hi-C filtered contact matrices, the Hi-C reads were aligned against the mouse reference genome (mm10) by bowtie2 (version 2.3.1) using local mode for the second step of HiC-Pro alignment (--very-sensitive -L 20 --score-min G,20,8 --local) to ignore the molecular barcode UMI sequences and dark bases introduced by Arima library prep kit at the 5’ end of the read. Singleton, ambiguous and duplicated reads were filtered out. Ligation sites for Arima HiC kit were set up using GATCGATC,GANTGATC,GANTANTC,GATCANTC. Low mapping quality (MAPQ<30), self circle, dangling ends and re-ligation reads were filtered out through the HiC-pro pipeline. Valid pairs represented more than 80% for all the samples. Samples were downscaled to the same number of good quality uniquely aligned reads using HiCExplorer ^70^. Of note, we did not use the number of valid pairs to downscale the samples to not overcorrect some conditions such as IAA and mutant H455R which were expected to show a dramatic biological decrease in valid interactions.

Downscaled interaction matrices were generated separately for each replicate and merged for visualization and downprocessing as mcool files using cooler ^71^. Samples were balanced using the ICE correction method ^72^ with HiCExplorer ^70^. HiC maps were generated from the merged downscaled and corrected matrices using GENOVA and HiCExplorer ^44,70^. RCP (relative contact probabilities) and insulation score were generated using GENOVA ^44^. After log-log transformation and Loess smoothing using R, the first derivative (P’max) was estimated as described in Gassler et al., ^41^.

### CTCF peak motif analysis

Motifs were called using MEME ^21^ with a 1^st^ order background model on CTCF peaks called using MACS2 after separating the peaks into 3 groups for each pairwise comparison between WT and mutant CTCF: WT only, common and mutant only peaks. Peak intersection sets were generated using the intervene package ^73^and motifs were searched within 100 bp of each peak summit.

### AlphaFold3 prediction of the DNA-CTCF complexes in the mutants

As a quality control, to compare the prediction to experimental data, we downloaded the structure deposited by Yang *et al*. ^13^ from PDB for WT CTCF ZF1-ZF7 (accession number PDB 8SSS, with oligo CCAGCAGGGGGCGCTAGTGAG and CTCACTAGCGCCCCCTGCTGG), K365T CTCF ZF1-ZF7 (accession number PDB 8SST with the same oligos as WT), WT CTCF ZF3-ZF11 (accession number PDB 8SSQ with oligo GTGCAGTACCACATTTAACCAGCAGGGGGCGCTAA and TTAGCGCCCCCTGCTGGTTAAATGTGGTACTGCAC), K365T CTCF ZF3-ZF11 (accession number PDB 8SSR with oligo with the same oligos as WT). For the Alphafold3 predictions we used the same DNA sequences and the CTCF ZF1-11 positions obtained from Yang *et al* ^13^. AlphaFold3 predictions ^53^ were also carried out for WT and K365T CTCF ZF3-ZF11 and CTCF ZF1-7 using the same DNA and CTCF sequence to compare the prediction to the experimental data. We, then, predicted the CTCF (ZF1-11)-DNA complex for WT and each mutant CTCF using the concatenated DNA sequences from Yang *et al* ^13^ (GTGCAGTACCACATTTAACCAGCAGGGGGCGCTAGTGAG and CTCACTAGCGCCCCCTGCTGGTTAAATGTGGTACTGCAC). The predicted protein-DNA structures were visualized in PyMOL 3.1.0. The hydrogen bonds between DNA and CTCF were identified using PyMOL dist function with a cutoff of 5 Å. The root mean square deviation (RMSD) between the structures was calculated using PyMOL 3.1.0.

### Stratified analysis at enhancers and promoters

H3K27ac peaks from 120/Ola mESCs were downloaded from ENCODE (ENCFF519QMV). CTCF, cohesin and ATAC-seq peaks were classified as active promoter sites when they overlap with an H3K27ac peak and are located in a gene promoter region, defined as 2kb upstream and 1 kb downstream gene TSSs. Active enhancer sites when they overlap with an H3K27ac peak and are located outside gene promoter regions.

### Multivariate CTCF/SMC3 overlap and insulation score analysis

To test the independent of CTCF and Accessibility on SMC3 overlap, CTCF ChIP-seq and ATAC-seq signal strengths were categorized into weak, intermediate and strong signal using the inter-quartile values of the signal across WT and mutant samples. Weak binding or accessibility was defined as sites with signals inferior to the 1^st^ quartile, intermediate binding or accessibility as sites with signals between the 2^nd^ and 3^rd^ quartiles and strong binding or accessibility as sites with signals superior to the 3^rd^ quartile. For SMC3 overlap, a multivariate logistic model testing the effect of CTCF strength, ATAC strength and their interaction was performed in WT mESCs after averaging the replicates. The log2 odd ratios were computed to estimate the enrichment in SMC3. For insulation score a mixed multivariate model testing the effect of CTCF strength, ATAC strength and their interaction was performed across sample. For both models the marginal effects and 95th confidence intervals were computed. These analyses were performed using STATA v17.

### ATAC-seq footprinting analysis

Footprinting analysis was performed on ATAC-seq signal using TOBIAS ^35^. Briefly, ATAC-seq signals were first corrected for Tn5 cutting sequence bias using the ATACorrect function and footprinting score calculated using FootprintScores at peaks detected by MACS. Finally differential footprinting estimated pariwise between WT and mutant CTCF using BINDetect and HOCOMOCO v11 core motif probability weight matrices ^74^. Peaks were annotated for gene promoter using UROPA ^75^ and GENCODE version M25 ^76^. Gene promoters were defined as regions within 2kb upstream and 1 kb downstream the gene transcription starting site.

### Loop calling and annotation

Significant loops (q-value<0.05) were called on the downscaled and corrected interaction matrices using FitHIC at 10kb resolution ^77^ and annotated for CTCF peaks and gene TSS using pairToBed ^78^.

### Differential aggregated peak and TAD analyses

Aggregated peak and TAD analysis on downscaled and corrected interaction matrices was performed using GENOVA at 10kb resolution ^44^. For aggregated peaks analyses, the union of loops called by FitHIC ^77^ in WT and mutant CTCF for each pairwise comparison was used to generate and compare aggregated interaction at loop anchors. For aggregated TAD analyses, the insulation score was first calculated to identify TAD and TAD boundaries. The union of TADs called by GENOVA in WT and mutant CTCF was used to generate and compare aggregated intra and inter-TAD interactions.

### Differential loop analysis

Differential loops (adjusted p-value<0.05) were called for each pairwise comparison between WT and mutant CTCF using MultiHiCCompare which performed a cross-sample normalization before testing for differential loops using an exact test ^79^. Enrichment for differentially expressed genes among differential loops was tested using logistic regression after annotating loop anchors with gene TSS.

### Direct vs indirect effect analysis

Direct effect was defined as differential TF binding with a predicted differential CTCF binding identified using TOBIAS ^35^ or ChIP-seq peak at the gene promoter and/or differential loops identified using MultiHiCCompare ^79^ with a CTCF peak at either anchor. Gene promoters were defined as regions spanning 2 kb upstream and 1 kb downstream gene TSSs. Overlap between gene promoters and differential loop were performed using pairToBed ^78^.

## SUPPLEMENTAL INFORMATION

### Excel table title and legends

**Table S1:** Number and percentage of SMC3 peaks overlapping a CTCF or ATAC-seq peak, stratified by CTCF and ATAC signal strength. This table is related to Figure 4B.

**Table S2:** Differentially expressed genes (DEGs) identified using DE-seq. DEGs were defined as adjusted genes with p-value<0.05 and absolute log2 fold change ≥ log2(1.5) compared to WT. This table is related to **Figure 5A**.

**Table S3:** CTCF mutations altered cell differentiation. Tab A, Expression of germ layer genes (TPM) in LIF versus no LIF condition in WT and the mutant. Tab B, Differentially expressed genes during in LIF condition in mutant compared to WT. Tab C, Differentially expressed genes during in no LIF condition in mutant compared to WT. Tab D, Differentially expressed genes during the exit from pluripotency (LIF to no LIF) in mutant compared to WT. This table is related to **Figure 5C**.

**Table S4:** CTCF mutations altered TF binding. Tab A, Differentially bound transcription factors predicted by footprinting analyses. This tab is related to **Figure 5D**. Tab B, Enrichment of the target genes of differentially bound TFs among the DEGs. This tab is related to **Figure 5E**. Tab C, List of the target genes of differentially bound TFs with differential expression. This tab is related to **Figure 5E**.

## SUPPLEMENTAL FIGURES

**Figure S1.**
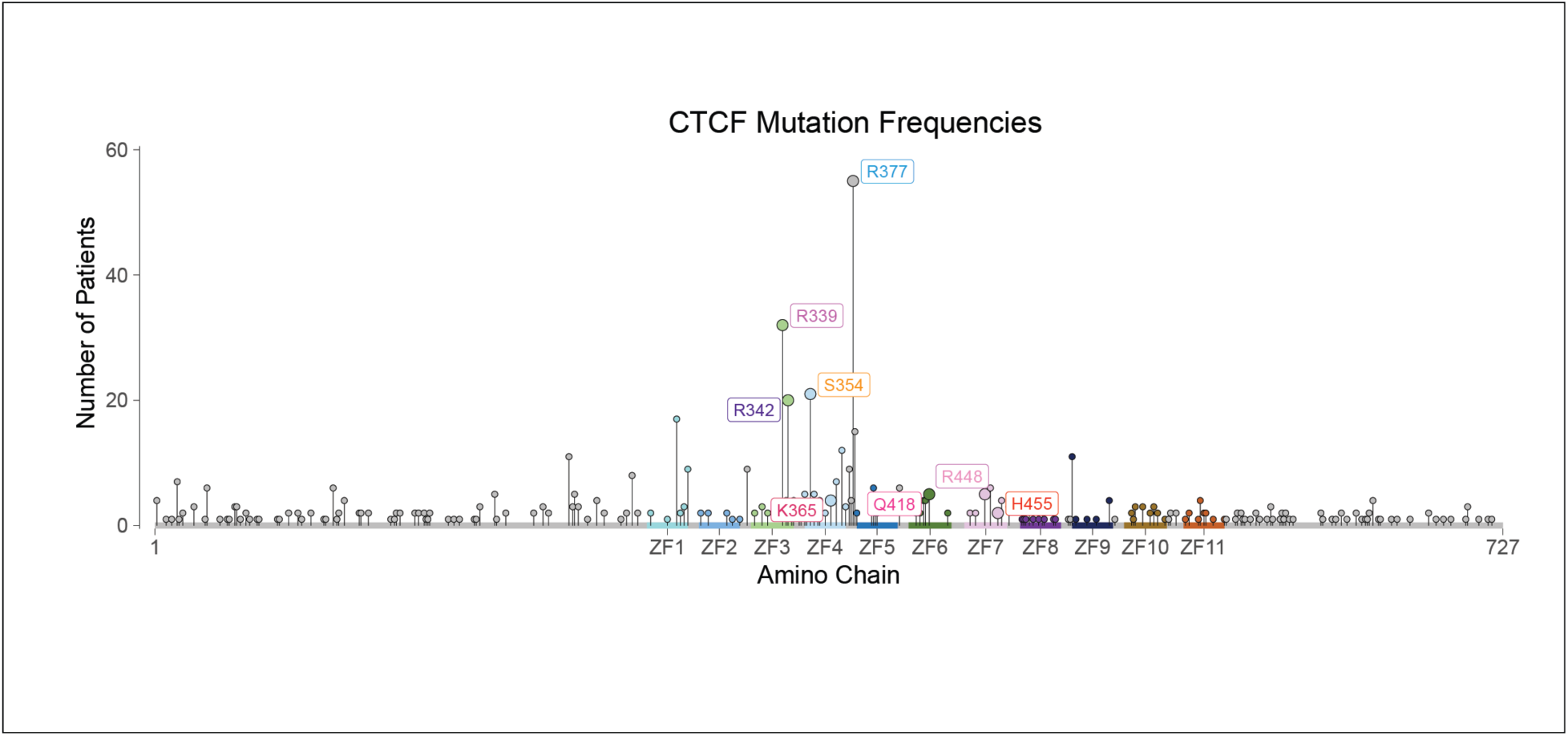
Prevalence of missense CTCF mutations on ZF3-ZF7 in human cancers. Schematic representation of CTCF showing the location and frequency of each mutation under investigation. This figure relates to **Figure 1**.

**Figure S2:**
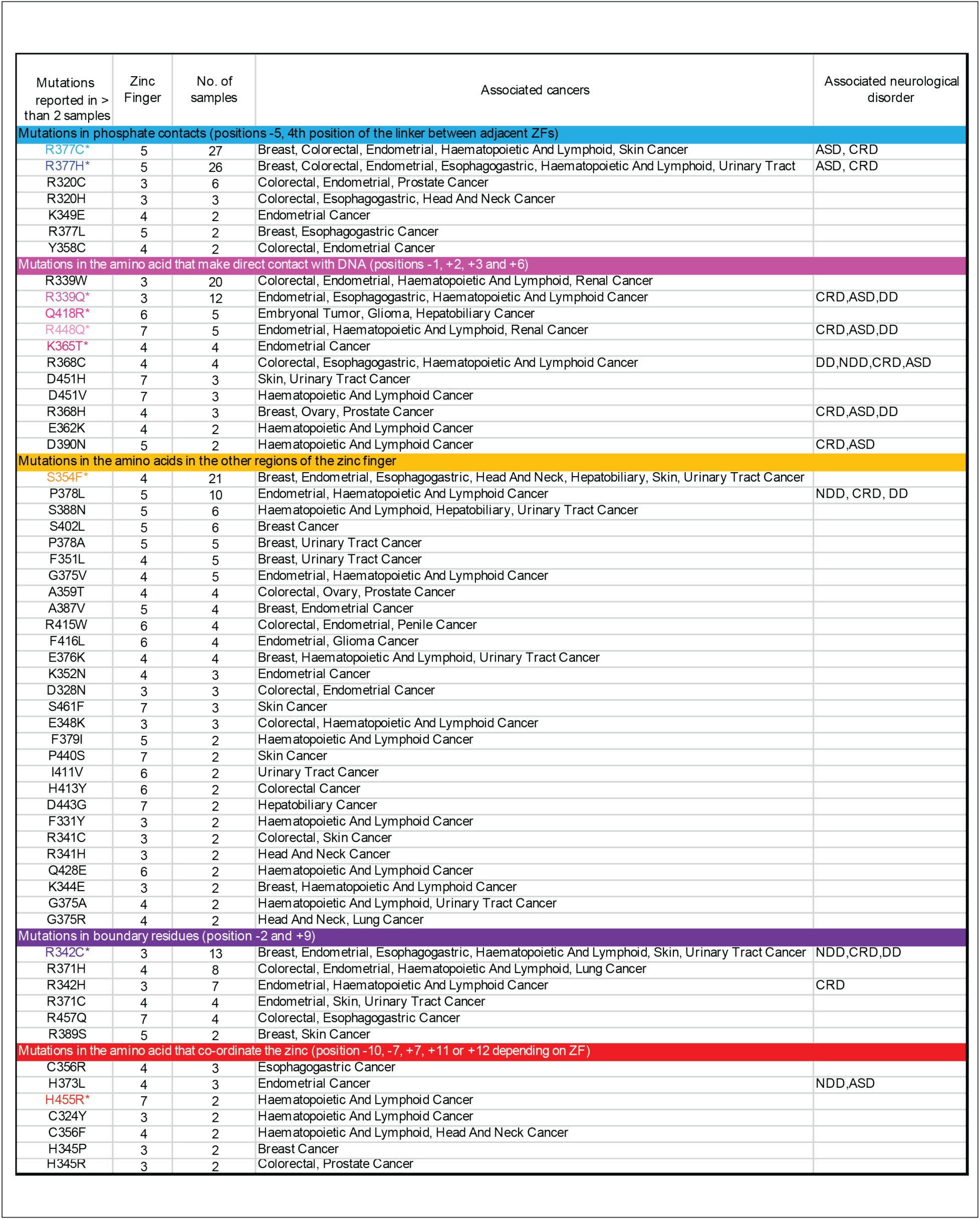
Table of CTCF mutations. Table showing CTCF mutations identified from cBioPortal [S1] and COSMIC [S2]. We selected a subset of mutations that occur at high frequency in cancer patients, focusing on those found in ZFs that make contact with the 12-15bp consensus CTCF binding motif. These are highlighted with an asterisk and color coded as in **Figure 1D**. Mutations analyzed included those found in **(i)** amino acids that make base-specifying contacts with the alpha helix of DNA (at locations -2, 2, 3 and 6 on the ZF), **(ii)** residues that coordinate the zinc ion which are essential for providing stability to ensure the proper folding of the ZF domain, **(iii)** two other classes of amino acid residues in the ZFs: boundary residues (that are important for interactions between ZFs), and residues that contact the sugar phosphate backbone of DNA, and **(iv)** other highly mutated residues in the region. The table identifies which zinc finger each mutation is found in, the number of patients it was identified in and the cancers each is associated with. Mutations were annotated for neurodevelopmental or neurological disorders using the CTCF variant catalog published in Price et al [S3]. ASD, autism spectrum disorder; CRD, CTCF related neurodevelopmental disorder; DD developmental disorder; NDD, neurodevelopmental disorder. This figure relates to **Figure 1**.

**Figure S3:**
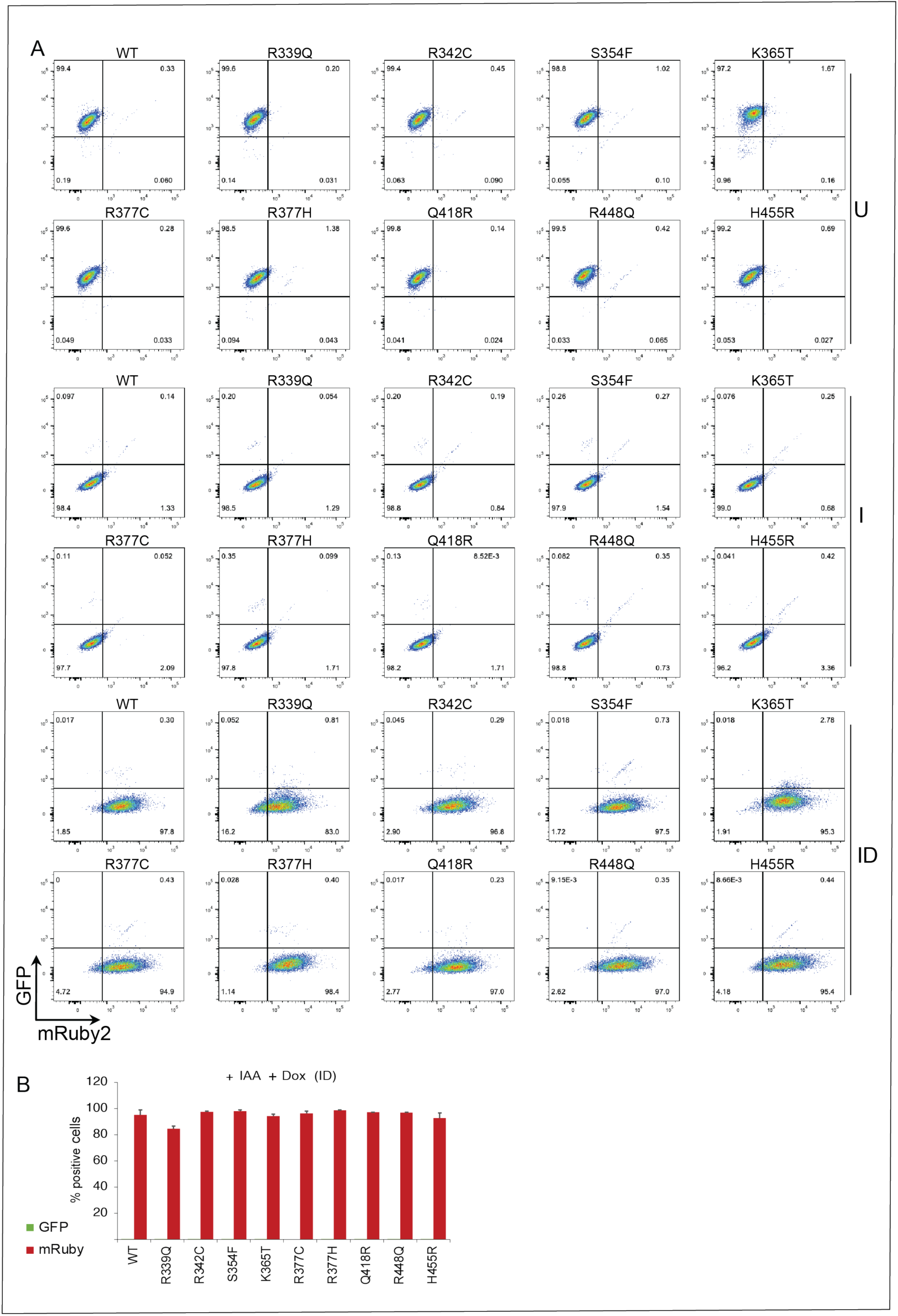
Flow cytometry analysis of mutant expression. (**A**) Flow cytometry showing the level of GFP (endogenous CTCF) and mRuby (transgenic CTCF) in the IAA and Dox treated cells (ID). (**B**) Bar graph showing the percentage of GFP (endogenous CTCF) and mRuby2 (transgene WT and mutant CTCF) positive cells after treatment with IAA and Dox (ID). The presence of the IAA leads to degradation of GFP labelled endogenous CTCF. The error bars represent the standard deviation between the 2 replicates. This Figure relates to **Figure 1**.

**Figure S4:**
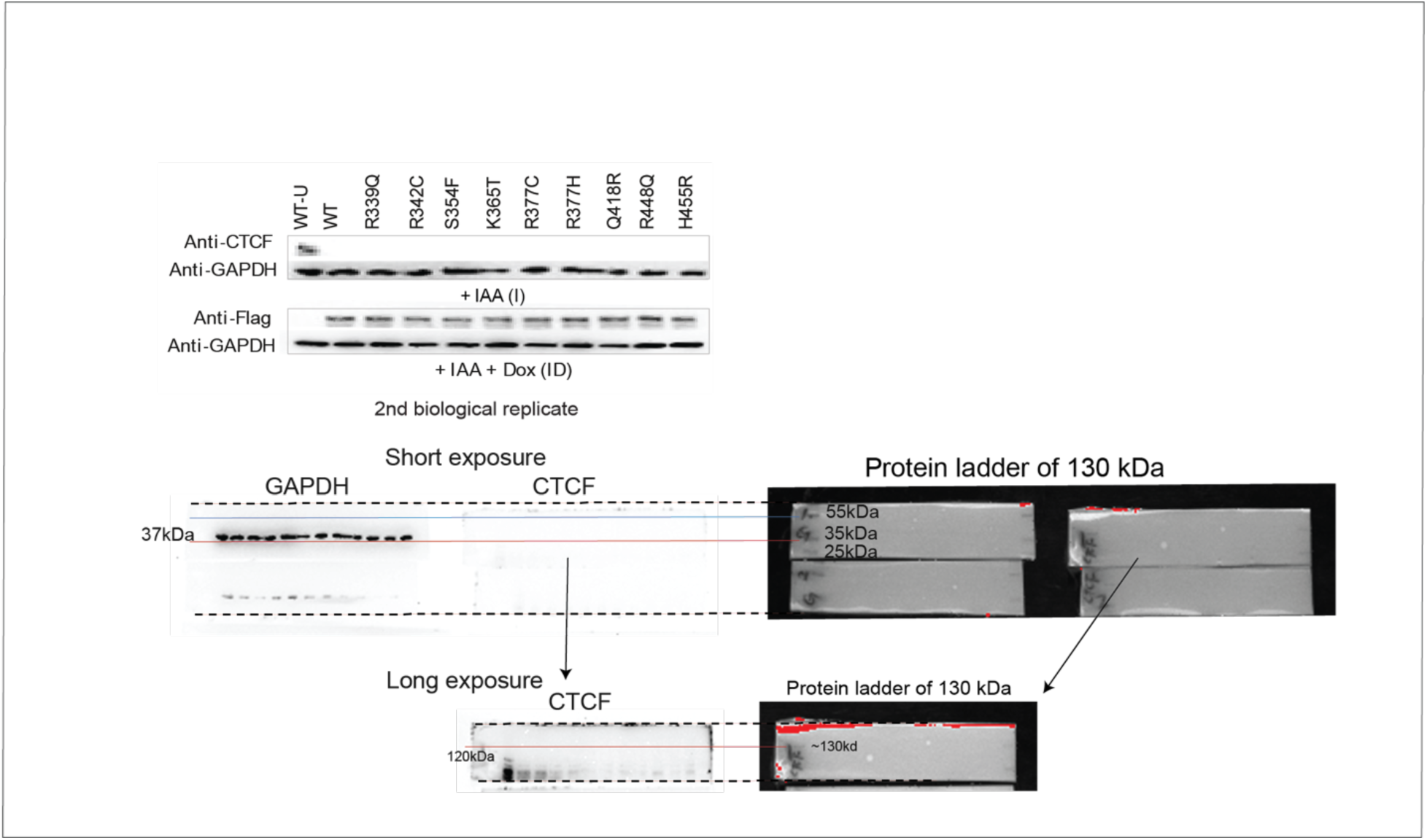
Western Blot of endogenous and transgenic CTCF. Western blot showing degradation of endogenous CTCF as detected with an antibody to CTCF, and the induction of transgenic CTCF as detected using an antibody to FLAG. The first replicate is shown in **Figure 1**. The uncropped pictures of the WB gels are shown below. This figure relates to **Figure 1**.

**Figure S5.**
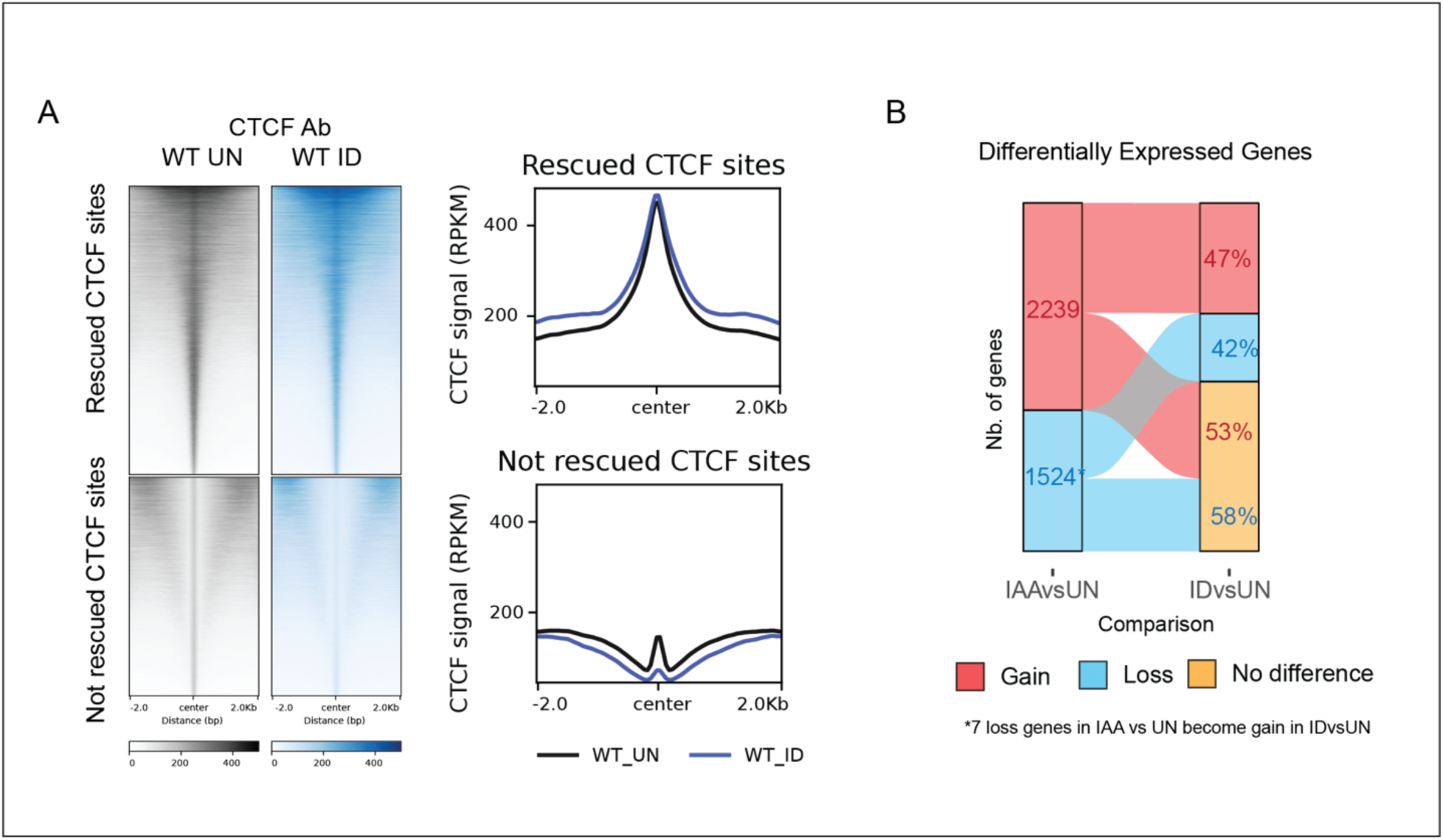
Rescue of CTCF binding and gene expression by the transgene WT CTCF. (**A**) Heatmap and profiles showing the rescued CTCF binding sites in WT-ID compared to WT-UN. (**B**) Alluvial plot showing the number of DEGs in IAA versus WT untreated cells and WT-ID versus WT untreated fells. The plot shows the percentage of genes that return to baseline in the WT-ID condition (no significant expression level changes). DEGs were defined as fold change>1.5 and adjusted p-value<0.05. This figure relates to **Figure 1**.

**Figure S6:**
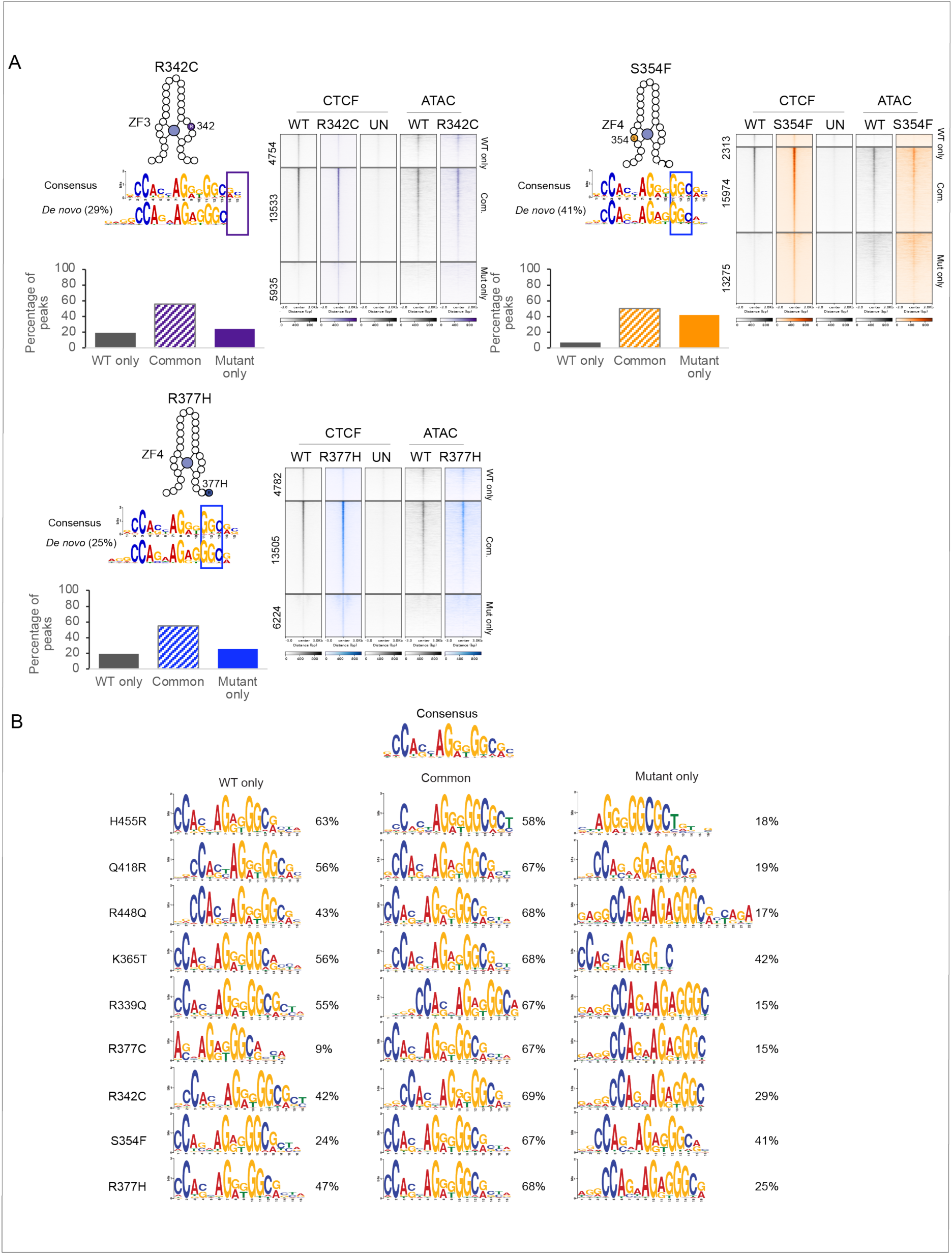
CTCF mutations have unique chromatin binding profiles. **(A**) Scheme showing the locations of the different CTCF mutations within the ZF. The consensus CTCF motif highlights the triplet to which each mutant zinc finger binds (top) as well as the motif identified by MEME for the *de novo* mutant only binding sites (bottom). Each graph shows the percentage of WT and mutant CTCF binding at WT only, common and mutant only sites and heatmaps show the profile of CTCF binding and ATAC-seq signal for WT only, common and mutant only sites. The remaining mutants are shown in **Figure 2**. (**B**) Most prevalent motifs found at WT only, common and mutant only sites. The motifs were identified using MEME de novo motif search. The percentages correspond to the percentage of peaks containing the motif. This figure relates to **Figure 2**.

**Figure S7.**
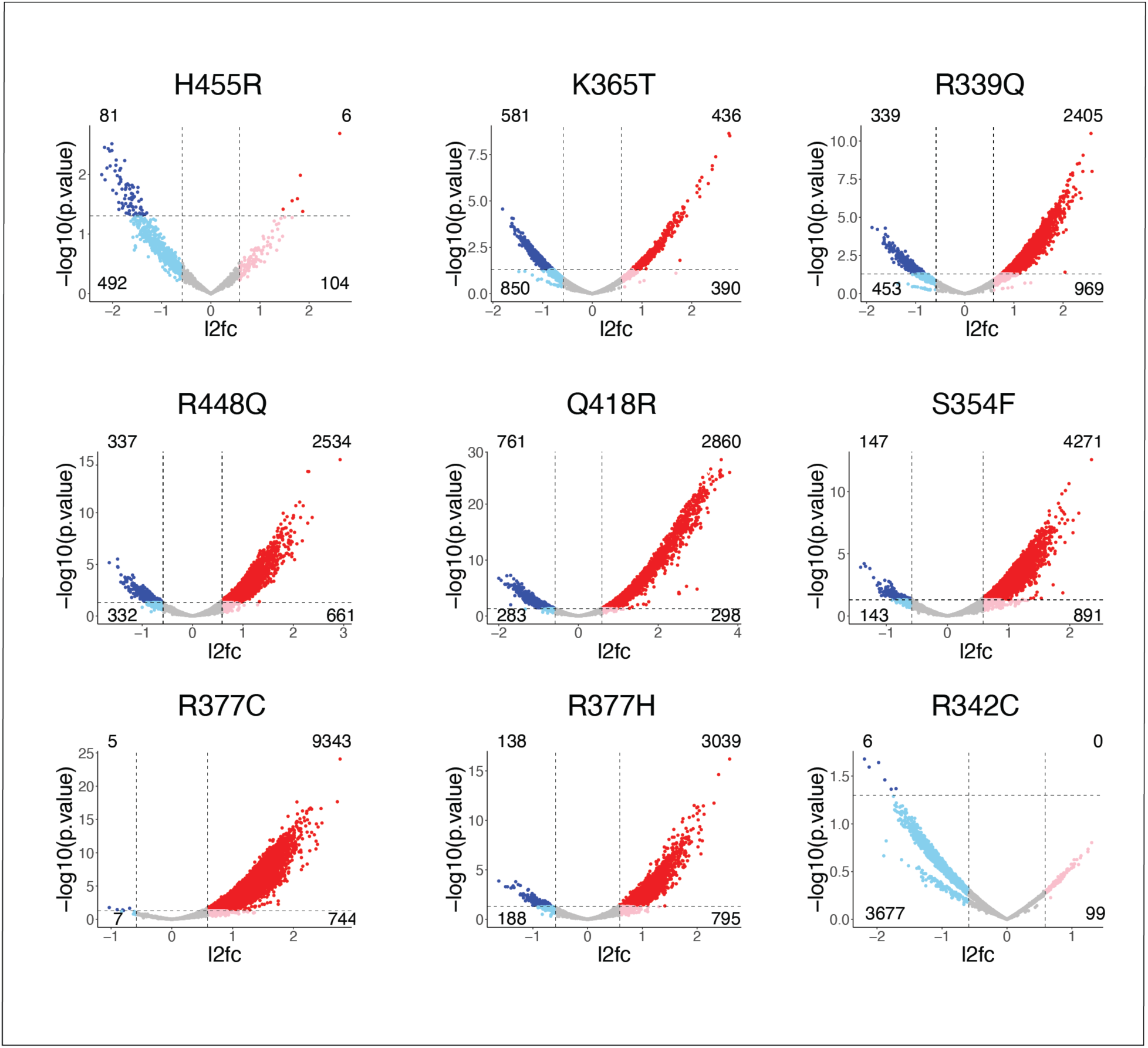
Differential CTCF binding in mutant versus WT-ID. Volcano plots showing significant differences in WT and mutant transgenic CTCF binding (p<0.05 and fold change > 1.5 (red) or fold change < 1.5 (blue). The subthreshold of differentially bound sites are shown in light blue and pink. The p-values were computed using DiffBind. This figure relates to **Figure 2**.

**Figure S8:**
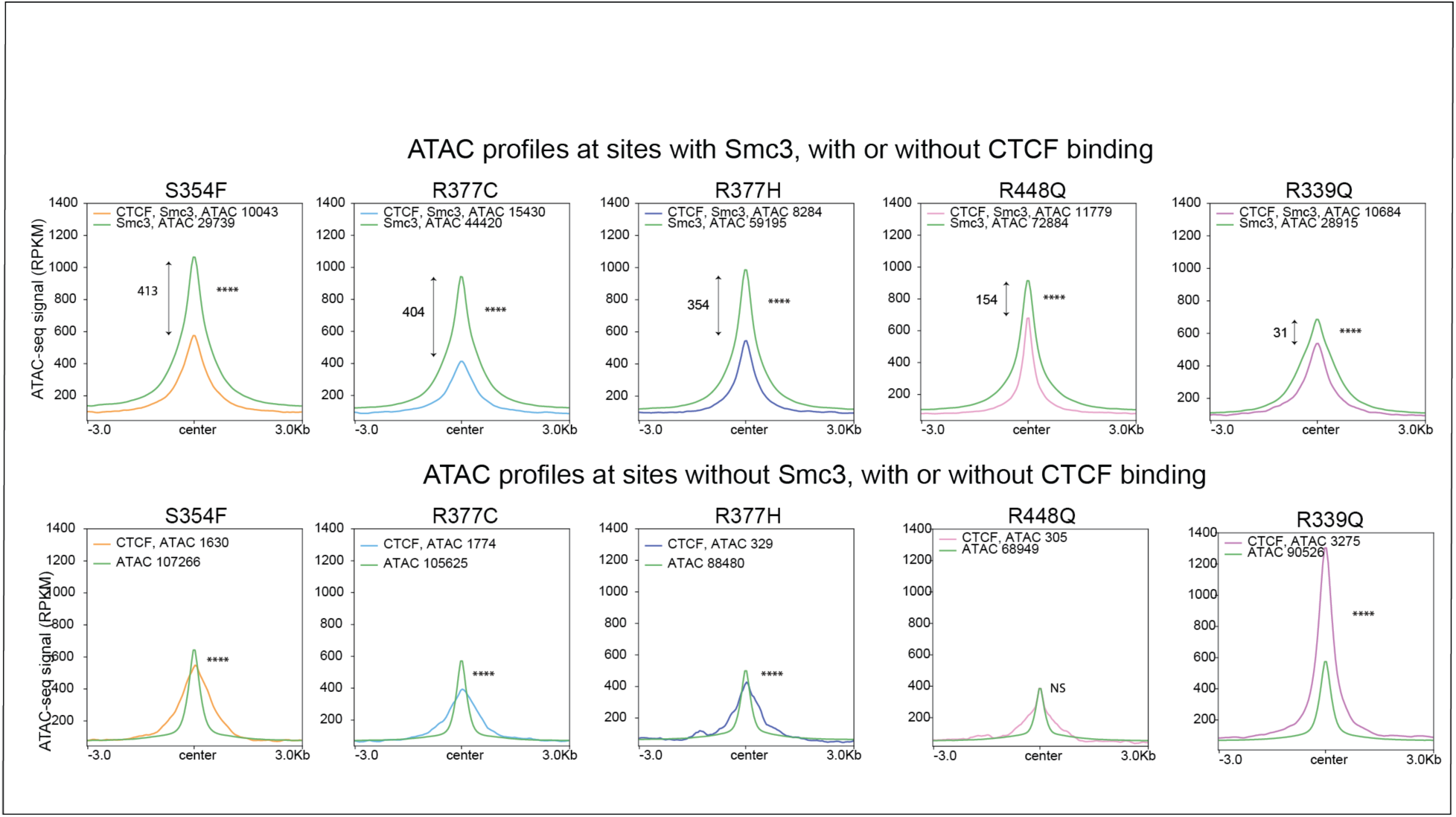
At CTCF WT and mutant sites, accessibility is more or less reduced in a cohesin-dependent manner. Profiles of ATAC-seq in WT and mutant CTCF expressing cells are shown for sites, with (color-coded for each mutant) or without (green) CTCF binding in the presence or absence of SMC3. The remaining mutants are shown in **Figure 2B**. Wilcoxon p-values were coded as follow: * 5×10^-2^-5×10^-3^, ** 5×10^-3^-5×10^-4^, *** 5×10^-4^-5×10^-5^, ****<5×10^-5^. This figure relates to **Figure 2**.

**Figure S9:**
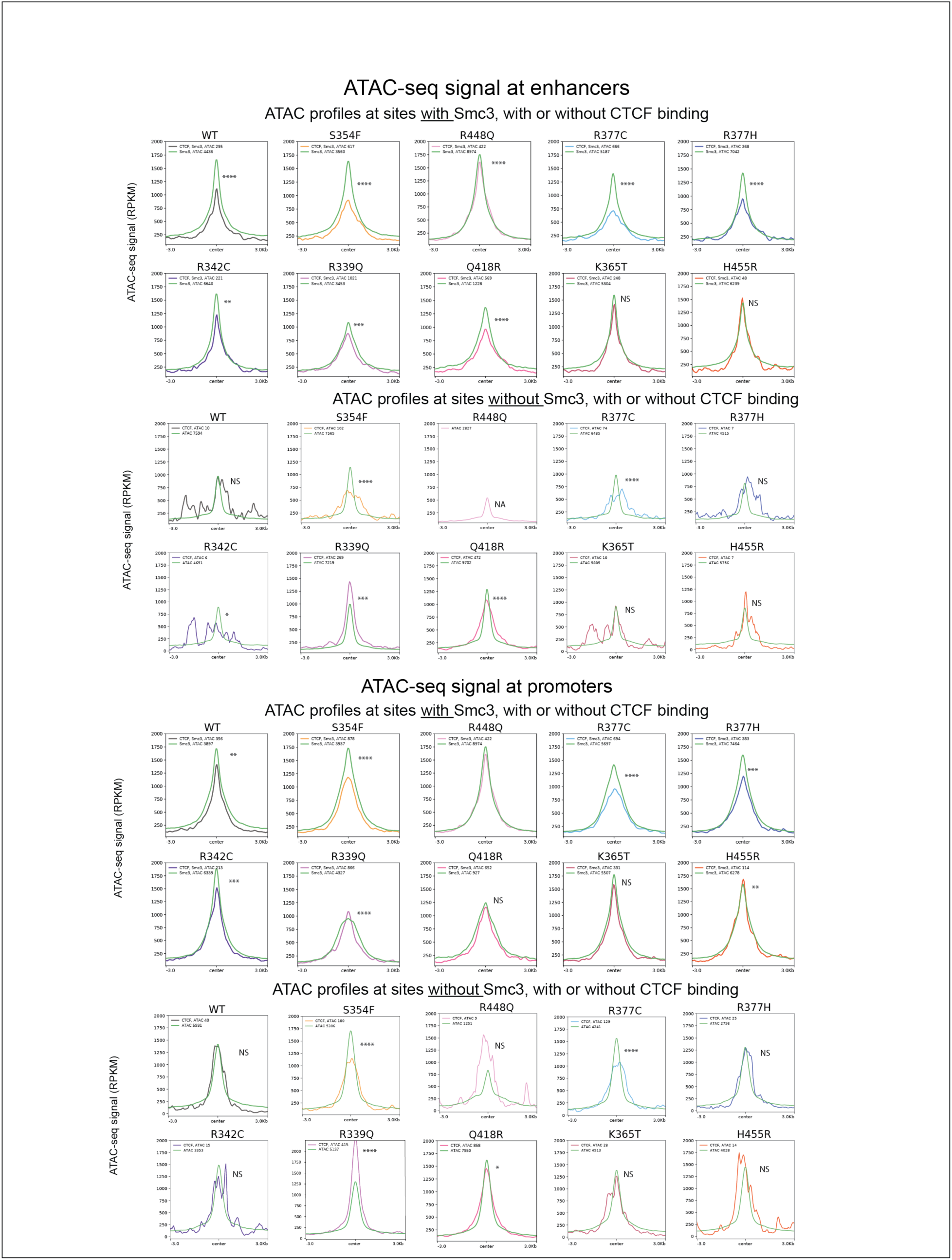
At CTCF WT and mutant sites overlapping gene promoters and enhancers, accessibility is more or less reduced in a cohesin-dependent manner. Profiles of ATAC-seq in WT and mutant CTCF expressing cells are shown for sites, with (color-coded for each mutant) or without (green) CTCF binding in the presence or absence of SMC3 at gene enhancers (top) and gene promoters (bottom). In the absence of SMC3, for WT and most of the mutants, very few accessible CTCF peaks were observed at enhancers or promoters. Those data should be interpreted with caution. Wilcoxon p-values were coded as follow: * 5×10^-2^-5×10^-3^, ** 5×10^-3^-5×10^-4^, *** 5×10^-4^-5×10^-5^, **** <5×10^-5^. This figure relates to **Figure 2**.

**Figure S10.**
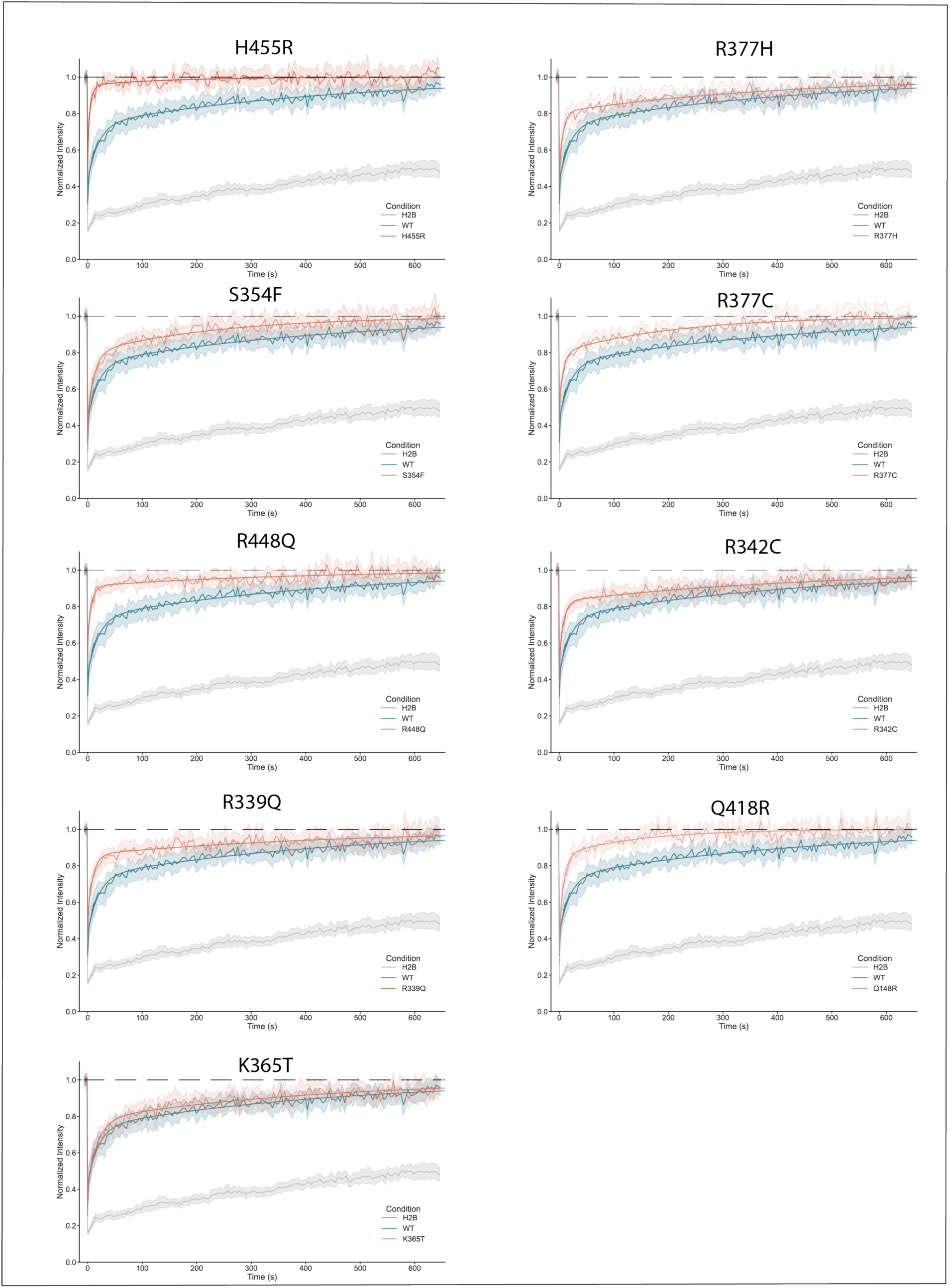
Plots of FRAP dynamics for WT and mutant mRuby-CTCF for each pairwise comparison between mutant and WT. The scatterplots show the average recovery across all movies analyzed, and the outlines give the 95% CI. The bold line shows the fitted model. This figure is related to **Figure 3**.

**Figure S11:**
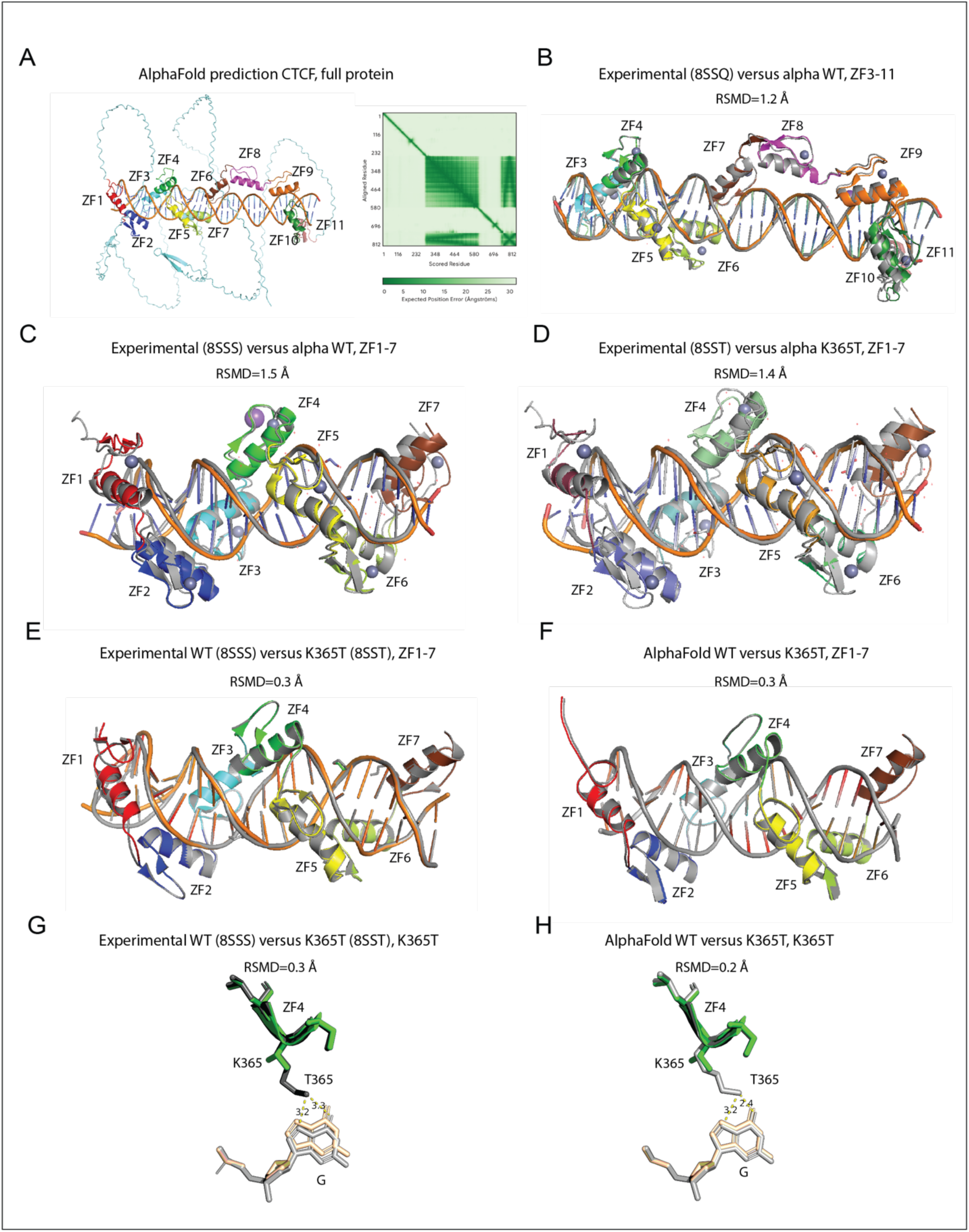
Comparison between the prediction the CTCF-DNA complex and the experimental structure. (**A-H**) Comparison of Alphafold3 prediction versus experimental crystal structure of WT and K365T-DNA complex. (**A**) Prediction of the full CTCF structure. On the right the pLDDT matrix shows a high confidence prediction of CTCF ZF1-11 and its positions relative to the DNA double helix. (**B**) Comparison between the experimental and predicted DNA-WT CTCF ZF3-11 complex. (**C**) Comparison between the experimental and predicted DNA-WT CTCF ZF1-7 structure. (**D**) Comparison between the experimental and predicted DNA-K365T mutant CTCF ZF1-7 structure. For **B-D**, the experimental structures are shown in grey and the prediction in color. Each ZF is color-coded. (**E**) Comparison of the DNA-CTCF complex between K365T mutant and WT from the experimental data and (**F**) between K365T mutant and WT from the AlphaFold3 prediction. For **E** and **F**, the WT structures are shown in grey and the mutant in color. (**G**) Comparison of the DNA-CTCF structure between the K365T mutant and WT showing the hydrogen bond between the mutated or WT residue and interacting DNA nucleotide from the experimental data and (**H**) from the prediction. The experimental crystal structures of the DNA-WT and DNA-K365T complexes were generated in Yang et al ^29^. This figure is related to **Figure 3**.

**Figure S12:**
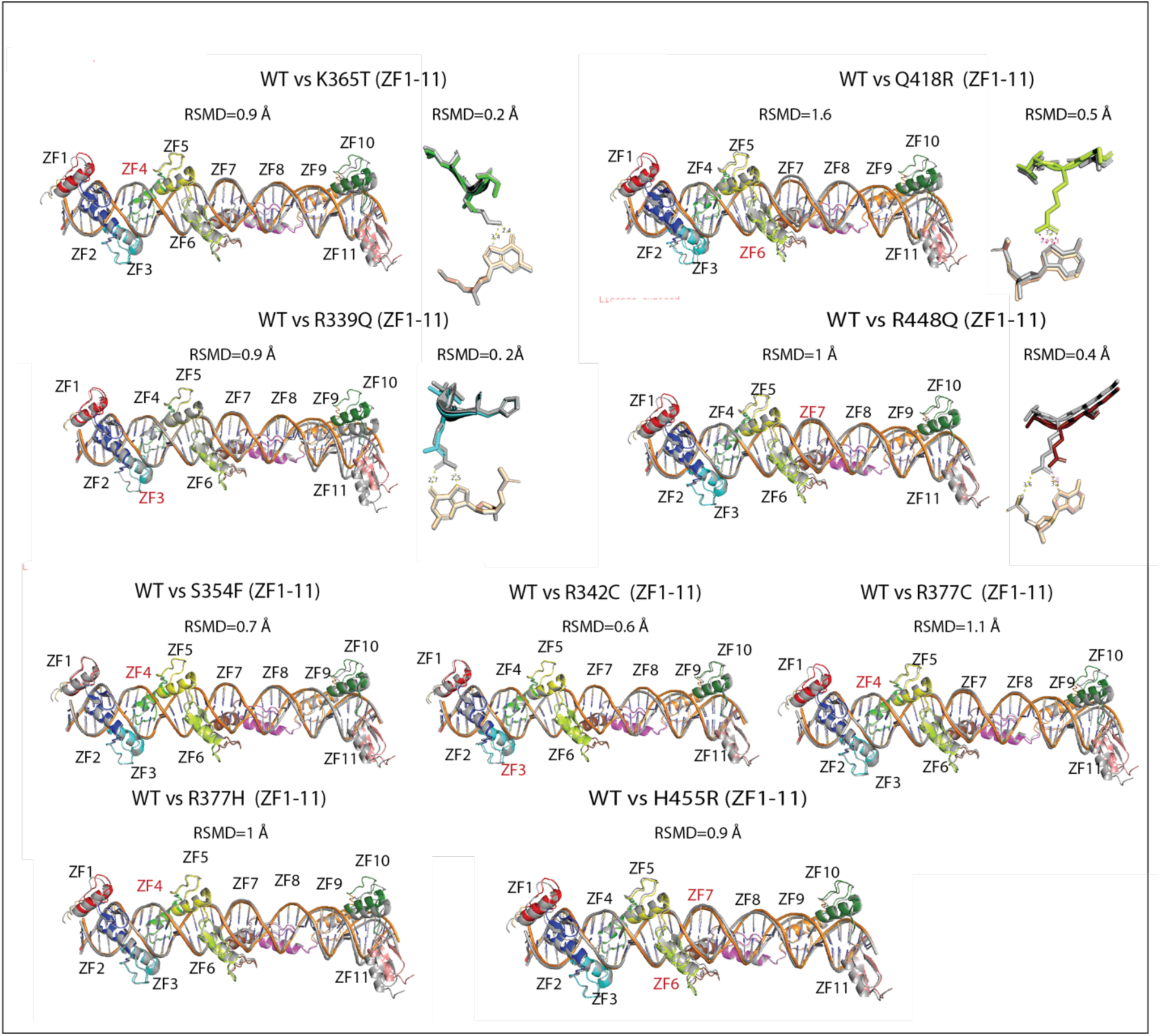
Comparison of the predicted DNA-CTCF complexes between WT and CTCF mutants. Comparison of the DNA-CTCF (ZF1-11) complexes between WT and each mutant CTCF predicted by AlphaFold3. For mutated residues making direct contact with the DNA, the Zoom-in shows the hydrogen bond between the mutated residue and interacting DNA nucleotide compared to WT. The WT structures are shown in grey and the mutant in color. Each ZF is color-coded. The ZFs bearing the mutations are labelled in red. This figure is related to **Figure 3**.

**Figure S13:**
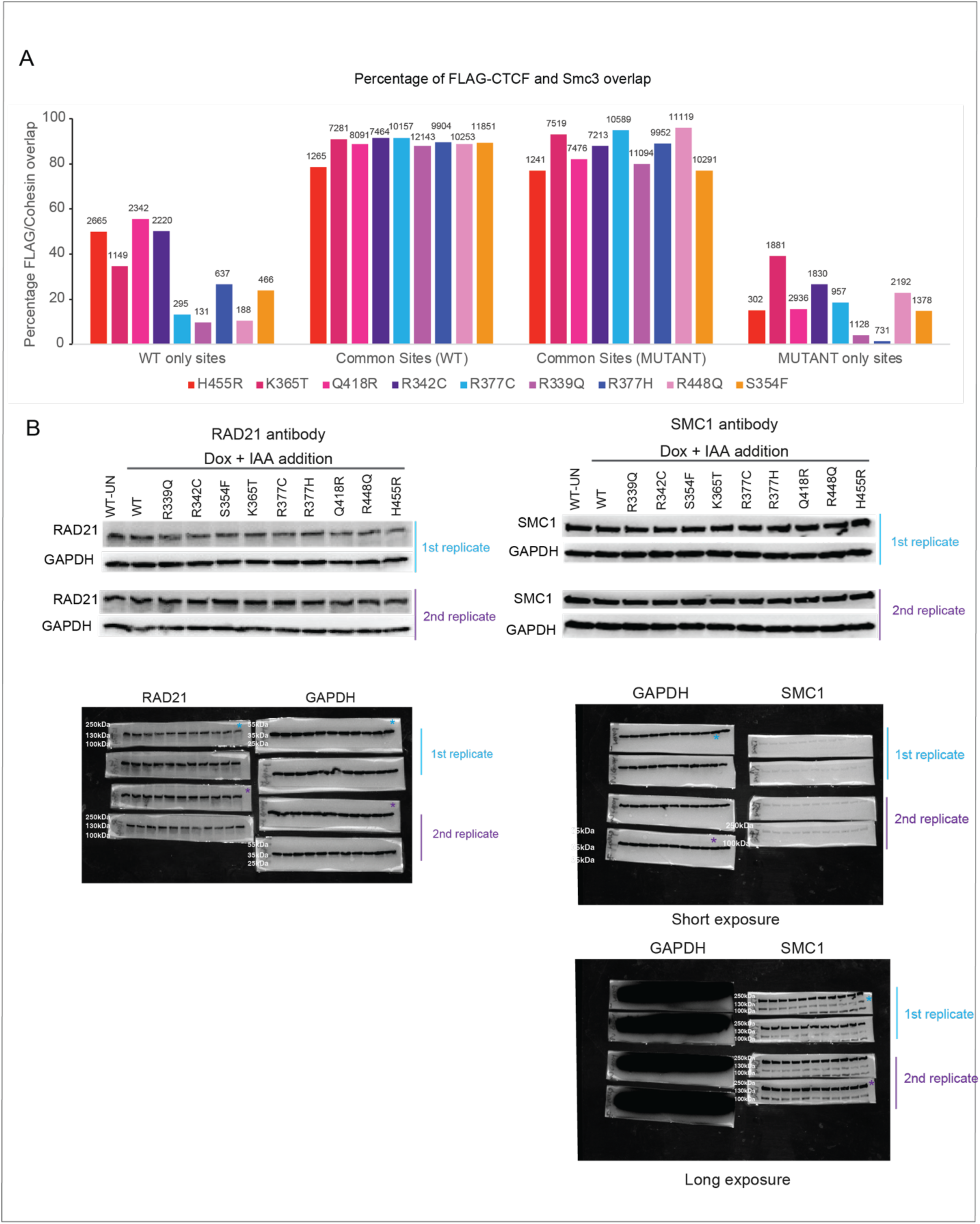
The ability of the CTCF mutants to block cohesin is mostly retained at common sites. Each mutation uniquely. **(A)** Bar graphs showing the percentage of CTCF-cohesin overlap for each mutant at WT only, common and mutant only sites. For common sites, the CTCF peaks were overlapped with either the WT or mutant SMC3 peaks. (**B**) Western blot showing the protein levels of 2 components of the cohesin complex (SMC1 and RAD21) in WT untreated (UN), IAA (CTCF degraded) and WT and CTCF mutant ID treated cells (cells expressing WT and mutant transgenes in the absence of endogenous CTCF). The uncropped pictures of the WB gels are shown below. This figure is related to **Figure 3**.

**Figure S14:**
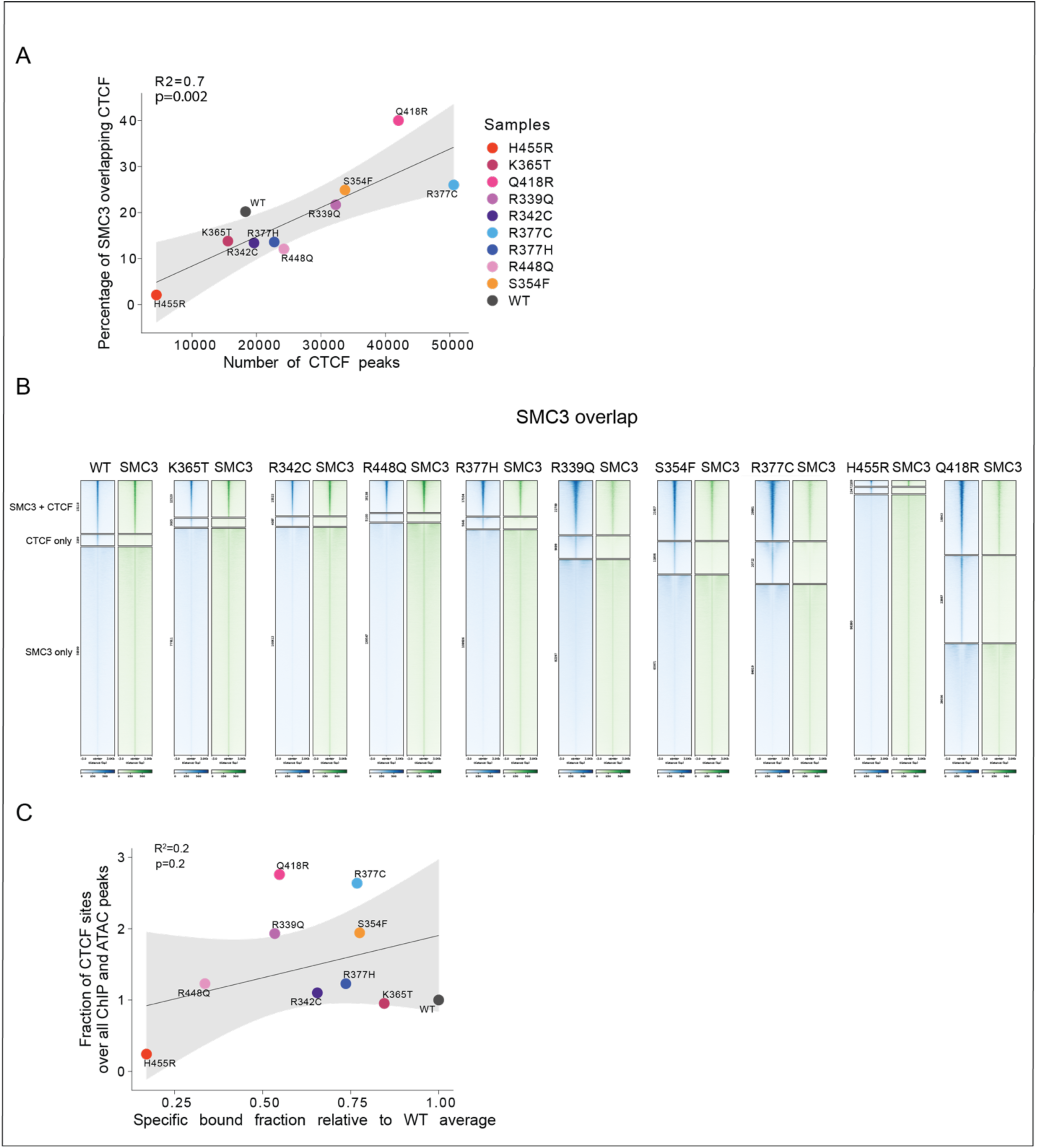
The percentage of SMC3 overlapping CTCF and the overall fraction of bound CTCF sites are less informative. **(A)** Correlation between the percentage of SMC3 overlapping CTCF peaks and the number of CTCF peaks. (**B**) Heatmaps of CTCF (blue) and SMC3 (green) in CTCF WT and mutant expressing cells at CTCF only, CTCF and SMC3, and SMC3 only sites. (**C**) Non-significant correlation between the chromatin bound mutant versus WT fraction detected by FRAP and the fraction of all CTCF binding sites relative to all FLAG and SMC3 ChIP-seq and ATAC-seq peaks. This figure is related to **Figure 3**.

**Figure S15:**
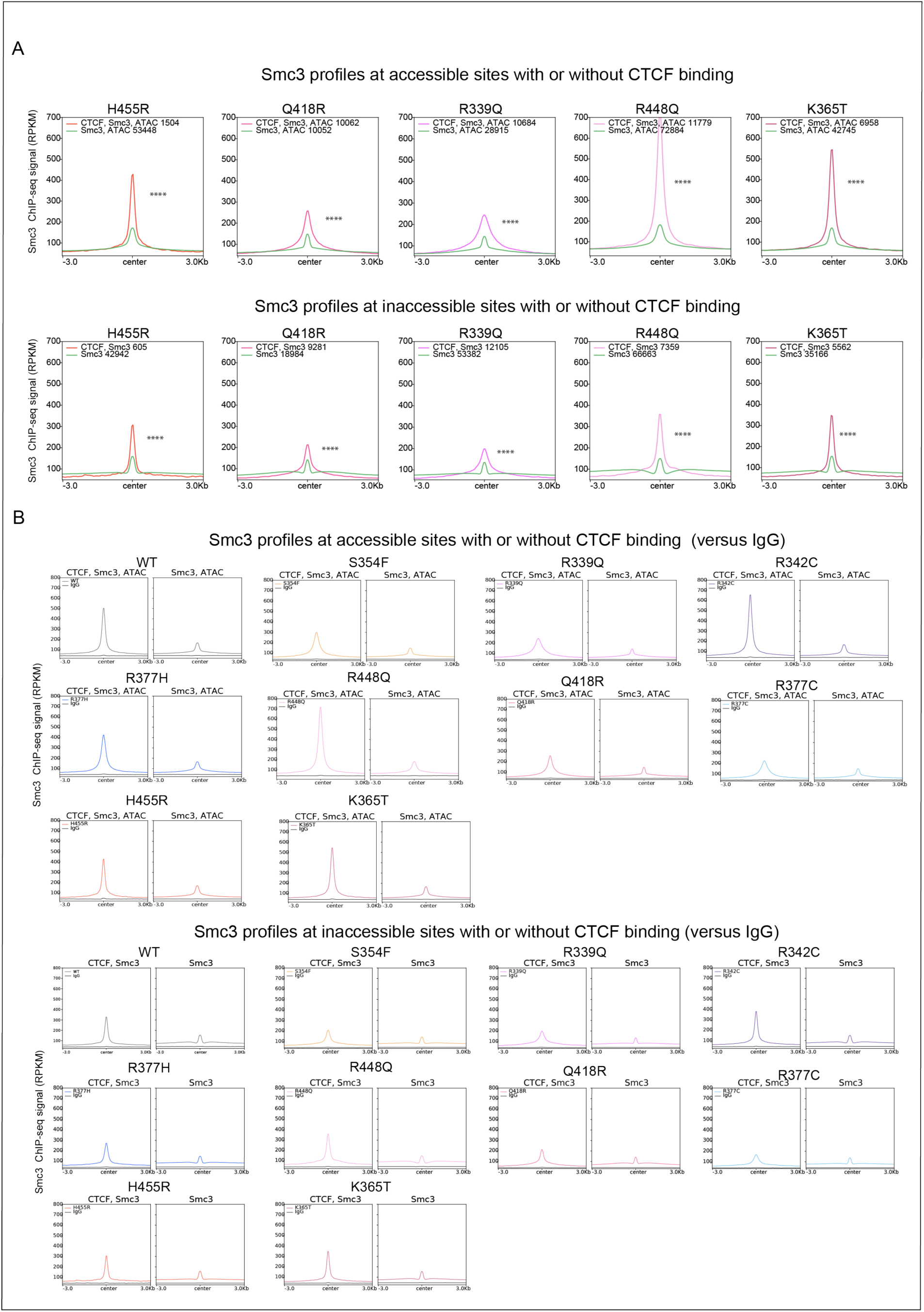
The relationship between CTCF binding, cohesin overlap and accessibility. **(A)** Profiles of SMC3 binding in WT and mutant CTCF expressing cells is shown for accessible (top) and inaccessible (bottom) sites with (color-coded for each mutant) or without (green) CTCF binding. The remaining mutants are shown in **Figure 4C**. Wilcoxon p-values were coded as follow: * 5×10^-2^-5×10^-3^, ** 5×10^-3^-5×10^-4^, *** 5×10^-4^-5×10^-5^, **** <5×10^-5^. (**B**) Profiles of SMC3 binding in WT and mutant CTCF expressing cells compared to IgG at accessible (top) and inaccessible (bottom) sites with or without CTCF (right and left profiles, respectively). Wilcoxon p-values were coded as follow: * 5×10^-2^-5×10^-3^, ** 5×10^-3^-5×10^-4^, *** 5×10^-4^-5×10^-5^, **** <5×10^-5^. This figure relates to **Figure 4**.

**Figure S16:**
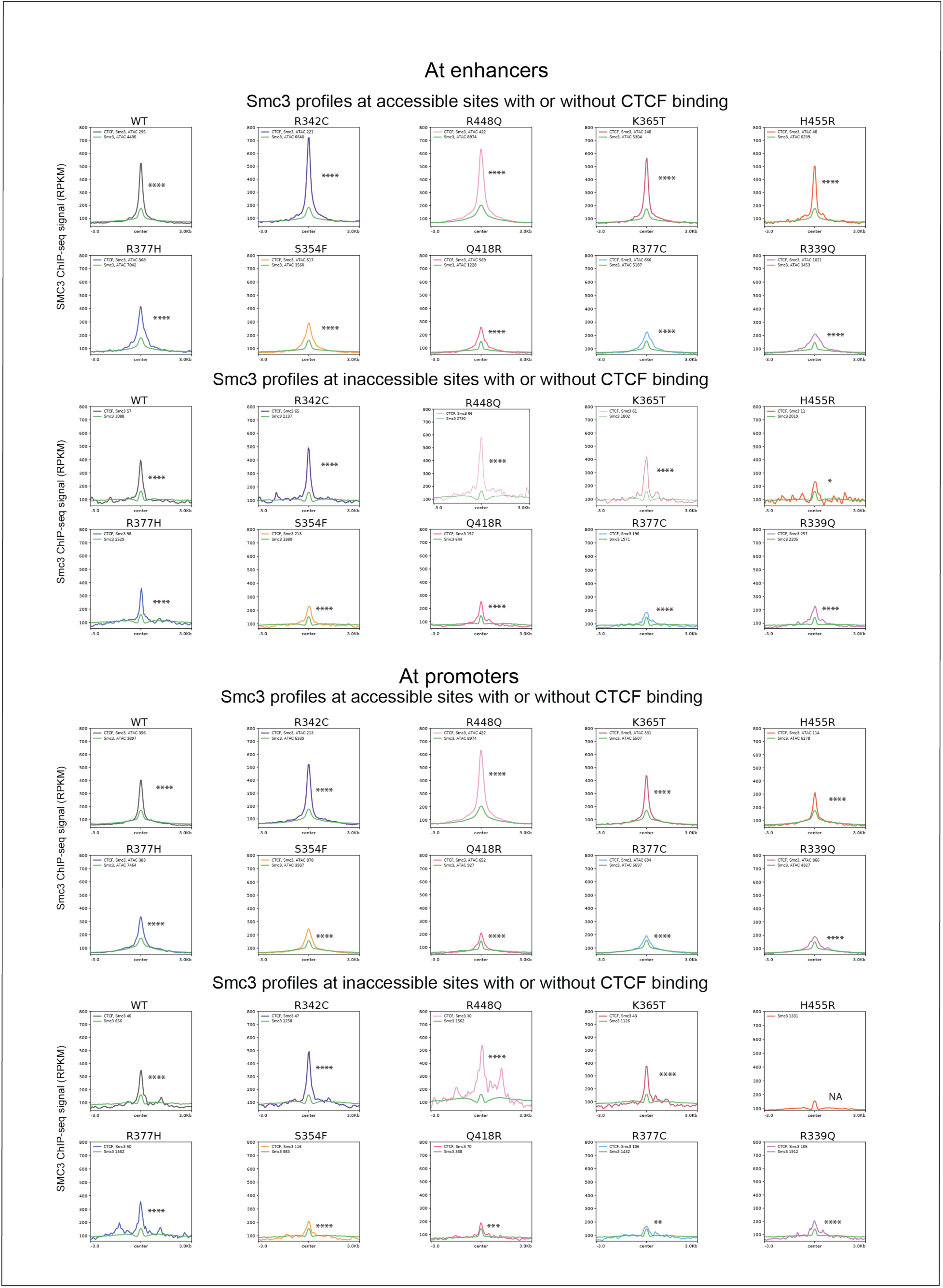
The relationship between CTCF binding, cohesin overlap and accessibility at enhancers and promoters. Profiles of SMC3 binding in WT and mutant CTCF expressing cells is shown for accessible and inaccessible CTCF binding at enhancers (top) and gene promoters (bottom) with (color-coded for each mutant) or without (green). Wilcoxon p-values were coded as follow: * 5×10^-2^-5×10^-3^, ** 5×10^-3^-5×10^-4^, *** 5×10^-4^-5×10^-5^, **** <5×10^-5^. This figure relates to **Figure 4**.

**Figure S17:**
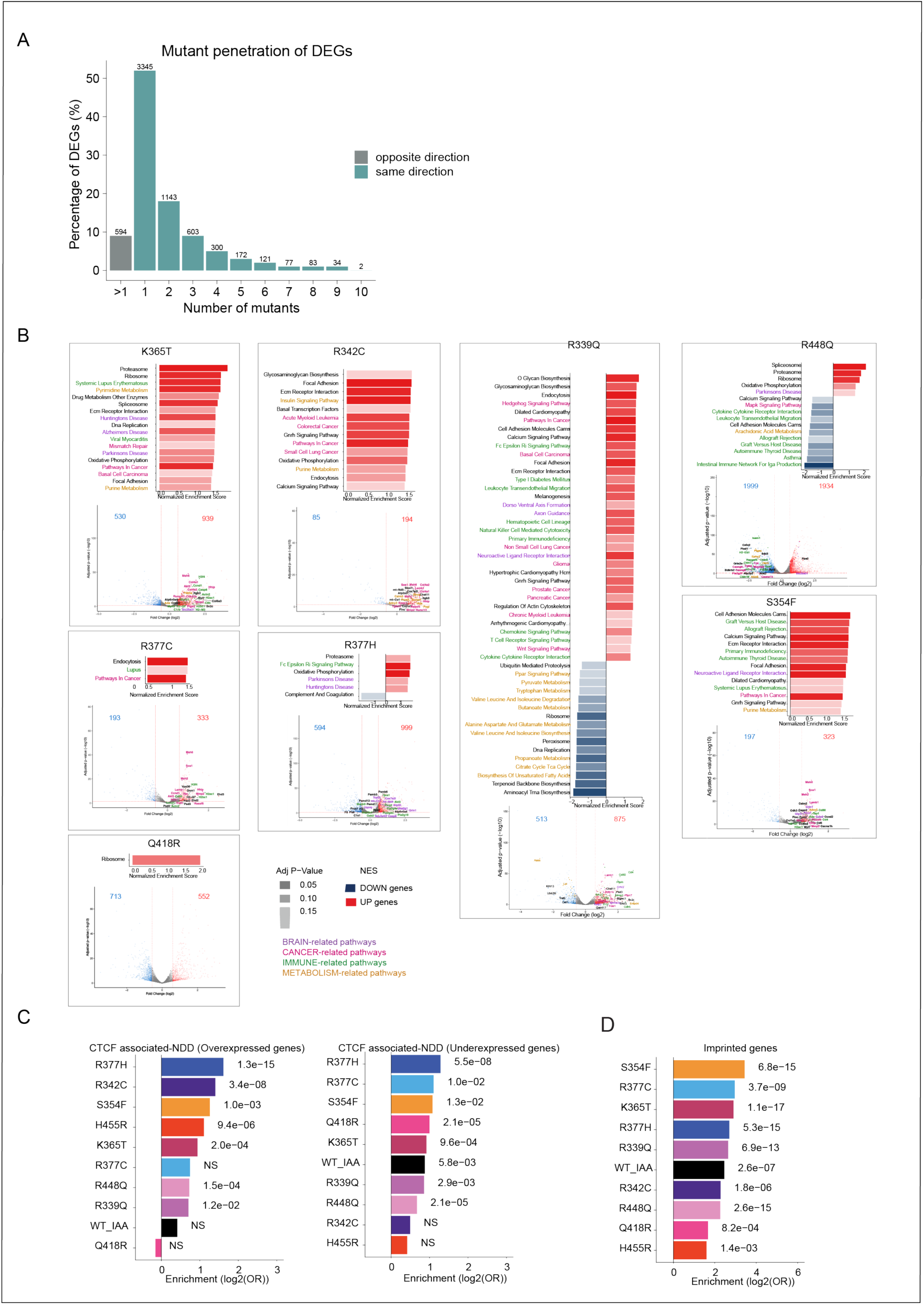
CTCF mutations alter gene expression. (**A**) Bar graph showing the percentage of DEGs found in 1 to 10 mutants, including IAA. The list of genes is reported in **Table S2**. (**B**) Gene set enrichment analysis of DEGs. All other mutants are shown in **Figure 5B**. For each condition, bar graphs show significantly enriched KEGG pathways. The volcano plots below highlight the DEGs belonging to these enriched pathways with genes outside of these pathways shown in black. Brain related pathways are shown in purple, cancer in pink, immune in green and metabolism in mustard. (**C**) Enrichment of the DEGs in over- and under-expressed genes reported in CTCF associated human neuro-developmental disorders [S4]. (**D**) Enrichment of the DEGs in imprinted genes reported in Geneimprint (https://www.geneimprint.com). This figure relates to **Figure 5**.

**Figure S18:**
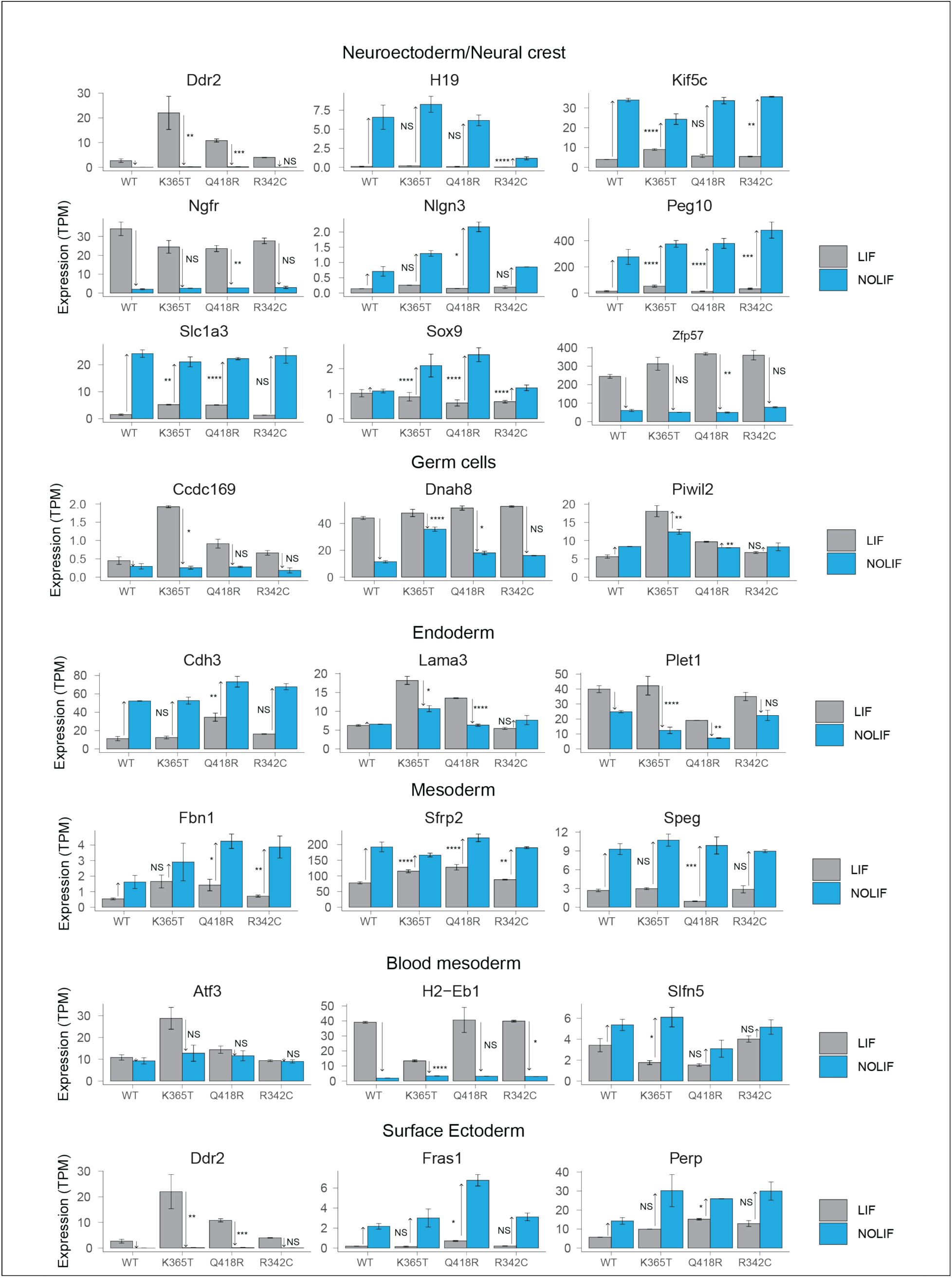
Examples of genes implicated in altered differentiation trajectories in WT versus CTCF mutant expressing cells. Bar graphs showing the comparison of WT versus mutant CTCF expression levels (TPM) in LIF and no LIF medium. The p-values assessing the significance of the longitudinal differences between mutant and WT were computed using linear regressions at the gene level. p-values are coded as followed *: 5×10^-2^-5×10^-3^, **5×10^-3^-5×10^-4^, *** 5×10^-4^-5×10^-5^, ****<5×10^-5^. This figure relates to **Figure 5**.

**Figure S19:**
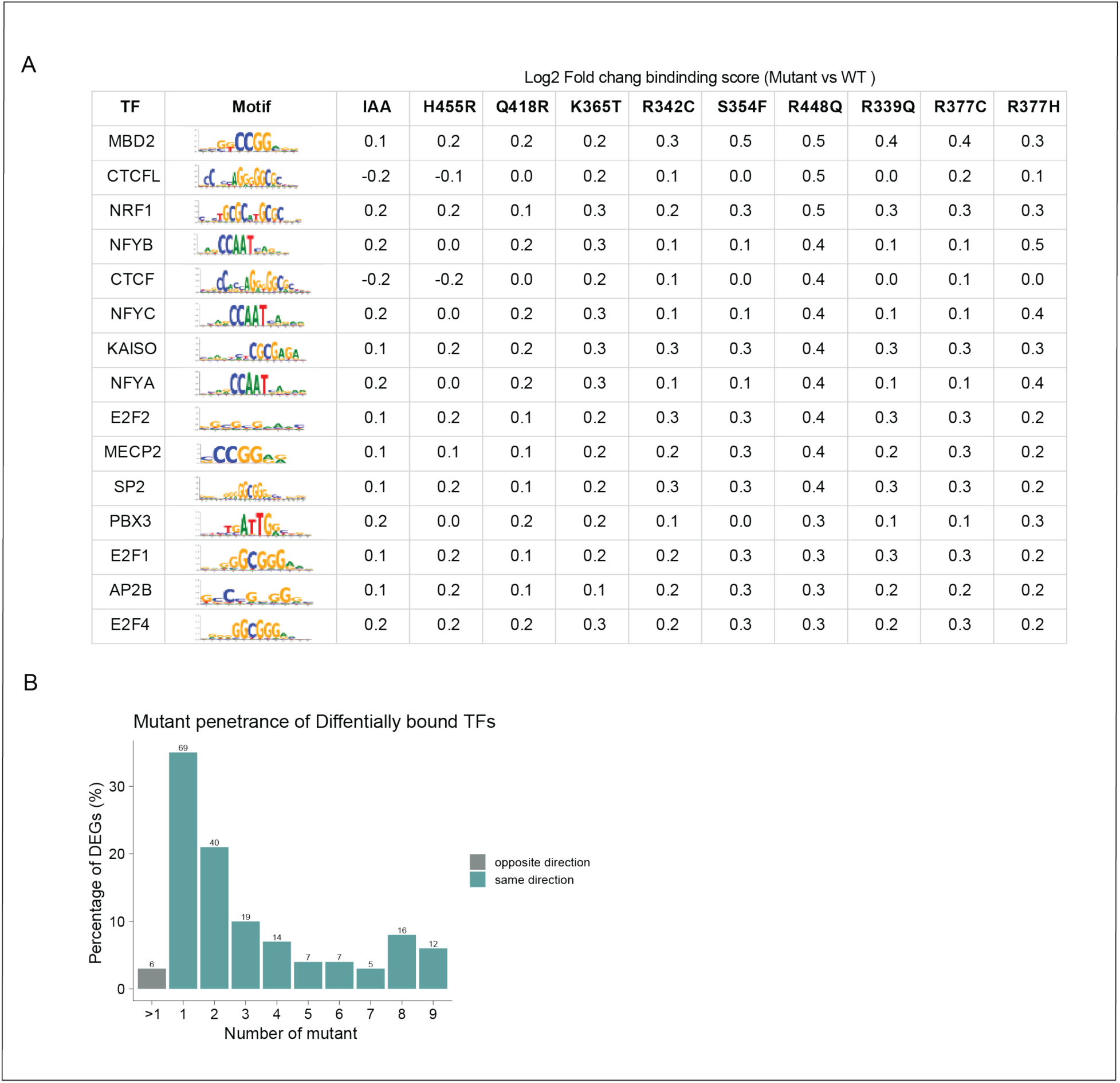
CTCF mutations alter TF binding. **(A)** Top 15 predicted differentially bound TFs. The TFs were sorted by the maximum absolute log2 fold change of binding score between mutant versus WT observed across mutants. The total number of sites and p-value for these TFs, as well as the full list of differentially bound TFs for each mutant are reported in **Table S4A**. (**B**) Bar graph showing the percentage of differentially bound TFs found in 1 to 9 mutants including IAA (none were found in all the conditions). The list of significant differentially bound TFs is reported in **Table S4A**. This figure relates to **Figure 5**.

**Figure S20:**
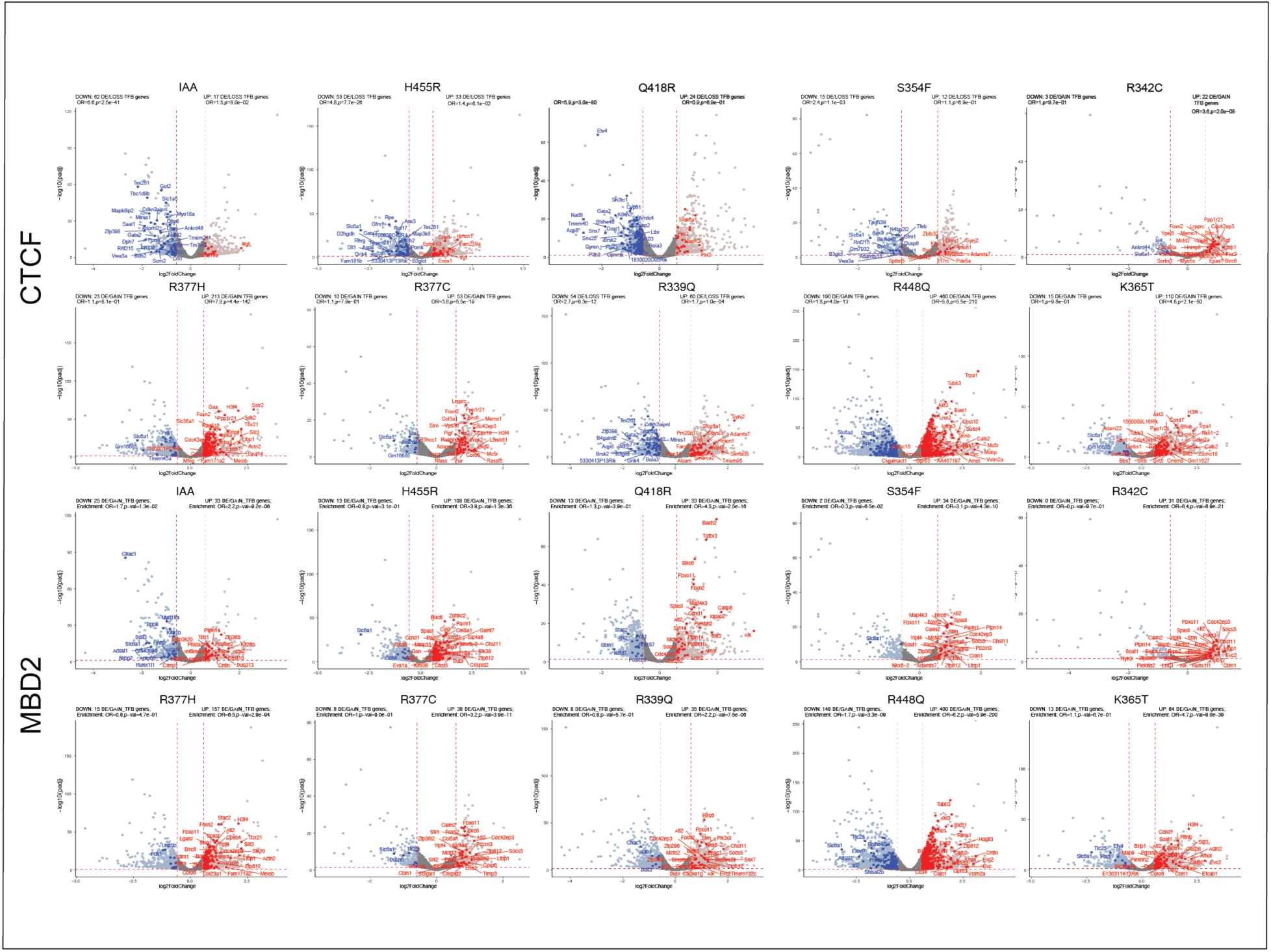
The DEGs are enriched in the target genes of differentially bound TF. Volcano plots show examples of two differentially bound TFS (CTCF, MBD2) overlapping the promoters of DEGs in CTCF mutant expressing cells (ID). The remaining mutant comparisons are shown in **Figure 5E**. The enrichment of the TF target genes among the up- and down-regulated genes are reported on top on the volcanos (Odds Ratios (ORs), and p-values). Enrichment for all the differentially bound TFs and their differentially expressed target genes are reported in **Table S4B** and **S4C**. This figure relates to **Figure 5**.

**Figure S21:**
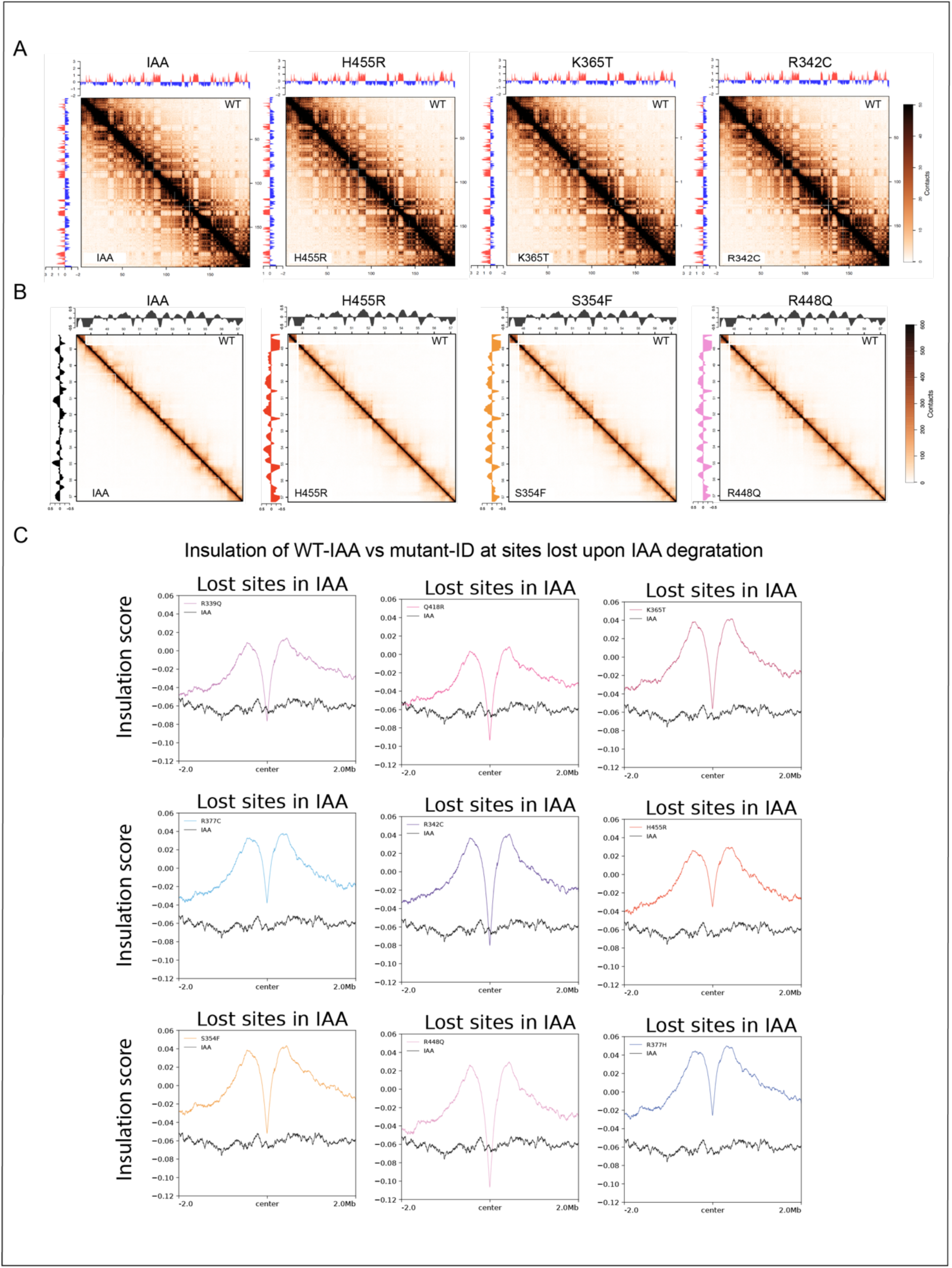
CTCF mutations alter insulation and chromatin interactivity. (**A**) Absolute Hi-C matrices of chromosome 1 using a 100kb resolution for IAA and 3 mutants (bottom left) versus WT (top right) showing the minimal changes in chromatin compartmentalization. The compartment scores are shown on the left for the mutant and top for WT. (**B**) Absolute Hi-C matrix for IAA and the mutant shown in Figure **6E** compared to WT. The insulation scores are shown on the left for the mutant and top for WT. (**C**) Profiles comparing the insulation score in WT-IAA versus mutant-ID at lost CTCF sites upon IAA degradation. This figure relates to **Figure 6**.

**Figure S22:**
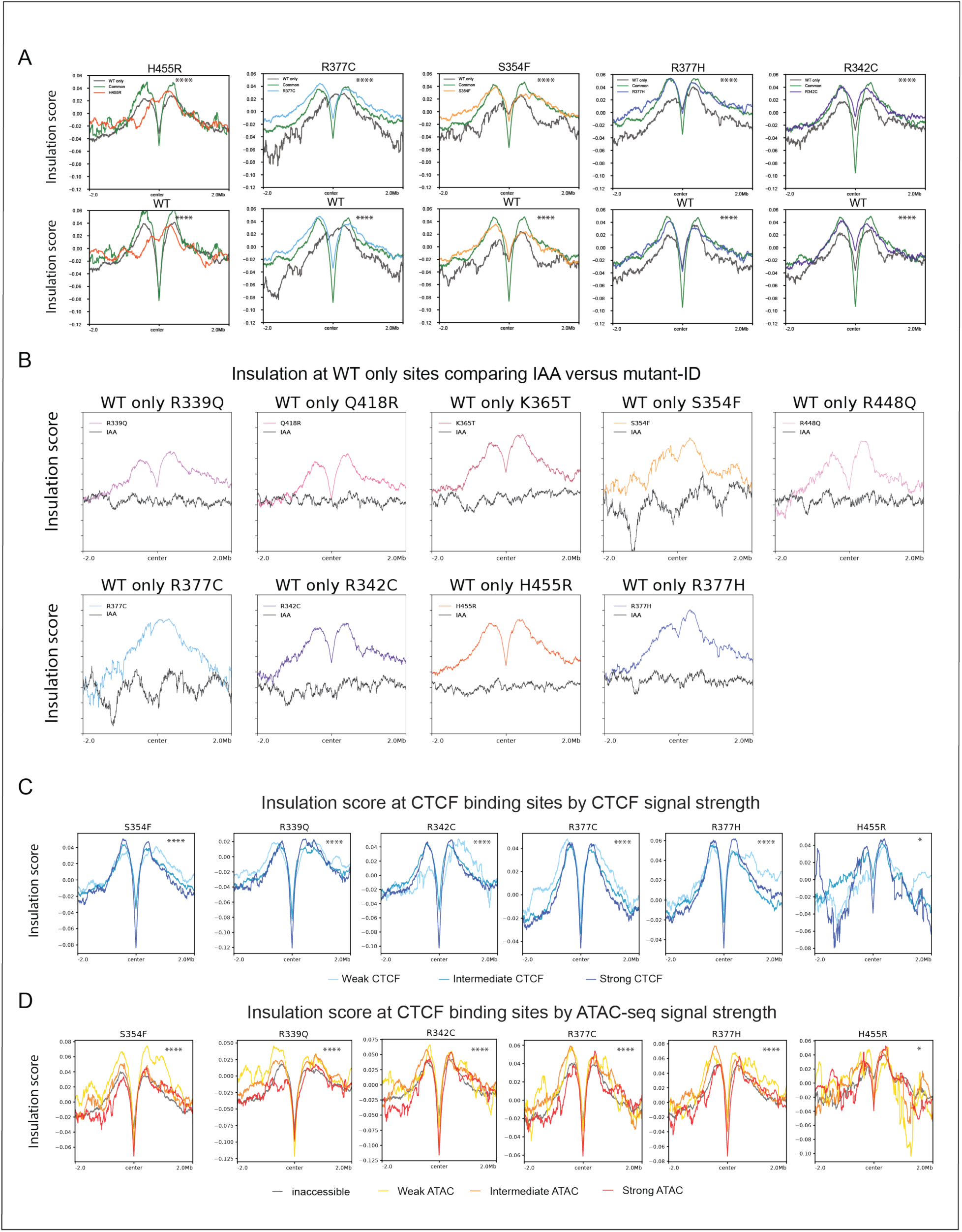
Insulation is a function of CTCF and ATAC-seq signal strength. (**A**) Profiles show the aggregated insulation score at WT only, common and mutant only binding sites for four mutants. The profiles for the remaining mutants are shown in **Figure 6F**. (**B**) Profiles comparing the insulation score in mutant-ID versus IAA conditions at WT only sites. (**C**) Profiles show the aggregated insulation score at CTCF binding sites with weak, intermediate and strong CTCF signals for different mutants. The profiles for the remaining mutants are shown in **Figure 6G**. (**D**) Profiles show the aggregated insulation score at CTCF binding sites with no, weak, intermediate and strong ATAC signals for different mutants. The profiles for the remaining mutants are shown in **Figure 6H**. For (**C**) and (**D**), Kruskal-Wallis p-values were coded as follow: * 5×10^-2^-5×10^-3^, ** 5×10^-3^-5×10^-4^, *** 5×10^-4^-5×10^-5^, **** <5×10^-5^. This figure relates to **Figure 6**.

**Figure S23:**
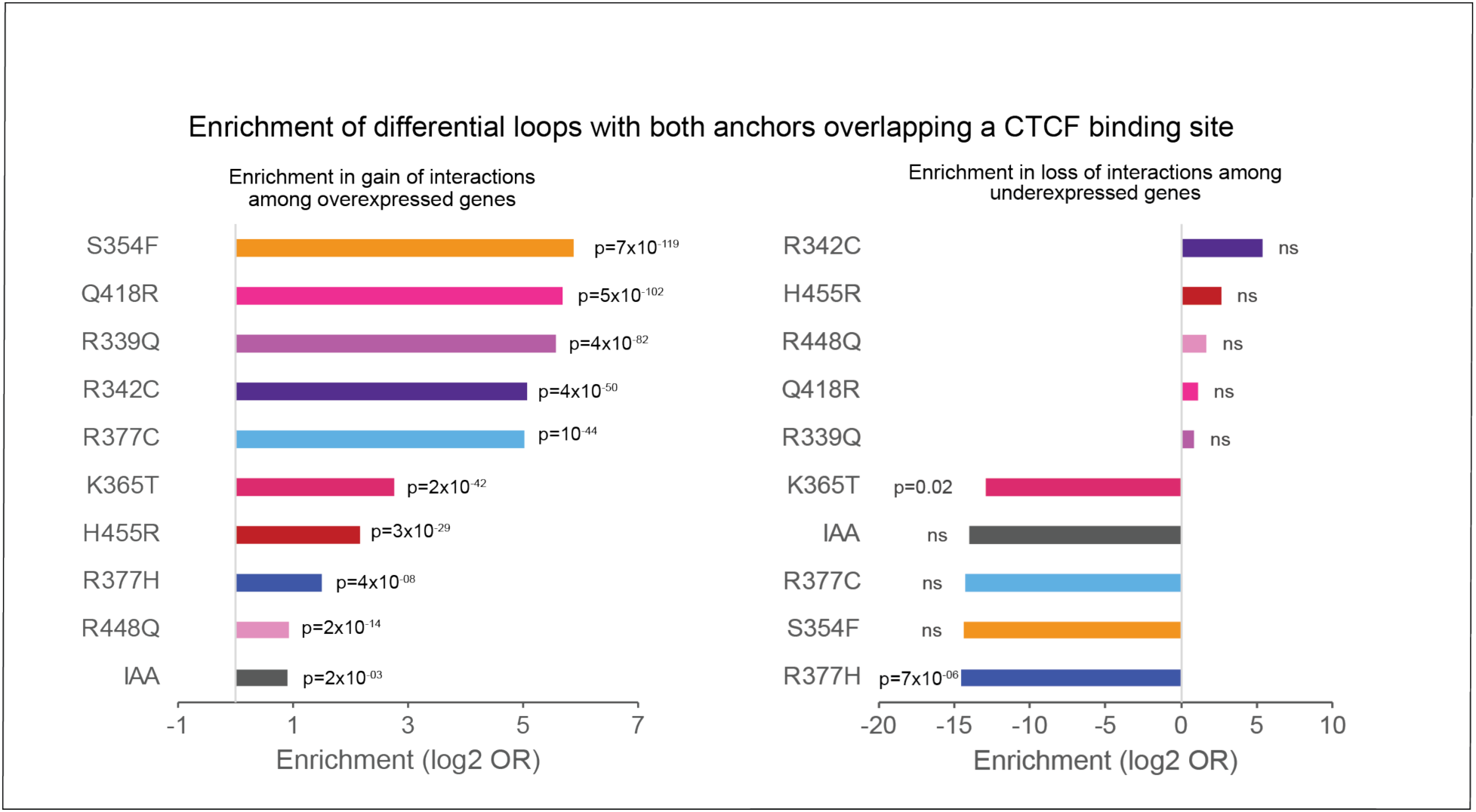
Differentially expressed genes are enriched in gain of loops with a CTCF binding site at both anchors. Bar graphs showing the enrichment (calculated over WT ID) of over-expressed (top) or under-expressed (bottom) genes in gained (top) or lost (bottom) loops with both anchors overlapping a CTCF binding sites and at least one anchor overlapping the promoter of DEGs in IAA and CTCF mutants. This figure relates to **Figure 7**.

**Figure S24:**
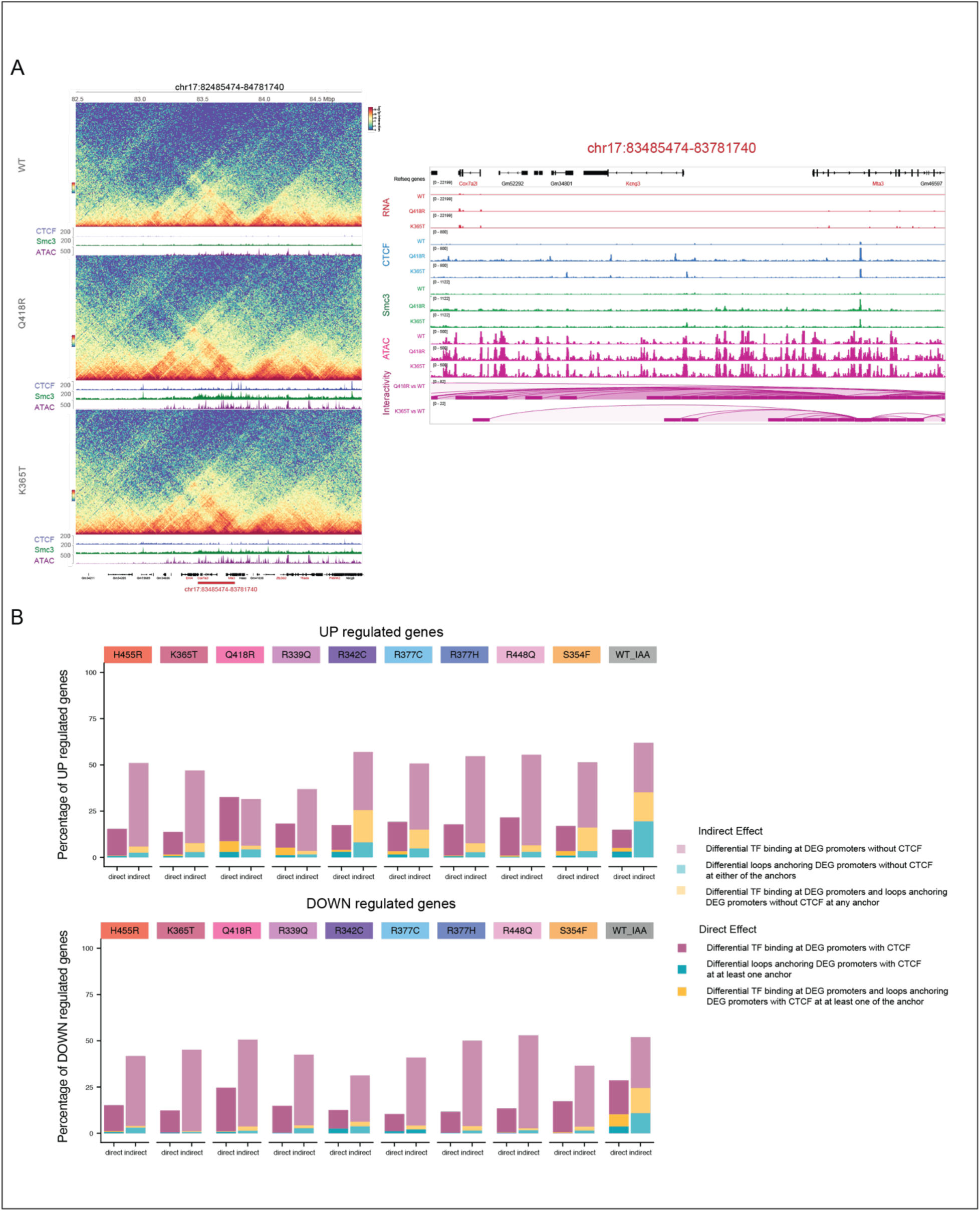
Indirect effects resulting from changes in TF binding at gene promoters account for the majority of CTCF mutation-mediated changes in expression. (**A**) Example of a locus on chromosome 7 (red rectangles) with a direct effect of gain in CTCF binding and chromatin interactivity. The left panel shows the Hi-C interaction matrices in WT, Q418R and K365T with both gain of intra- and inter-TAD interactions in the mutant compared to WT. The left panels show the zoom-in tracks of CTCF (blue) SMC3 (green), ATAC (red) and significant differential chromatin loops (purple) with one anchor overlapping the differential CTCF binding sites. Overexpressed genes are highlighted in red. Of note, only gained interactions were detected at these loci. This analysis was performed at 10 kb resolution. (**B**) Bar graphs showing the percentage of UP and DOWN DEGs associated with differential TF/CTCF binding, differential loops or both, stratified by direct and indirect effect. A direct effect was defined as predicted differential CTCF binding at the gene promoter and/or differential loops anchoring the DEG promoters with a CTCF peak at either anchor. This figure relates to **Figure 7**.

